# Cortical Organoids Model Early Brain Development Disrupted by 16p11.2 Copy Number Variants in Autism

**DOI:** 10.1101/2020.06.25.172262

**Authors:** Jorge Urresti, Pan Zhang, Patricia Moran-Losada, Nam-Kyung Yu, Priscilla D. Negraes, Cleber A. Trujillo, Danny Antaki, Megha Amar, Kevin Chau, Akula Bala Pramod, Jolene Diedrich, Leon Tejwani, Sarah Romero, Jonathan Sebat, John R. Yates, Alysson R. Muotri, Lilia M. Iakoucheva

## Abstract

Reciprocal deletion and duplication of 16p11.2 region is the most common copy number variation (CNV) associated with Autism Spectrum Disorders. We generated cortical organoids from skin fibroblasts of patients with 16p11.2 CNV to investigate impacted neurodevelopmental processes. We show that organoid size recapitulates macrocephaly and microcephaly phenotypes observed in the patients with 16p11.2 deletions and duplications. The CNV has mirror-opposite effect on neuronal maturation, proliferation, and synapse number, in concordance with its effect on brain growth in humans. We demonstrate that 16p11.2 CNV alters the ratio of neurons to neural progenitors in organoids during early neurogenesis, with excess of neurons and depletion of neural progenitors observed in deletions, and mirror phenotypes in duplications. Transcriptomic and proteomic profiling revealed multiple dysregulated pathways, including defects in neuron migration. Inhibition of activity of the small GTPase RhoA rescued migration deficits. This study provides insights into potential neurobiological mechanisms behind the 16p11.2 CNV during neocortical development.

## Introduction

Over the last decade it has been convincingly demonstrated that deletions and duplications of large genomic regions, or *copy number variants* (CNVs), are associated with multiple neurodevelopmental disorders ^1–4^. The deletions (DEL) of a genomic region spanning 29 genes on human chromosome 16, 16p11.2 CNV, had been identified as one of the strongest risk factors for Autism Spectrum Disorder (ASD) and Intellectual Disability (ID), whereas the duplications (DUP) of the same region were associated with ASD, ID, schizophrenia (SCZ) and Bipolar Disorder (BD) ^2, 3, 5–7^. Most importantly, DEL and DUP were associated with macrocephaly and microcephaly in human carriers, respectively ^8, 9^. This phenotype, however, had not been recapitulated in mouse models ^10–12^. Some of the animal studies have reported mirror effect of 16p11.2 CNV on regional brain volume along with changes in brain cytoarchitecture, behavior and viability, but there was little direct concordance in the phenotypes between human and mouse models. For example, at least one of these models observed phenotypes opposite to humans: DEL 16p11.2 mice were smaller and lean, whereas DUP 16p11.2 mice were larger and obese ^12^.

Significant progress has been made for implicating various biological mechanisms that may be impacted by the 16p11.2 CNV. RNA sequencing of cortex from 16p11.2 deletion and duplication mice identified altered expression of genes and networks that converged on general ASD-associated pathways including synaptic function, chromatin modification and transcriptional regulation ^13^. Transcriptome profiling of lymphoblastoid cell lines of 16p11.2 CNV human carriers identified expression dysregulation of neuronal-related gene in deletion, but not in duplication ^14^. Dysregulation of ciliopathy genes ^15^, ERK/MAPK signaling ^16, 17^, and metabotropic glutamate receptor 5 (mGluR5)-dependent synaptic plasticity and protein synthesis ^18^ in mouse models were all implicated. Despite the progress made with regard to understanding of the general mechanisms disrupted by the 16p11.2 CNV in animal models and non-neuronal human cells, the question of how 16p11.2 variants impact early human brain development remained unanswered.

Recent advances in stem cell technologies opened a window of opportunities for investigating brain disorders using human-based *in vitro* systems ^19^. During the last decade, 3D cortical organoid models have been successfully developed ^20^. Characterization of these models demonstrated that they closely resemble human fetal brain, forming structures reminiscent of deeper cortical layers and sharing cell types and transcriptomic signatures ^21–24^. The 3D organoid models have proven advantages over 2D models for investigating human brain diseases ^25–27^. Several studies had used 3D cortical organoids to model lissencephaly ^28, 29^, non-syndromic autism ^30^, autosomal recessive primary microcephaly ^20^, and Timothy syndrome ^31^. Here, we used cortical organoids to perform 3D modeling of fetal brain development of the most common autism subtype associated with deletions and duplications of the 16p11.2 CNV.

In this study, we generated induced pluripotent stem cells (iPSCs) and cortical organoids from 16p11.2 DEL and DUP patient fibroblasts and unrelated healthy control (CTRL) individuals, and investigated molecular and cellular processes that are disrupted by this genetic variant (**Fig. 1A**). We found that the size of deletion organoids is larger, and duplication organoids is smaller, recapitulating mirror effect of 16p11.2 CNV on brain size in humans. Transcriptomic and proteomic profiling of organoids identified differentially expressed genes, proteins, and co-expression modules impacted by the 16p11.2 CNV. The results were validated by a panel of orthogonal assays. Cellular assays confirmed that 16p11.2 deletions and duplications exhibit defects in neuronal maturation, migration, morphology and synaptic abnormalities, implicating disrupted neurogenesis. These mechanisms have not been previously associated with 16p11.2-linked autism, likely due to lack of appropriate 3D models of human fetal brain. We identified multiple pathways disrupted by the 16p11.2 CNV, and found activation of RhoA signaling to be a likely contributor to defects in migration. Our study makes significant contribution to understanding of neurobiological mechanisms that may be disrupted during early human neocortical development in the 16p11.2 CNV carriers, and offers potential path for therapeutic interventions.

**Figure 1.**
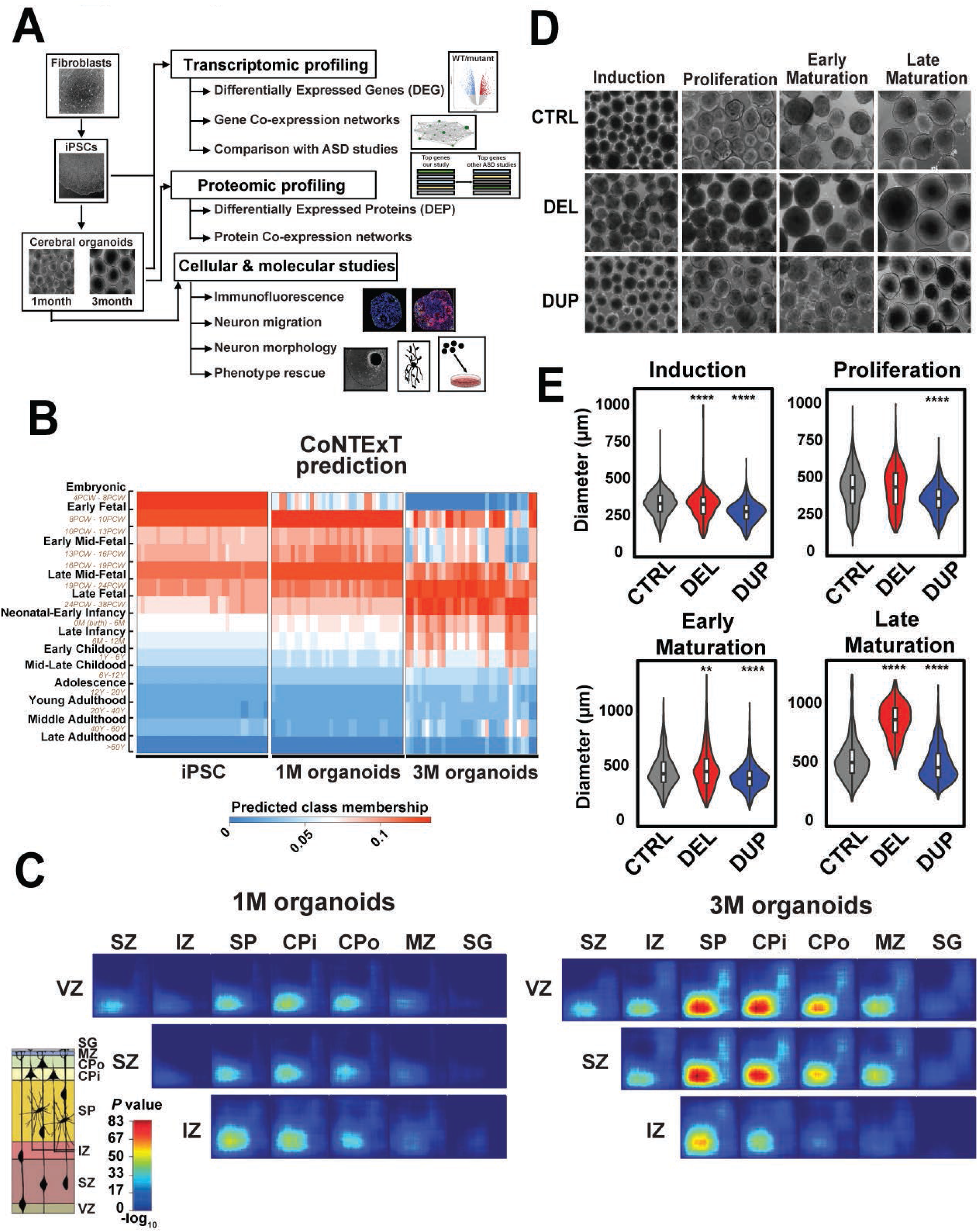
Generation and characterization of cortical organoids from 16p11.2 patient and control iPSCs. (A) Overview of Study Design and Analyses. (B) Comparison of iPSCs and organoids transcriptomes with the human developing brain transcriptome using CoNTExT ^36^. The samples from 3 individuals of each genotype (CTRL, DEL, DUP), with 2 clones per individual and 2 replicates per clone (n=36 datasets) are shown for each group (iPSC, 1M and 3M organoids). PCW – post conception weeks, M – months, Y-years. (C) Predicted laminar transitions for 1M and 3M organoids using TMAP ^36^ and the transcriptome of laser capture microdissected cortical laminae from postmortem human fetal brain (15–21 PCW) ^37^. Rank-rank hypergeometric overlap (RRHO) maps for CTRL organoids (n=12 datasets) from 3 patients, 2 clones per patient, 2 replicates per clone are shown, with CTRL iPSCs (n=12 datasets) used as a second time point. Each pixel represents the overlap between fetal brain and organoids transcriptome, color-coded according to the −log_10_ p-value of a hypergeometric test. On each map, the extent of shared upregulated genes is displayed in the bottom left corner, whereas shared downregulated genes are displayed in the top right corners. (D) Representative images of cortical organoids for each genotype (CTRL, DEL, DUP) at different time points of differentiation: Induction (6 days of differentiation), Proliferation (16 days of differentiation), Early maturation (1M of differentiation) and Late maturation (3M of differentiation). (E) Analysis of size differences between cortical organoids of each genotype (CTRL, DEL and DUP) at different time points of differentiation. Organoids diameter (*n*>100 for each genotype) was measured using ImageJ, and average diameter for each genotype was computed. P-values were calculated using one-way ANOVA. Stars above DEL and DUP represent comparison to CTRLs. \*\*\*\**p*<0.0001, \*\*\**p*<0.001, \*\**p*<0.01.

## Results

### Cortical organoids maturation resembles stages of human brain development

To investigate how 16p11.2 CNV impacts early stages of human brain development, and what molecular pathways are dysregulated by this genetic variant, we generated cortical organoids from the 16p11.2 CNV carriers. We first obtained iPSCs by reprogramming patient- and control-derived fibroblasts using episomal transduction, and then differentiated iPSCs into cortical organoids as previously described ^32^.

We selected six male 16p11.2 CNV carriers with extreme head size phenotypes (age-normalized head circumference Z score range from 2.51 to 4.32 in DELs; and from −0.69 to − 1.63 in DUPs) for this study. We decided to focus this phenotype, because previous studies from our and other laboratories using infection with Zika virus were able to successfully recapitulate microcephaly in organoid models ^33, 34^. The restriction to only male gender was due to samples availability. The details of patients’ phenotypes are described in **Table S1**. Three gender-matched healthy unrelated individuals that did not carry 16p11.2 CNV were used as controls. We performed rigorous quality control assessment of reprogrammed iPSCs clones by comparing them to parental fibroblasts using immunofluorescence (**figs. S1-S2**), qRT-PCR and SNP array genotyping (**figs. S3-S4**). After confirming the presence of 16p11.2 CNV in patient samples and making sure that no additional CNVs were introduced by reprogramming, we selected two clones per individual for organoids production. We performed bulk RNA sequencing (RNA-seq) of a total of 108 samples derived from iPSCs, 1 month-old (1M) and 3 month-old (3M) organoids (36 samples at each time point). We sequenced two clones per individual, and two replicates per clone for all three genotypes (DELs, DUPs and CTRLs) (**fig. S5**). RNA sequencing quality control parameters are shown in **Table S2**.

To investigate whether developmental maturity and laminar organization of our organoids resembles human brain, we compared transcriptional profiles of iPSCs and organoids with the atlas of the developing human brain ^35^ using CoNTExT approach ^36^. Transcriptional profiles of iPSCs from all individuals closely matched those of embryonic (4-8 PCW) and early fetal (8-10 PCW) human brain, independently validating successful conversion of fibroblasts into pluripotent state by reprogramming (**Fig. 1B**). Transcriptional profiles of 1 month-old organoids resembled those of early mid-fetal (13-16 PCW) through late mid-fetal (19-24 PCW) periods. Likewise, transcriptional profiles of 3 month-old organoids mostly recapitulated those of late mid-fetal (19-24 PCW) through neonatal-early infancy (birth to 6 months) developmental periods, and even further into late infancy (6-12 M).

Next, we examined the degree of overlap between *in vivo* cortical development of prenatal human brain and our *in vitro* differentiated organoids using TMAP ^36^. We compared transcriptional profiles of our organoids with those derived from laser capture microdissected cortical laminae of postmortem human fetal brain (15-21 PCW) ^37^. TMAP performs serialized differential expression analysis between any two *in vivo* developmental periods and any two *in vitro* differentiation time points, followed by quantification of overlap ^36^. Laminar matching by TMAP demonstrated transitions between proliferative layers (ventricular VZ, subventricular SZ and intermediate IZ zones) and post mitotic upper layers for both, 1M and 3M old organoids (**Fig. 1C**). We observed that in 3M organoids laminar transition into these upper layers manifested greater shift than in 1M organoids. For example, greater correspondence to upper layers (subplate SP, cortical plate inner CPi and outer CPo layers and marginal zone MZ) was visible at 3M compared to 1M, consistent with increased maturity at 3M. We replicated this maturation shift using an additional independent dataset from fetal human brain ^38^ (**fig. S6**). Together, the results suggest that cortical organoids from DEL, DUP and CTRL individuals mature over time, closely recapitulating human brain development in terms of temporal transitions and laminar organization. Furthermore, organoids between 1M and 3M of differentiation most closely resemble human mid-fetal brain development, and represent suitable models for studying molecular basis of neurodevelopmental disorders, considering a proven role of this period in ASD and SCZ pathogenesis ^39–41^. These results are in agreement with a previous study that concluded that brain organoids faithfully recapitulate fetal development at the transcriptional level ^42^.

### Patient-derived organoids recapitulate macrocephaly and microcephaly phenotypes

Since our patients with 16p11.2 DELs and DUPs were selected based on the extreme head circumference phenotypes, we investigated whether organoids recapitulate these phenotypes. We measured the diameter of 16p11.2 and control organoids at four time points, at day 6 (D6, neural induction), day 16 (D16, proliferation), 1 month (1M, early maturation) and 3 months (3M, late maturation). We observed significant size differences between CTRL and DEL or DUP organoids at almost all time points, with DEL organoids being larger and DUP organoids being smaller than CTRL (**Fig. 1D-E** and **Table S3**). These differences were most apparent at the late maturation stage, corresponding to 3M of differentiation. These results demonstrate that cortical organoids recapitulate 16p11.2 patients’ brain size phenotypes.

### Differential gene expression analysis points to dysregulation of multiple pathways by 16p11.2 CNV

To understand molecular pathways dysregulated by the 16p11.2 CNV, we performed differential gene expression analyses of 108 transcriptomes derived from iPSCs, 1M, and 3M organoids (**fig. S5 and STAR Methods**). An extensive quality control and normalization included sample outlier detection, principal component analyses, surrogate variable analysis, and covariates selection with multivariate adaptive regression splines (**figs. S7-S8 and STAR Methods**). For gene differential expression analyses, we implemented limma-voom model with ‘duplicateCorrelation’ function to account for duplicate samples (clones and replicas) from the same individuals, and to avoid pseudo-replication in the analyses ^43^.

We identified 1044, 118 and 430 differentially expressed genes (DEGs) in DELs *vs* DUPs at 10% FDR for iPSCs, 1M and 3M organoids, respectively (**Fig. 2A-C and Table S4**). Genes from the 16p11.2 *locus* were most significantly dysregulated, confirming the expected *cis*-effect of CNV on gene expression. In addition, 16p11.2 CNV had a significant effect on the expression of many genes outside of the *locus*. Gene Ontology (GO) analyses of DEGs from iPSC revealed enrichment in cell migration and motility, actin filament-based movement and potassium ion homeostasis; whereas DEGs from organoids were enriched in regulation of neuron migration, actin cytoskeleton-related functions, layer formation in cerebral cortex and neuron differentiation (**Fig. 2A-C** and **Table S5**). The enrichment of migration and motility-related functions were consistently observed across all three time points (iPSCs, 1M and 3M organoids), whereas neurogenesis and synaptic functions were observed only in organoids. This underscores the differences between iPSCs and organoids, with the prevalence of neurons in organoids.

**Figure 2.**
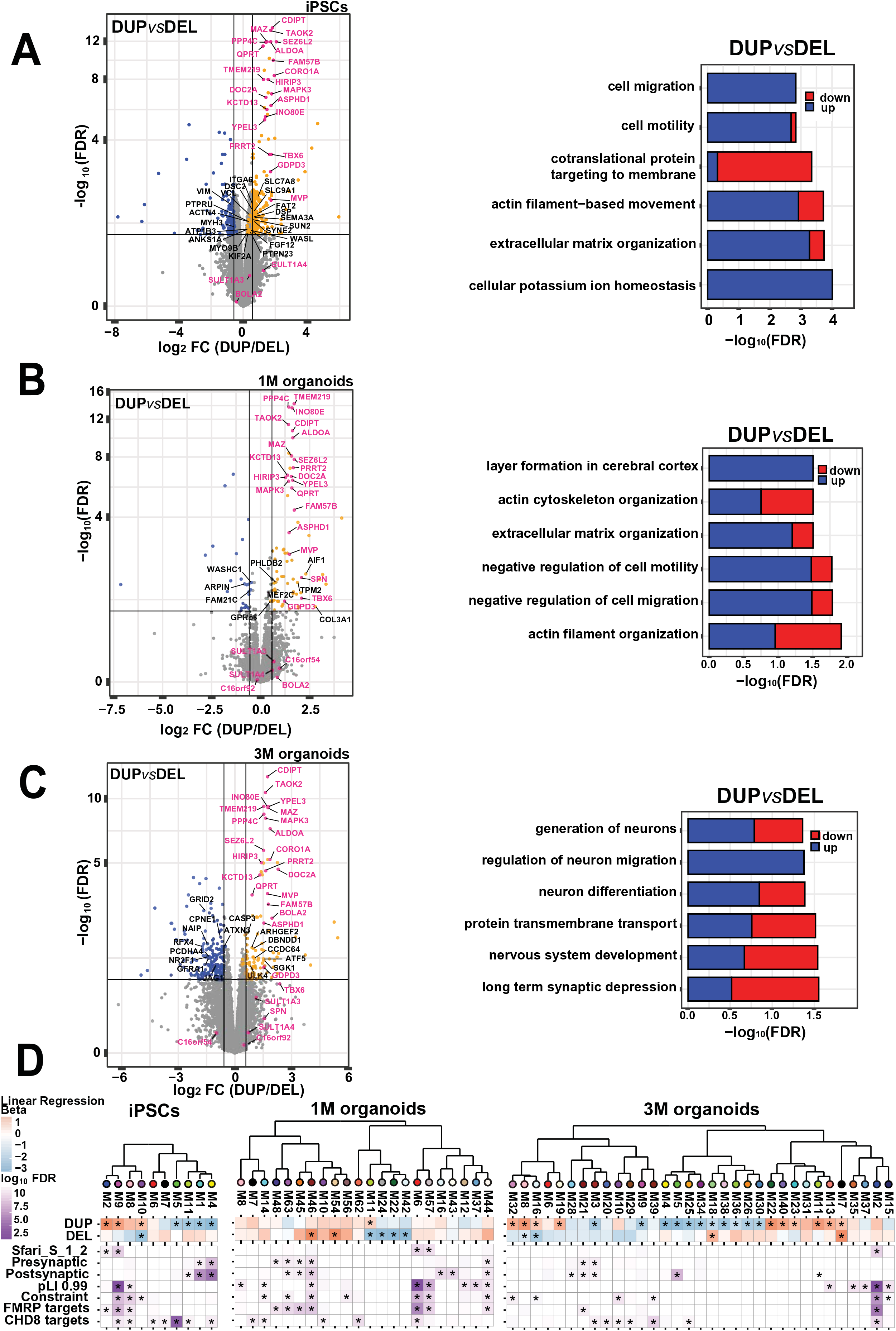
Differential gene expression and gene co-expression analyses of iPSCs, 1M and 3M cortical organoids. (A-C) Volcano plots of differentially expressed genes between DEL and DUP iPSCs, 1M and 3M organoids. Genes within 16p11.2 CNV are colored in pink. Genes colored in orange are upregulated in DUP and downregulated in DEL; genes colored in blue are downregulated in DUP and upregulated in DEL, in DUP *vs* DEL comparison. Gene Ontology enrichment analyses are shown as bar plots on the right. The contribution of up- or down-regulated genes to specific GO terms are shown in blue and red, respectively. (D) Hierarchical clustering of gene co-expression modules by module eigengene. Module-genotype associations (* FDR<0.1) are shown below each module. Module enrichment analyses against literature-curated gene lists with previous evidence for involvement in autism are shown at the bottom (* FDR<0.05). The lists include syndromic and highly ranked (1 and 2) genes from SFARI Gene database (https://gene.sfari.org/database/gene-scoring/); pre- and post-synaptic genes from SynaptomeDB ^87^; genes with probability of loss-of-function intolerance (pLI)>0.99 as reported by the Exome Aggregation Consortium ^88^; constrained genes ^89^; FMRP target genes ^90^, and CHD8 target genes ^91^.

### Gene co-expression analyses identifies migration and synaptic/neuronal modules

To further characterize signaling pathways and molecular processes dysregulated by the 16p11.2 CNV, we performed weighted gene co-expression network analysis (WGCNA) ^44^ and identified modules of genes with correlated expression in DEL, DUP and CTRL samples. Overall, we identified 11, 63 and 41 modules in iPSC, 1M and 3M organoids, respectively (**Table S6**). When these modules were statistically tested for association with DEL and DUP genotypes, thirty five modules (7 in iPSC, 7 in 1M, and 21 in 3M organoids) were detected as positively or negatively associated with genotypes at 10% FDR (**figs. S9-S11**). We detected a single module in each dataset that contained 16p11.2 CNV genes (*10purple* in iPSCs, *11greenyellow* in 1M, and *16lightcyan* in 3M organoids), and this module was positively associated with DUPs and negatively associated with DELs at each of the time points (**fig. S12**). Other notable modules included cell migration and motility (*22darkgreen* in 1M and *32violet* in 3M organoids), synaptic signaling and neuron differentiation (*46brown4* in 1M and *25orange* in 3M organoids), chromatin organization (*11greenyellow* in iPSCs), cilium assembly (*19lightyellow* in 3M organoids), and mitochondrial respiration (*3brown* in 3M organoids) (**Table S7**). Whereas some modules were unique to iPSCs or organoids, modules with migration and synaptic/neuronal functions were shared between 1M and 3M organoids. Interestingly, these modules were associated with genotypes in the opposite directions – migration module had negative association with DELs (*22darkgreen*) and positive association with DUPs (*32violet*), while synaptic/neuronal module had positive association with DELs (*46brown4)* and negative with DUPs (*25orange*) (**figs. S10-S11**). This suggests that 16p11.2 CNV differently impacts migration and neurogenesis.

To further investigate how co-expression modules and their function contribute to existing knowledge of ASD genetics, we performed statistical enrichment analyses against curated gene lists with previous evidence for involvement in autism (**STAR Methods**). We observed one module in each dataset with similar enrichment signatures (*9magenta* M9 in iPSCs, *6red* M6 in 1M and *2blue* M2 in 3M organoids) (**Fig. 2D**). These modules were enriched in highly confident ASD risk genes, constrained and highly intolerant to mutations (pLI>0.99) genes, as well as CHD8 and FMRP target genes in all datasets. GO analyses revealed shared biological functions related to histone modification and chromatin organization, with many ASD risk genes found within these modules. Chromatin modifying and remodeling genes (CHD8, ARID1B, ASH1L, KMT2A, SETD5) are known to be frequently mutated in ASD patients, suggesting that 16p11.2 CNVs impact gene regulatory networks that overlap with other ASD (and NDD) genes. We also observed several modules enriched in presynaptic or postsynaptic genes. In summary, both DEG and WGCNA analyses, suggest that the processes dysregulated by the 16p11.2 CNV at the transcriptome level converge on migration, synaptic/neuronal and chromatin-related functions.

### The 16p11.2 CNV impacts organoids proteome

In addition to impacting organoid’s transcriptome, the deletion and duplication of 29 genes within 16p11.2 CNV could have profound impact at the post-transcriptional level. To fully characterize the impact of the 16p11.2 CNV and to detect underlying molecular mechanisms, we performed proteomic profiling of organoids with Tandem Mass Tag Mass Spectrometry (TMT-MS), from the same samples as those used for RNA-seq experiments (**fig. S13**). We detected a total of 6,126 proteins in 1M and 5,481 proteins in 3M organoids, with 13 and 11 proteins from within 16p11.2 CNV, respectively. When DEL and DUP proteomes were compared to each other, we identified 305 and 970 differentially expressed proteins (DEPs) at 10% FDR in 1M and 3M organoids, respectively (**Fig. 3A-B** and **Table S8**). In proteomic data, the *cis*-effect of 16p11.2 CNV was weaker than in RNA-seq, possibly due to lower dynamic range between RNA and protein detectability in transcriptomic and proteomic experiments ^45^. However, patterns of proteome-wide effect of the 16p11.2 CNV on proteins outside of the *locus* were similar to transcriptome-wide effect, with more DEPs observed in 3M organoids compared to 1M organoids. Biological functions of DEPs agreed remarkably well with those of DEGs, with most notable pathways related to actin cytoskeleton, synaptic, neuronal functions and locomotion (**Fig. 3A-B** and **Table S9**). The DEPs shared by 1M and 3M organoids included synaptic (SYN1, STX1B, SYNJ1), cytoskeletal (MAPT, TUBB4A, TRIO) and cell adhesion (NCAM, CNTN1) proteins that were downregulated in DUPs and upregulated in DELs. Similar trends were observed for several high-confident autism-associated proteins (ANK2, DPYSL2, STXBP1, and DYNC1H1). This suggests that 16p11.2 CNV impacts proteins outside of the *locus,* with particular effect on cytoskeletal, synaptic, and autism-relevant proteins.

**Figure 3.**
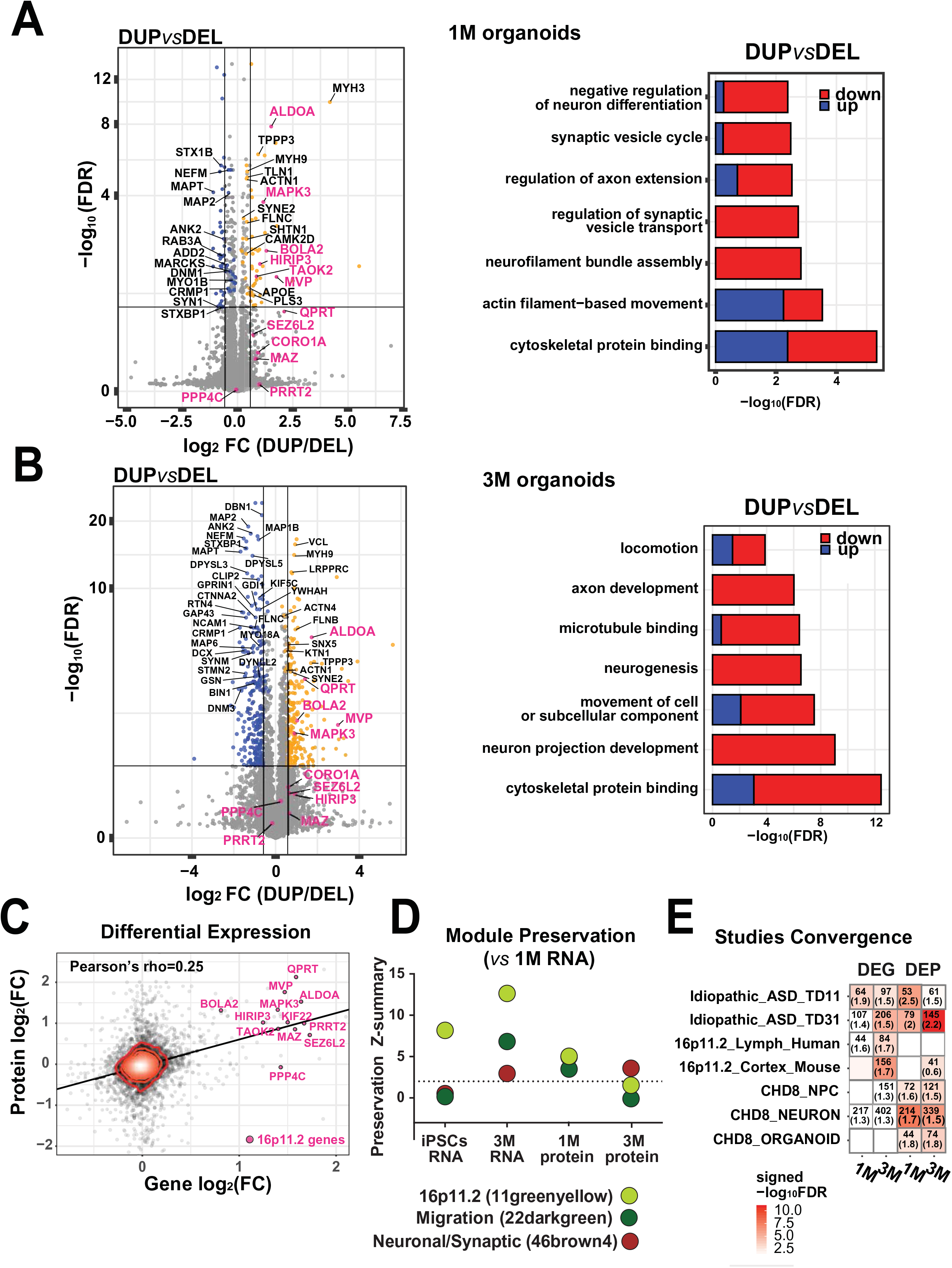
Proteomic analyses of 1M and 3M cortical organoids and correlation of protein expression with transcriptome. (A-B) Volcano plots of differentially expressed proteins between DEL and DUP 1M and 3M organoids derived from LC-MS/MS with TMT labeling proteomic profiling. Proteins within 16p11.2 CNV are colored in pink. Proteins colored in orange are upregulated in DUP and downregulated in DEL; proteins colored in blue are downregulated in DEL and upregulated in DUP. Gene Ontology enrichment analyses are shown as bar plots on the right. The contribution of up- or down-regulated proteins to specific GO terms are shown in blue and red, respectively. (C) Correlation of differentially expressed genes and proteins in 1M organoids. Genes/proteins within 16p11.2 CNV locus are colored in pink. (D) Module preservation analyses for migration, neuronal/synaptic and 16p11.2 modules detected in transcriptomic profiling of 1M organoids as compared to other datasets (iPSC and 3M transcriptomic, and 1M and 3M proteomic datasets). The neuronal/synaptic module *vs* 1M Protein is located behind 16p11.2 module and is not visible. (E) Comparison of differentially expressed genes and proteins from this study with differentially expressed genes from other relevant studies. Idiopathic_ASD_TD11 and Idiopathic_ASD_TD31 are differentially expressed genes from organoids derived from individuals with idiopathic ASD at 11^th^ and 31^st^ day of differentiation from Mariani study ^30^; 16p11.2_Lymph_Human and 16p11.2_Cortex_Mouse are differentially expressed genes from human lymphoblastoid cell lines of ASD patients with 16p11.2 CNV and 16p11.2 deletion mouse cortex, respectively, from Blumenthal study ^13^; CHD8_NPC, CHD8_NEURON and CHD8_ORGANOID are differentially expressed genes from isogenic CRISPR/Cas9 generated heterozygous CHD8 neural progenitors, monolayer neurons and organoids from Wang study ^48^. Number of overlapped genes and odds ratio (in parenthesis) are indicated inside each cell, and provided only for FDR≤0.05 and OR>1.

To further investigate how protein co-expression modules are impacted by the 16p11.2 CNV in DEL and DUP patient-derived cortical organoids, we performed Weighted Protein Co-expression Network Analysis (WPCNA) using TMT-MS proteomic data. We identified 21 and 17 protein co-expression modules in 1M and 3M organoids, respectively (**Table S10**). Twelve modules (5 in 1M and 7 in 3M organoids) were associated with DEL or DUP genotypes at 10% FDR (**figs. S14-S15**). The dysregulated modules included those enriched in RNA splicing and chromatin organization (*5green* in 1M organoids), ribosome and translation (*2blue* in 3M organoids), cytoskeleton and microtubule (*7black* in 3M organoids), and mitochondrial respiration (*8pink* in 3M organoids) GO functions (**Table S11**). One module detected in 3M organoids by WPCNA, *1turquoise* (M1), was enriched in pre- and postsynaptic, constrained and FMRP target proteins (**fig. S16**). It included proteins involved in neuron differentiation and neurogenesis, neuron projection development, synaptic signaling, cytoskeleton organization, actin filament processes, as well as migration and locomotion. All these functions were also identified by RNA-seq profiling, pointing to convergence of molecular processes at the transcriptome and proteome levels.

### Biological convergence of organoids transcriptome and proteome

To determine the extent of convergence between organoids transcriptomes and proteomes, we calculated correlation coefficient of expression levels for genes and proteins (**Fig. 3C**). Globally, we observed positive correlation between DEGs and DEPs in 1M organoids (Pearson r=0.25) (**Fig. 3C**) and in 3M organoids (Pearson r=0.1) (**fig. S17**). We then carried out module preservation analyses to identify conserved modules across these two levels of regulation (**Fig. 3D**). This analysis demonstrated high degree of preservation at the RNA and protein level for most modules that were significantly associated with genotype (**fig. S18**), and especially for 16p11.2, migration and neuronal/synaptic modules in both 1M (**Fig. 3D**) and 3M (**fig. S19**) organoids. The module containing 16p11.2 genes had highest conservation within transcriptome modules (**Fig. 3D**). Overall, we observed remarkably high correlation between organoids transcriptomes and proteomes, given previously noted lack of conservation at these two levels of cellular regulation observed in other studies ^46, 47^.

To put our results into a context of previous studies, we performed enrichment analyses our DEGs and DEPs in other datasets with relevance to ASD. Specifically, our DEGs and DEPs were compared with transcriptomes of 16p11.2 patients’ lymphoblast lines and cerebral cortex of 16p11.2 mice ^13^, idiopathic ASD patient-derived organoids ^30^ and CHD8 KO organoids, NPCs and neurons ^48^ (**Fig. 3E**). Overall, we observed greater overlap of our DEGs and DEPs with DEGs identified in idiopathic ASD organoid models, suggesting that 16p11.2 organoids share transcriptomic signatures with other ASD subtypes. With regards to transcriptomes from 16p11.2 patients’ lymphoblastoid cell lines and 16p11.2 mouse cortex, our 3M organoids captured transcriptomic signatures of these models better than 1M organoids, potentially due to their more advanced stages of maturation. We observed lesser overlap of our DEPs with DEGs from human lymphoblastoid lines and adult mouse cortex, potentially reflecting general low correlation between protein and mRNA levels due to various biological and technical factors ^46, 47^. In summary, we observed greater overlap of our 16p11.2 organoid data with organoid models of idiopathic ASD and other ASD genes than with 16p11.2 models from human lymphocytes or mouse brain. This highlights the importance of using human-derived models for investigating neurodevelopmental disorders, and highlighting similarities between different genetic subtypes of ASD.

### The dosage of 16p11.2 CNV alters cell type composition of organoids

The findings from transcriptomic and proteomic analyses highlight molecular processes that are disrupted by the 16p11.2 CNV in the context of fetal brain development. Given complex cell type composition of human brain, these signatures may be in part related to effects of the CNV on cell-type composition of the organoids. To better understand how 16p11.2 dosage may impact cell type composition of organoids, we performed cell type enrichment analyses of organoid transcriptomes using single cell RNA-seq (scRNA-seq) from the developing human neocortex ^49^.

We have previously demonstrated by scRNA-seq that at 1M organoids primarily consist of progenitor cells, with smaller fractions of glutamatergic neurons, glial cells, and intermediate progenitors ^32^. Here, we used recent scRNA-seq data from fetal human neocortex ^49^ to identify cell types significantly enriched in 1M and 3M old organoids. We observed significant enrichment of different cell types in co-expression modules for 1M and 3M organoids (**Fig. 4A**). The analyses of the significant modules by genotype at 1M revealed that DEL organoids were enriched in neuronal cell types (**Fig. 4B**). Interestingly, DUPs were enriched in intermediate progenitors and radial glia (**Fig. 4C-D**). In support of cell type enrichment results, GO functions of most significant modules reflected processes typically associated with corresponding cell types (**Fig. 4B-D, right panels**). For example, GO functions for 1M *45darkorange2* “Neuron” cell type module included “neurogenesis,”, “neuron development” and “neuron differentiation”. The GO functions of the intermediate progenitor “IP” *42lighcyan1* module included “pattern specification process”, “nervous system development”, and “generation of neurons”, along with differentiation- and proliferation-related functions. The GO functions for radial glia “RG” *2blue* module captured cilium and microtubule-based processes. These results support a hypothesis that 16p11.2 copy number has a quantitative effect on the ratio of neurons to progenitor cells, with duplications having a reduced proportion of neurons and deletions having an excess.

**Figure 4.**
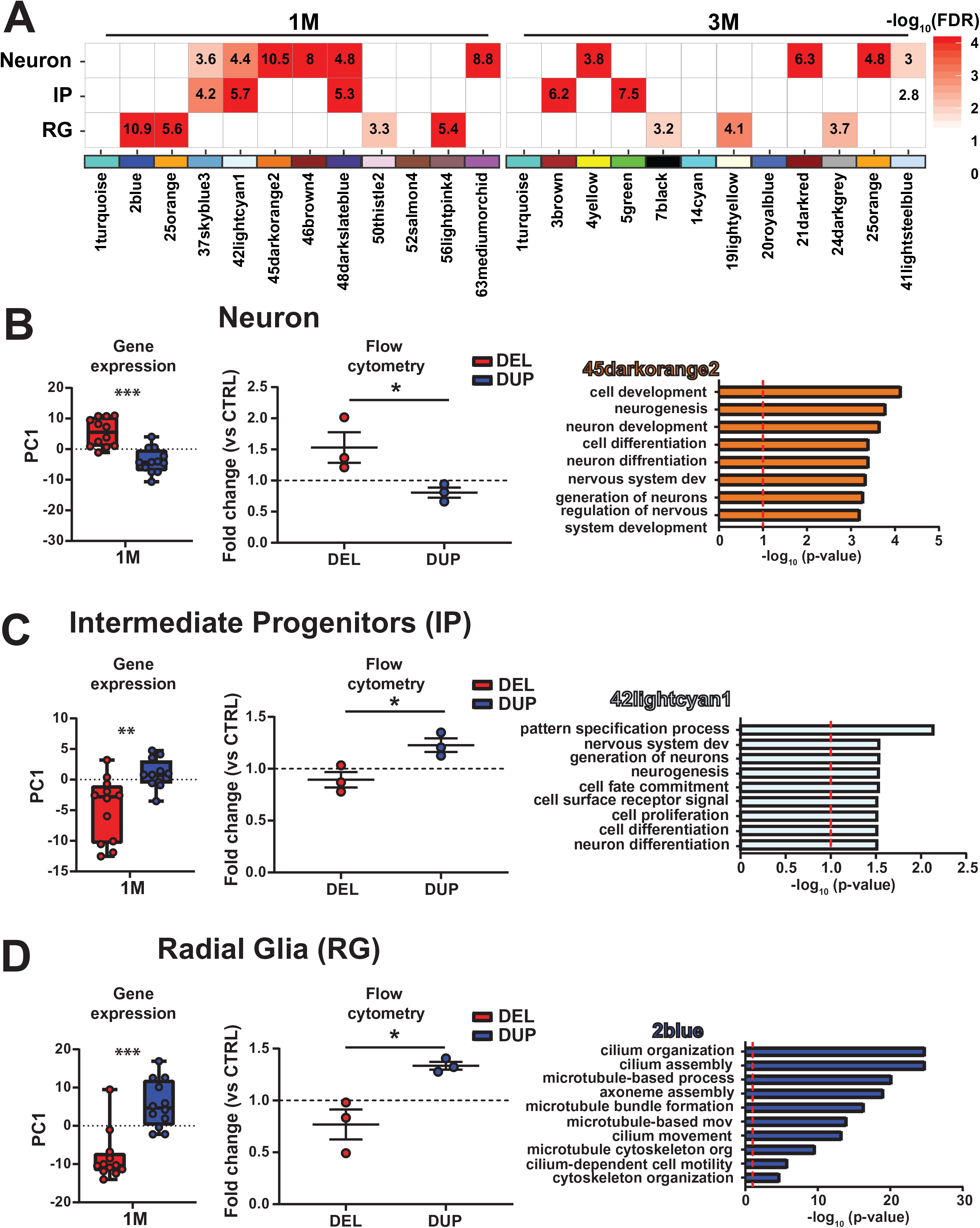
Cell type enrichment analyses of 1M and 3M organoid transcriptomes. (A) Cell type enrichment analyses of RNA-seq co-expression modules from 1M and 3M old organoids using cell types from scRNA-seq of fetal neocortex ^49^. Modules significantly enriched in at least one cell type are shown. “Neuron” category includes combination of excitatory and interneurons; IP – intermediate progenitors; RG – radial glia. Enrichment was evaluated using bootstrapping. Z-score was estimated by the distance of the mean expression of the target gene set from the mean expression of bootstrapping replicates and is shown inside each cell. P-values were corrected for multiple comparisons using FDR. (B-D) Right panel: Principal Component 1 (PC1) of enriched organoid modules at 1M plotted by genotype. PC1 was computed for a union of genes from all modules significantly enriched in a specific cell type. All comparisons between DELs and DUPs are significant using t-test statistics. \*\*\**p*<0.001, \*\**p*<0.01. Middle panel: Flow cytometry analysis of dissociated cerebral organoids. Graphs represent quantification of the percentage of each cell population compared to CTRL. The data shown is representative of three independent experiments (\**p*≤0.05). Right panel: GO terms for one representative most enriched module from 1M dataset were obtained using g:Profiler.

We sought to test this hypothesis directly by flow cytometry analysis on 1M dissociated cerebral organoids (**STAR Methods**). Single-cell suspensions were labelled with NeuN, TBR2, and SOX2 for Neurons, Intermediate Progenitors (IP) and Radial Glia (RG), respectively, and the percentages of positive cells were quantified (**fig. S20**). We observed that the percentages of positively labeled cells between genotypes from flow cytometry experiments correlated well with single cell enrichment analyses from gene expression. The percentage of NeuN^+^ cells was significantly higher in DELs compared to DUPs, suggesting an increase in the number of neurons (**Fig. 4B, middle panel**). In contrast, the percentages of TBR2^+^ and SOX2^+^ cells were significantly higher in DUPs, suggesting an increase in progenitor populations of RG and IP (**Fig. 4C-D, middle panel**). These results point to potentially increased neuronal maturation in DEL organoids, and the opposite effect in DUP organoids. Overall, cell type enrichment results provided further insight into cell type composition of organoids and correlated with previous findings from ASD brain. For instance, excess neuron number has been observed as a hallmark of brain overgrowth in ASD patients during first year of life ^50, 51^, supporting “Neuron” cell type enrichment and macrocephaly phenotype in DELs.

### Increased neuronal maturation in 16p11.2 DEL organoids

Transcriptome signatures and analysis of cell types suggests that 16p11.2 copy number quantitatively affects the proportion of neurons and neural progenitor populations. In addition, transcriptomic module *46brown4* in 1M organoids was enriched in “Neuron” cell type, among other neuron-enriched modules (**Fig. 4A**). This module was significantly upregulated in DELs (**Fig. 5A**), and contained genes with neuronal and synaptic functions (**Fig. 5B**). The expression levels of genes from this module highly correlated with corresponding protein expression (Pearson correlation coefficient PCC=0.62), with 42.2% of genes within this module also detected by proteomics (**Fig. 5C**). One of the high-confidence autism risk genes, *SCN2A* ^52^, was a highly connected hub in this module (**Fig. 5D**).

**Figure 5.**
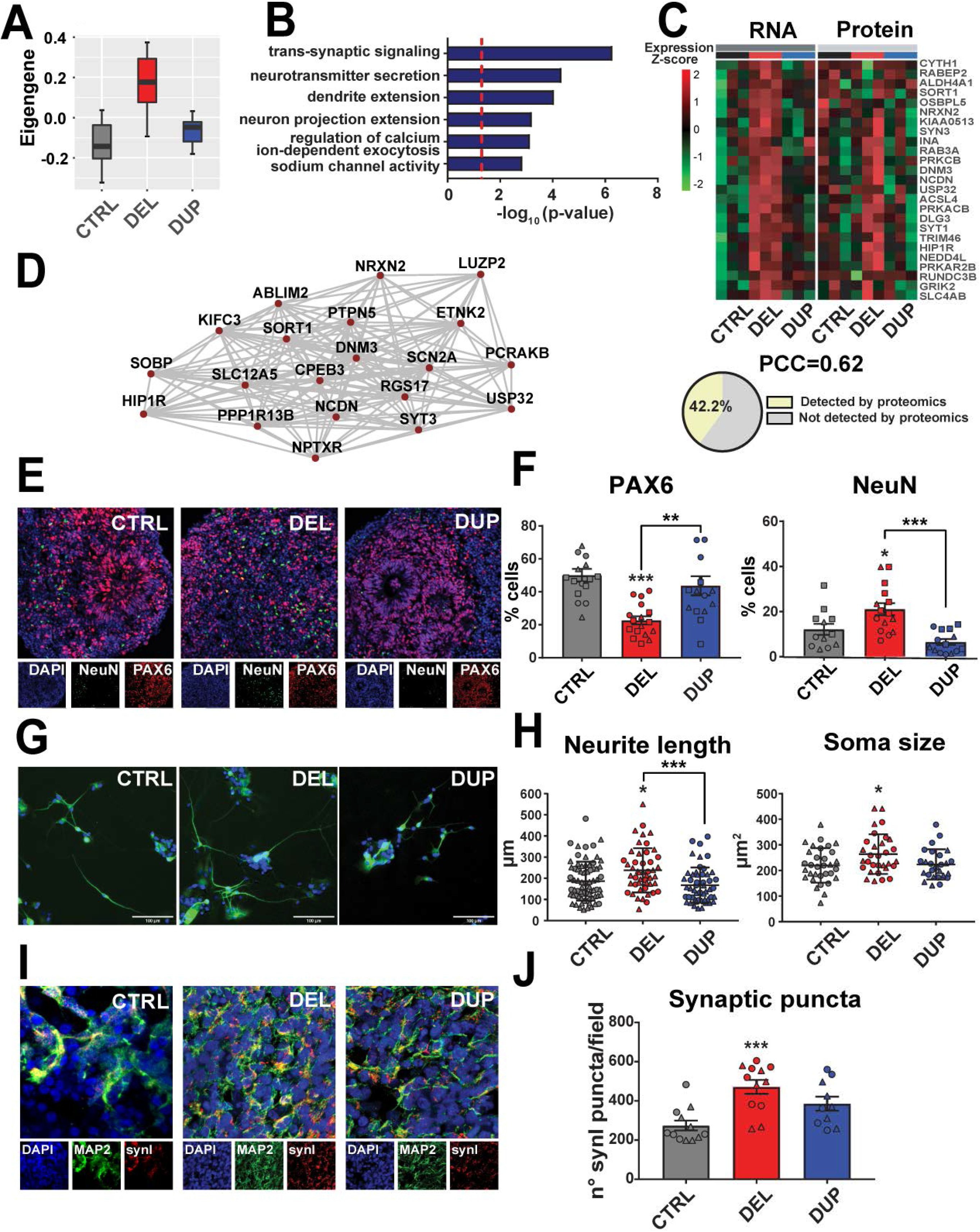
Altered neuronal maturation, morphology and synaptic defects in 16p11.2 cortical organoids. (A-B) Module eigengene and GO functional annotations for *46brown4* transcriptomic module in 1M organoids. Dots (*n*=12) correspond to each replicate derived from 3 patients (CTRL, DEL or DUP), 2 clones per patient and 2 replicates per clone. Two replicates were removed from CTRL before the analyses during outlier detection procedure (see **STAR Methods**). (C) Heat plot representing gene (RNA) and protein expression from *46brown4* module. Pearson correlation coefficient between RNA and protein expression and the proportion of genes whose protein products were also detected by proteomics are shown below the plot. (D) Twenty top hub genes from *46brown4* module. Edges represent co-expression. (E-F) Representative images of 1M organoid slices (CTRL, DEL and DUP) immunostained with DAPI, PAX6 and NeuN. Quantification of the percentage of positive cells for each marker is shown. Symbols represent organoids from the same differentiation batch, where batch is defined as CTRL, DEL, DUP from one patient, one clone and one replica. Data is presented as mean ± SEM (*n=*2 patients per genotype, at least 4 organoids per patient). Significance was calculated using one-way ANOVA; \*\*\**p*<0.001, \*\**p*<0.01, \**p*≤0.05. Significance above bars represents comparison against CTRL. (G-H) Representative images of neurons from dissociated 1M organoids immunostained with DAPI (blue) and MAP2 (green). Quantification of total neurite length and soma size is shown. Symbols represent neurons derived from organoids from the same differentiation batch. Data is presented as mean ± SD (*n=*2 patients per genotype, at least 15 neurons per patient). Significance was calculated using one-way ANOVA; \**p*≤0.05. Significance above bars represents comparison against CTRL. (I-J) Representative images of 1M organoid slices immunostained with DAPI, MAP2 and SynI. Quantification of the total Synapsin I to estimate synaptic puncta is shown. Symbols represent organoids from the same differentiation batch. Data is presented as mean ± SEM (*n=*2 patients per genotype, at least 3 organoids per patient). Significance was calculated using one-way ANOVA; \*\*\**p*<0.001. Significance above bars represents comparison against CTRL.

To validate these findings experimentally and to better understand cellular basis of neuronal dysregulation in organoids, we quantified neural progenitors and neurons by immunohistochemistry in 1M organoid slices. We observed mirror cellular phenotypes in DEL and DUP organoids: DEL organoids had greater number of neurons (i.e. NeuN^+^ cells) and lower number of neural progenitors (Pax6^+^), and DUP organoids had an opposite phenotype (**Fig. 5E-F**), which is also consistent with flow cytometry results. This confirms that DEL organoids mature faster than DUP organoids, and that progenitor proliferation and differentiation dynamics is disrupted by the 16p11.2 CNV. Higher number of neurons in DELs is also in agreement with the increased expression of synaptic genes that we have observed by transcriptomic profiling. The DEL organoids also had decreased proliferation rate, most likely due to the depletion of progenitor pool by 1M (**fig. S21**). Given an increased number of neurons in DELs at 1M, it is plausible that increased proliferation prior to 1M could lead to depletion of progenitors by 1M. Indeed, accelerated proliferation of neural progenitors from iPSCs has been previously quantified at much earlier time point than 1M in other models ^53^, suggesting that at 1M we may be capturing later or even terminal stages, at which progenitor pool in DELs has already been depleted. Cell cycle exit determined by the ratio of Edu^+^ and Ki67^-^ cells was not affected in 1M DEL or DUP organoids. The summary of this and all follow-up experiments by clones and replicates is shown in **Table S12**.

### Neuronal morphology and synaptic defects in 16p11.2 organoids

Neuronal maturation defects in DEL and DUP organoids, along with differences in their size suggest that neuronal morphology could be affected by the 16p11.2 CNV. To test this hypothesis and to replicate previous observations from 2D neuronal cultures of the 16p11.2 carriers ^54^, we investigated neuron morphology by measuring soma size and neurite length in the dissociated 1M organoids (**STAR Methods**). We have previously demonstrated that cortical organoids at 1M of differentiation produce functional cortical neurons ^32^. Therefore, we stained dissociated cells with MAP2 neuronal marker and performed measurements eight days after plating. We observed increased soma size in DEL organoids compared to CTRL (one-way ANOVA p=0.034) (**Fig. 5G-H**). The total neurite length was increased in DEL *vs.* CTRL (one-way ANOVA p=0.01), and in DEL *vs.* DUP (p=0.0007), with a trend for decreased neurite length in DUPs *vs* CTRL that did not reach statistical significance. These results suggest that one of many factors contributing to organoid size differences could be soma size and neurite length.

Changes in neuronal morphology together with altered neuronal maturation could impact synaptogenesis in organoids. We therefore analyzed synaptic puncta by co-staining 1M organoid slices with presynaptic marker Synapsin-I (SynI) and neuronal marker MAP2. We observed significant increase in the number of synaptic puncta in DEL organoids (one-way ANOVA p=0.0003) (**Fig. 5I-J**). This result is in agreement with the increased number of neurons, and with upregulation of neuronal/synaptic transcriptomic module observed in DELs.

### Severe neuronal migration defects in 16p11.2 organoids

Neuronal migration during early fetal brain development could be one of the mechanisms that is disrupted in neurodevelopmental disorders. Here, we observed that gene sets and modules involved in neuronal migration and locomotion were dysregulated across 16p11.2 transcriptomes (DEG and WGCNA) and proteomes (DEP and WPCNA). For example, transcriptomic *22darkgreen* module from 1M organoids was significantly downregulated in DELs, and annotated with locomotion, migration, and motility (**Fig. 6A-B**). Other GO functions within this module included Wnt signaling and regulation of Rho protein signal transduction, both related to cell migration and cytoskeletal functions. Specifically, many genes from Wnt signaling pathway were strongly downregulated in DEL organoids (**fig. S22**). Similarly to synaptic module, gene and protein expression of *22darkgreen* migration module were positively correlated (Pearson correlation coefficient PCC=0.46) **(Fig. 6C)**. One of the genes in this module, ARHGEF2, is a microtubule-regulated Rho guanine exchange factor, known to be involved in cell migration (**Fig. 6D**).

**Figure 6.**
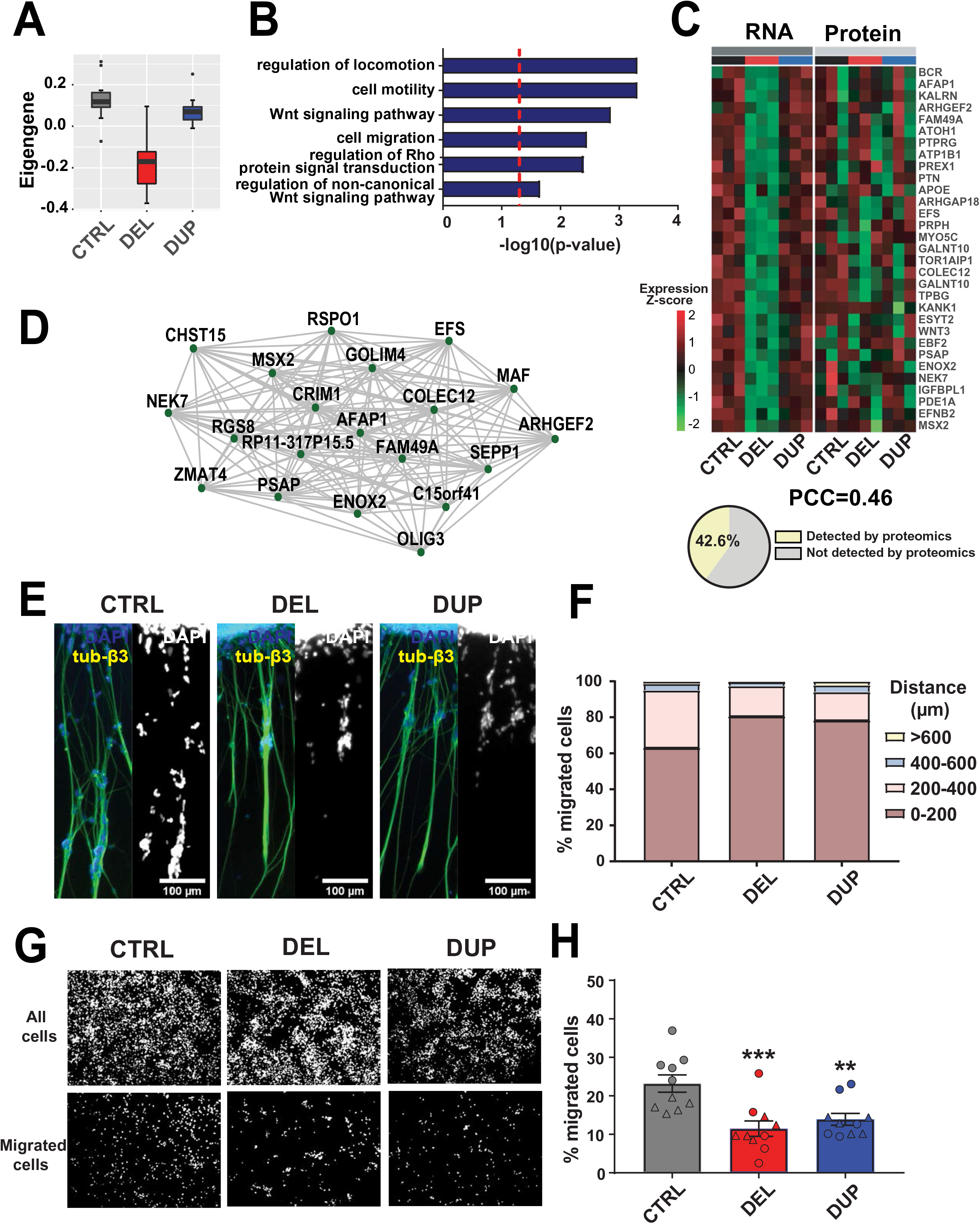
Neuronal migration defects in 16p11.2 cortical organoids. (A-B) Module eigengene and GO functional annotations for *22darkgreen* transcriptomic module in 1M organoids. Dots (*n*=12) correspond to each replicate derived from 3 patients (CTRL, DEL or DUP), 2 clones per patient and 2 replicates per clone. Two replicates were removed from CTRL before the analyses during outlier detection procedure (see **STAR Methods**). (C) Heat plot representing gene (RNA) and protein expression from *22darkgreen* module. Pearson correlation coefficient between RNA and protein expression and the proportion of genes whose protein products were also detected by proteomics are shown below the plot. (D) Twenty top hub genes from *22darkgreen* module. Edges represent co-expression. (E-F) Representative images of 1M cortical organoids 72hrs after attachment to Matrigel-coated plates, immunostained with DAPI and β-tubulin III. Graph represents percentages of migrating cells to a distance with 200-μm bins of displacement (n=2 patients per genotype, at least 5 organoids per patient). (G-H) Representative images of cells migrating from 1M dissociated organoids in Boyden chamber experiment. Immunostaining was performed with DAPI. Quantification of the percentages of migrated cells is shown. Symbols represent cells derived from organoids from the same differentiation batch. Data is presented as mean ± SD (*n=*2 patients per genotype, at least 5 images per patient). Significance was calculated using one-way ANOVA; \*\*\**p*<0.001, \*\**p*<0.01. Significance above bars represents comparison against CTRL.

To investigate migration defects in the 16p11.2 organoids, we performed two orthogonal *in vitro* migration assays as previously described ^29, 55^. First, we seeded organoids onto matrigel-coated plates, and quantified the number of migrated neurons and migration distance 72 hours after seeding. Within the first 24 hours after plating, protrusions of radial glia fibers from the organoid edges were observed. Then, neurons started to migrate along these fibers. While about 40% of neurons migrated to a distance of over 200 μm along the fibers in the CTRL organoids, only about 20% of neurons migrated to the same distance from the DEL or DUP organoids in this experiment (one-way ANOVA DEL/CTRL p=0.038; DUP/CTRL p=0.073) (**Fig. 6E-F**). Live imaging further verified that migration distance is shorter for both DEL and DUP organoids (**fig. S23 and movie S1**). We verified by immunostaining that the fibers consist of both neurites and radial glia bundles, and that the migrating cells are neurons and not neural progenitors (**fig. S24**). The orthogonal Boyden chamber assay further confirmed migration defects in 16p11.2 organoids (**Fig. 6G-H**). These results suggest that neuron migration defects are observed in both, DEL and DUP organoids, and that these abnormalities could be present in 16p11.2 carriers during fetal brain development.

### Inhibition of RhoA activity rescues migration defects in 16p11.2 organoids

Rho signaling is one of the pathways enriched in the migration module along with other pathways (**Fig. 6B**), consistent with previous findings from our group ^41^ and replicated by others ^56, 57^. Furthermore, RhoA is known to regulate neuronal migration, actin dynamics and neurite outgrowth during brain development ^58–60^. These observations suggest a hypothesis that RhoA signaling may be one pathway contributing to the neuronal migration defects that we observed.

We confirmed by Western Blot that RhoA is dysregulated in 16p11.2 organoids (**Fig. 7A-B and fig. S25**). KCTD13 protein level was significantly decreased in DEL and significantly increased in DUP organoids compared to CTRL (CI interval test, CTRL *vs* DEL p<0.001, CTRL *vs* DUP p<0.001), mirroring 16p11.2 CNV dosage. As would be predicted based on the expression of KCTD13, an adapter protein that regulates ubiquitination of RhoA, total RhoA levels had an inverse trend (CI interval test, CTRL *vs* DEL p<0.05, CTRL *vs* DUP p<0.01), as we have previously hypothesized ^41^. However, active GTP-bound form of RhoA (RhoA-GTP) was significantly upregulated in organoids of both genotypes (CI interval test, CTRL *vs* DEL p<0.05, CTRL *vs* DUP p<0.01) (**Fig. 6E-H**). These results indicate that the active form of RhoA was upregulated in both DELs and DUPs, consistent with the observed migration phenotype, and suggest putative dysregulation of the RhoA signaling pathway, either directly by the 16p11.2 CNV, or by other genes outside of the *locus* that this CNV is impacting.

**Figure 7.**
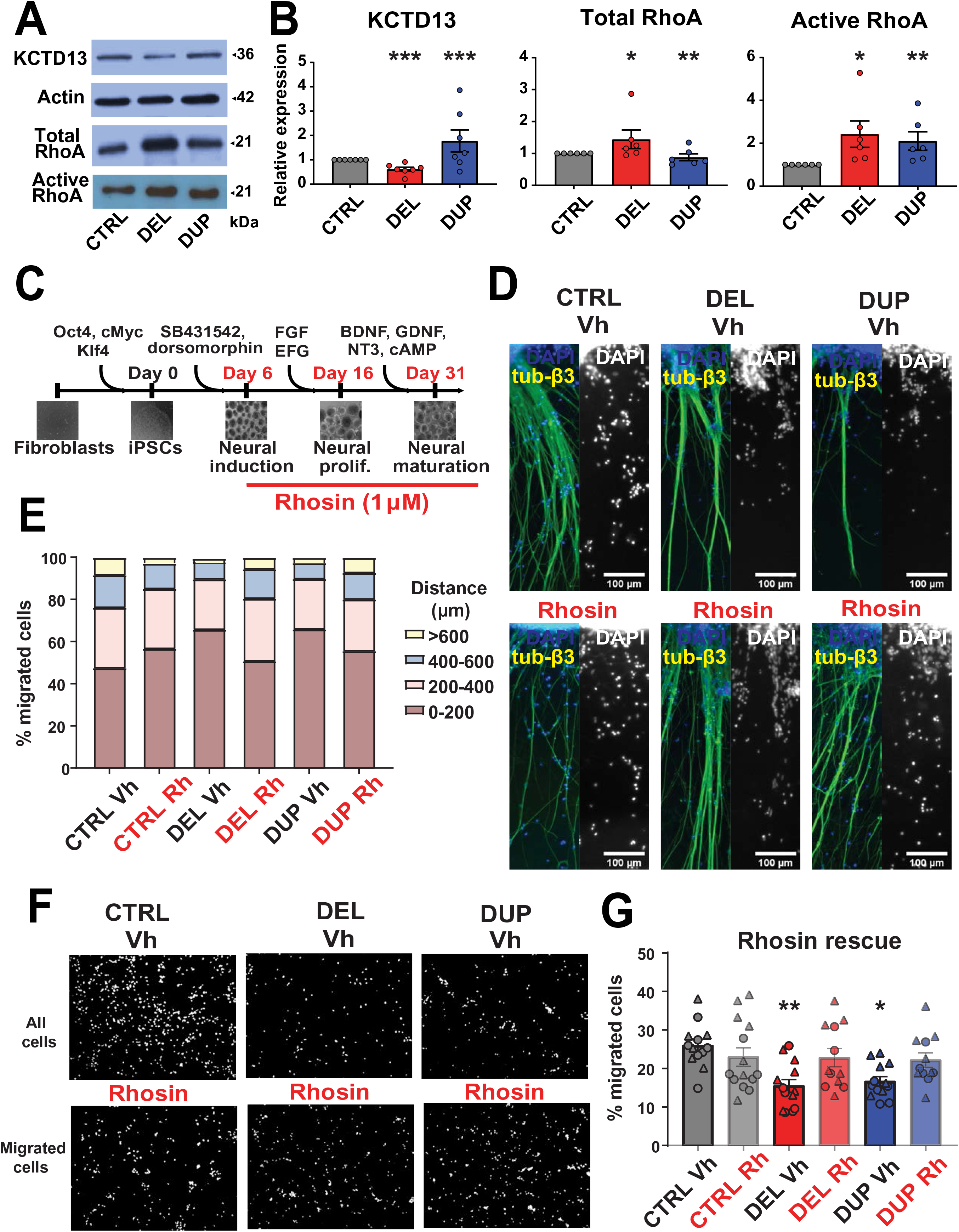
Rhosin treatment rescues neuronal migration deficits in 16p11.2 cortical organoids. (A-B) Representative images of Western Blot analysis of 1M organoids for KCTD13, actin as loading control, total RhoA, and active RhoA (RhoA-GTP). Densitometry analysis of Western Blot is shown. Data is represented as mean ± SEM (*n*=7 differentiation batches with at least one batch for each patient or control for KCTD13, n=6 batches with at least one batch for each patient or control for total RhoA, n=6 batches with at least one batch for each patient or control for active RhoA). Significance was calculated using Confidence Interval (CI) method; ***p<0.001, **p<0.01, *p≤0.05. Significance above bars represents comparison against CTRL. (C) Schematic representation of constitutive Rhosin treatment of organoids during differentiation. (D-E) Representative images of 1M vehicle- or Rhosin-treated cortical organoids 72hrs after attachment to Matrigel-coated plates, immunostained with DAPI and β-tubulin III. Graph represents percentages of migrating cells to a distance with 200-μm bins of displacement (n=2 patients per genotype, at least 5 organoids per patient). (F-G) Representative images of cells migrating from vehicle- or Rhosin-treated 1M dissociated organoids in Boyden chamber experiment. Immunostaining was performed with DAPI. Quantification of the percentages of migrated cells is shown. Symbols represent cells derived from organoids from the same differentiation batch. Data is presented as mean ± SD (*n=*2 patients per genotype, at least 5 images per patient). Significance was calculated using one-way ANOVA; \*\**p*<0.01, \**p*≤0.05. Significance above bars represents comparison against CTRL.

Neuron migration defects in 16p11.2 organoids were then rescued by inhibition of RhoA activity. After constitutive treatment of intact organoids with RhoA inhibitor Rhosin ^61^ starting from 6^th^ day of differentiation (**Fig. 7C**), migration defects in both DEL and DUP were rescued to the levels indistinguishable from CTRL (one-way ANOVA CTRL *vs* DEL_Rh p=0.53; CTRL *vs* DUP_Rh p=0.36) (**Fig. 7D-E**). The orthogonal Boyden chamber experiments replicated migration phenotype rescue with Rhosin (**Fig. 7F-G**). However, the increased neurite length in DELs was not rescued by Rhosin treatment (**fig. S26**). This suggests that Rho signaling may be one of the pathways that is contributing to decreased migration, but not to neurite length, and points to other potential pathways that may be involved in 16p11.2-impacted phenotypes.

## Discussion

Patient-derived and CRISPR/Cas9 genome-engineered isogenic brain organoids are becoming popular models for investigating molecular mechanisms underlying neurodevelopmental disorders ^25, 62^. Given lack of fetal brain tissues from ASD patients, there are numerous advantages in using brain organoids to model patient phenotypes. In the present study we model molecular and cellular mechanisms of ASD risk attributable to rare deletions and duplications at 16p11.2 *locus* using cortical organoids derived from skin fibroblasts of ASD patients with macrocephaly or microcephaly, respectively.

Organoid models of 16p11.2 CNV exhibit defects in neuronal maturation, migration, morphology and synaptic abnormalities. These genetic effects are consistent with brain growth phenotypes observed in the patients carrying 16p11.2 DEL or DUP. Accelerated neuronal maturation in DEL and delayed in DUPs, along with impaired neuron migration, are novel mechanisms that have not been previously implicated in 16p11.2-linked autism. Our study makes significant contribution to mechanistic understanding of cellular and molecular processes that may be disrupted during early neocortical development in the 16p11.2 CNV carriers.

In this study, we perform in-depth characterization of organoids’ transcriptomes and proteomes in parallel, from the same samples, at different developmental time points. This strategy provides the leverage for comparing mirror phenotypes of DELs and DUPs at two levels of regulation, transcriptional and translational. We obtain highly consistent results at the level of genes and proteins, by two independent methods, RNA-seq and quantitative proteomics.

At the molecular level, we observe perturbations of transcriptional programs associated with key processes involved in neurodevelopment. Dysregulation of genes with neuron differentiation, synaptic signaling and cortical layer formation functions are observed in organoids, but not in the iPSC lines. This suggests that disruption of neural processes, which may not be apparent at the very early embryonic stages in iPSCs, become more pronounced during organoids maturation. Transcriptional signatures of one month old organoids recapitulate those of the late mid-fetal human brain development, the most critical period for establishing network connectivity among 16p11.2 CNV genes ^41^. This period was also implicated in ASD by other studies ^39, 63^. Most importantly, dysregulated transcriptional modules associated with synaptic functions and neuronal migration identified in organoids were conserved at the proteomic level. Preservation of transcriptional signatures at the translational level further reinforces and validates our findings.

Our results are in agreement with other studies that include either organoids produced from idiopathic ASD individuals ^30^, or transgenic mice with 16p11.2 CNV ^13^. Significant overlap between differentially expressed genes and proteins from our study is observed with DEGs from Mariani ^30^, suggesting shared signatures among different genetic subtypes of ASD. Importantly, synaptic gene co-expression module is also dysregulated in organoids from idiopathic ASD patients. The degree of overlap is lower with DEGs from organoids engineered to knock-down CHD8, a top autism gene ^48^. It is likely that patient’s overall genetic background may be an important contributor to the observed transcriptional signatures. With regard to the 16p11.2 mouse model, our 3M organoids capture greater signal from 16p11.2 adult mouse brain than 1M organoids, likely due to their more advanced stages of maturation with closer similarity to brain tissues.

As observed previously in other brain diseases, organoid models can recapitulate patient’s microcephaly ^20, 29, 33^ and macrocephaly ^48, 64^ phenotypes. Here, we demonstrate that dosage changes of the same genetic variant could lead to opposite organoids size. In addition, we replicate altered neuronal morphology observed in 2D studies ^54^, suggesting that it may be one of the phenotypes contributing to size differences between DELs and DUPs. Aberrant control of cell proliferation and excess neuron number has been previously hypothesized to cause early brain overgrowth in ASD patients ^51, 65^. Consistent with this hypothesis, we observed excess of neurons and depletion of neural progenitors in DEL organoids, and a mirror phenotype was found in DUP organoids. We also found decreased proliferation in 1M old DEL organoids. Interestingly, previous ASD studies either did not find differences in proliferation ^54^, or demonstrated accelerated proliferation ^30^. It is possible that decreased proliferation at 1M is a result of accelerated proliferation at earlier time points that leads to premature depletion of neural progenitor pool by 1M. Further investigation of proliferation rates at various developmental time points (iPSCs, NPCs, and early maturation in organoids) is needed, and could uncover time-dependent mechanisms of proliferation defects in ASD.

One of the important findings from our study is impaired cortical neuron migration in 16p11.2 organoids. We confirmed reduced migration in DEL and DUP organoids by two orthogonal methods, recordings from intact organoids, and experiments in dissociated organoids. Previously, neuronal migration defects have also been observed in organoids derived from patients with lissencephaly ^29^, periventricular heterotopia ^66^, and in CHD8 deficient mice ^67^. Our results suggest that aberrant neuronal migration may be present in the brains of subjects with 16p11.2 CNV during early neurogenesis. Notably, observations from post-mortem ASD brains show patches of disorganized cortical neurons that may not be migrating properly during early brain development ^50^.

Our observations linking RhoA activity in 16p11.2 organoids with defects in neuron migration are consistent with neuron migration phenotypes observed in mouse conditional knockout of RhoA ^59^. This study revealed that RhoA-depleted neurons migrated faster and reached cortical plate sooner than GFP+ control neurons upon Cre electroporation into cerebral cortex of E14 mouse embryos. It has also demonstrated that electroporation of spontaneously activated (‘‘fast-cycling’’) mutant of RhoA caused slower neuronal migration in the condition of activated RhoA. Thus, overactivation of RhoA stalls migration of neurons, in agreement with our results from the 16p11.2 CNV model. Our data is also consistent with previous work rescuing delayed neuronal migration by inactivation of RhoA or inhibition of ROCK, a direct target of RhoA ^68–70^. Here, we demonstrate that inhibition of RhoA with Rhosin rescues delayed migration in 16p11.2 DEL and DUP organoids.

There is a myriad of biological pathways that could be dysregulated by the 16p11.2 CNV in ASD ^71^. Due to its polygenic nature, with 29 genes within the *locus* and hundreds of genes impacted outside of the *locus,* as demonstrated here, genetic and epistatic interactions among these genes are likely responsible for neuroanatomical and cellular phenotypes observed in the patients and animal models ^72, 73^. Investigation of this CNV creates apparent challenges in implicating a specific pathway, mostly due to combinatorial and synergistic effect of multiple genes ^74^. Rather, dysregulation of multiple pathways could lead to observed cellular and molecular phenotypes. For example, a number of genes within 16p11.2 CNV (MAPK3, MVP and TAOK2), are involved in MAPK/ERK and phosphatidylinositol 3-kinase PI3K/AKT signaling pathways. These pathways, regulating cell cycle and proliferation of neural progenitors, were shown to be dysregulated in 16p11.2 deletion mouse model ^16^, and are likely to also be impacted by this CNV. Here, we identified modules with genes and proteins involved in Wnt signaling, suggesting that 16p11.2 CNV may also impact this pathway. Given the cross-talk between Rho and Wnt signaling pathways ^75, 76^ that are both involved in the regulation of neuronal cytoskeleton during axon and dendrite growth, along with synapse formation, it is plausible that Wnt signaling contributes to the neurite length phenotype in our 16p11.2 CNV organoid model. Finally, as we demonstrated here, RhoA signaling is likely regulating neuronal migration in our 16p11.2 organoid model, and inhibition of RhoA activity rescues migration deficits. Thus, pleiotropy and epistasis of 16p11.2 CNV genes at the pathway level is a hallmark of its functional impact. Future studies using organoid models or fetal brain tissues from 16p11.2 CNV carriers are required to untangle the complexity of the phenotype-pathway relationships in ASD.

## STAR Methods

### Study design

The aim of this study was to investigate the impact of the autism-associated 16p11.2 CNV on early brain development using human-derived models. Specifically, our goal was to detect molecular pathways dysregulated by the dosage changes (i.e. deletions and duplications) of this CNV comprising 29 genes. To address this question, we generated cortical organoids derived from fibroblasts, reprogrammed into iPSCs, of 16p11.2 patients and healthy controls. We selected three patients of each genotype (3 DEL, 3 DUP and 3 CTRL), on the basis on the extreme head size phenotype and ASD diagnosis (**Table S1**). Due to samples availability, our study has been restricted to one gender (male). Given ASD prevalence of 4:1 male:female ratio, this study design is justified. To investigate changes in RNA and protein expression caused by the 16p11.2, bulk RNA sequencing and quantitative label-free TMT-MS proteomics experiments were performed. We profiled 2 clones per patient, and 2 replicas per clone, at 3 developmental stages (iPSCs, 1 month old and 3 month old organoids), for a total of 108 transcriptomes and 72 proteomes (iPSCs were not profiled by proteomics). The number of samples analyzed and the pipeline for the analysis are shown in the **Supplementary Figures S5** and **S13**. Changes in cell populations and neuron morphology were examined by immunostaining, and neuron migration phenotype observed in the transcriptomic/proteomics experiment was validated using *in vitro* experiments. Finally, RhoA and KCTD13 levels were examined using Western Blot. For all quantifiable experiments, investigators were blinded for the analyses. Different numbers of samples and replicates were used for different experiments, as specified in the Figure legends and Supplementary Table S12. Raw data for all figures are provided in **Supplementary Table S13**.

### Tissue collection

Skin fibroblasts of three patients with 16p11.2 deletions (DEL) and three patients with 16p11.2 duplications (DUP) were obtained from the Simons Searchlight https://www.sfari.org/resource/simons-searchlight/; formerly Simons Variation in Individuals Project or Simons VIP). Patients were selected based on fibroblasts availability, head circumference, ASD diagnosis, and were gender and age matched (see detailed information about the patients in **Table S1**). De-identified patients tissue samples are distributed to Simons Investigators following approved IRB protocol to Simons Foundation through Columbia University Medical Center (PIs Drs. Gerald Fischbach and Wendy Chung). Collection and use for research of fibroblasts from three de-identified control individuals (CTRL) was approved by UCSD IRB. Skin fibroblasts were maintained in DMEM F-12 (Life Technologies) containing 10% fetal bovine serum.

### Generation and maintenance of iPSCs

To generate induced pluripotent stem cells (iPSCs), skin fibroblasts were infected with Sendai virus vectors containing coding sequences of human OCT4, SOX2, KLF4 and c-MYC (Cytotune reprogramming kit, Thermo Fisher). Four days post infection, fibroblasts were trypsinized to single cells, plated on the inactivated mouse embryonic fibroblast feeders, and cultured using human embryonic stem cell medium (Gibco). After 3–4 weeks, iPSC clones were manually picked and propagated clonally on feeders. After 8-10 passages, iPSCs were transferred to a feeder-free system and grown on matrigel-coated dishes (Corning) in mTeSR1 media (StemCell Technologies). The cells were passaged by manually picking colonies.

### Quality Control of generated iPSC clones

The generated iPSC clones were examined for genomic integrity by microarray genotyping. Parental fibroblasts and 8 iPSC clones for each patient were genotyped using BeadChip Illumina microarray platform. Copy Number Variants (CNVs) were called using PennCNV (v1.0.3) ^77^ with default parameters. Deletions or duplications were stitched as previously described ^78, 79^. Briefly, variants were joined if the gap between two CNVs of the same type was less than 50% of the number of markers within the larger CNV. This rule was applied recursively until no more CNVs could be stitched. Only CNVs of over 100 kbp in size were retained for the subsequent analysis. In addition, if over 50% of the CNV overlapped with the regions that can confound CNV calling (such as assembly gaps, segmental duplications, centromeres, and telomeres), they were omitted from the analyses. We also removed CNVs if the number of markers supporting the call was less than 8 and/or if the PennCNV confidence score was less than 20. After applying these filters, we confirmed the presence of 16p11.2 DELs or DUPs in all fibroblast and iPSC clones. We then sought to remove those iPSC clones, for which CNV burden was significantly greater than that of parental fibroblasts. To compare iPSC clones and fibroblasts, we defined CNV burden as a total sum of base pairs that are copy number variable (excluding 16p11.2 CNV). For each patient, we defined the mean CNV burden as the CNV burden in fibroblasts, and standard deviation as the burden in all iPSC clones from the same patient. We then compared CNV burden between fibroblasts and iPSC clones for each patient, and discarded clones with the Z-scores > 1.5 SD. Most clones passed these filtering steps. Two iPSC clones with CNV burden closest to the parental fibroblasts of each patient were used for subsequent experiments.

### Generation of cortical organoids

To generate cortical organoids from iPSCs, we used the protocol described in Trujillo et al. ^32^. Briefly, feeder-free iPSCs at passage 15 or later were fed daily with mTeSR1 for at least 7 days before differentiation. Colonies were dissociated using Accutase (Life Technologies) in PBS (1:1) for 10 minutes at 37°C and centrifuged for 3 minutes at 100x*g*. The cell pellet was resuspended in mTeSR1 supplemented with 10 μM SB431542 (SB, Tocris) and 1 μM Dorsomorphin (Dorso, R&D Systems). Approximately 5 × 10^6^ cells were transferred to each well of a 6-well plate and kept in suspension under rotation (95 rpm) in the presence of 5 μM ROCK inhibitor (StemCell Technologies) for 24 hours to form free-floating spheres. Then, the media was replaced with mTeSR1 for additional 48 hours. After 72 hours, Media1 [Neurobasal (Life Technologies) supplemented with Glutamax, 2% Gem21 NeuroPlex (Gemini Bio-Products), 1% N2 NeuroPlex (Gemini Bio-Products), 1% MEM nonessential amino acids (NEAA, Life Technologies), 1% penicillin/streptomycin (PS, LifeTechnologies), 10 μM SB and 1 μM Dorso] was used for maintenance for 7 days, with media changes every other day. Subsequently, Media1 was replaced with Media2 [Neurobasal with Glutamax, 2% Gem21 NeuroPlex, 1% NEAA and 1% PS] supplemented with 20 ng/mL FGF2 (Life Technologies) for additional 7 days. Then, Media2 was supplemented with both 20 ng/mL FGF2 and 20 ng/mL EGF (Life Technologies) and spheres were cultured for additional 7 days with media changes every other day. Next, organoids were transferred into Media3 [Media2 supplemented with 10 ng/mL BDNF, 10 ng/mL GDNF, 10 ng/mL NT-3 (all from Life Technologies), 200 μM L-ascorbic acid (Tocris) and 1 mM dibutyryl-cAMP (StemCell Technologies)] for another 7 days with media changes every other day. After 28 days, cortical organoids were maintained in Media2 for as long as needed, with media changes every 3-4 days. All organoids were generated, grown and used for all experiments in the same plate with one DEL, one DUP and one CTRL (called a “batch” thereafter) to reduce batch effect from genotypes.

### Mycoplasma testing

All iPSC and organoid cultures were routinely tested for mycoplasma by PCR. Media supernatants (with no antibiotics) were collected, centrifuged, and resuspended in saline buffer. Ten microliters of each sample were used for Mycosplama testing using commercially available LookOut Mycoplasma PCR Detection Kit (Sigma Aldrich) following manufacturer’s instructions.

### Organoid size analyses

The diameter of individual organoids was measured using ImageJ software. Size measurements for organoid batches (1 DEL, 1 DUP and 1 CTRL) followed normal distribution, as verified with Prism software (GraphPad).

### Immunofluorescence staining

Cortical organoids were fixed in 4% paraformaldehyde (PFA) overnight. Next morning they were washed in PBS, transferred to a 30% sucrose solution and kept at 4°C. After the 3D structures sink, they were transferred into Tissue-Tek OCT medium (Sakura). Subsequently, 20 μm thick sections were obtained using a cryostat. For immunostaining of iPSC clones, cells were grown directly on Matrigel-coated coverslips.

Slides containing organoid slices were air-dried and then washed with PBS to remove excess OCT. Permeabilization and blocking was performed with 4% Fetal Bovine Serum (FBS, Gibco), 0.1% Triton X-100 (Sigma Aldrich) diluted in PBS for one hour at room temperature. The slides were then incubated overnight at 4°C with primary antibodies diluted in solution containing 4% FBS and 0.1% Triton X-100. PBS was used to wash the primary antibodies and the slides were incubated with secondary antibodies in solution containing 4% FBS for 1h at room temperature. The following primary antibodies were used for immunostaining: NANOG (goat, 1:500, R&D Systems), Oct4 (mouse, 1:500, Abcam), Tra-1-60 (mouse, 1:500, Abcam), Lin28 (rabbit, 1:500, Abcam), PAX6 (mouse, 1:300; DSHB), NeuN (mouse, 1:500, Millipore), NeuN (rabbit, 1:500, Cell Signaling), MAP2 (chicken, 1:2000; Abcam), Ki67 (rabbit, 1:1000, Abcam), β-tubulin III (mouse, 1:500, Abcam), Synapsin I (rabbit, 1:500, Abcam), SOX2 (rabbit, 1:500, Abcam). Alexa Fluor Dyes (Abcam) were used at 1:1000 dilution as secondary antibodies. Nuclei were visualized with Hoechst 33258 (1:25000, Life Technologies). EdU was visualized using the Edu Staining Kit (Abcam) following manufacturer’s instructions. Slides were mounted using ProLong Gold antifade reagent (Invitrogen) and analyzed under a fluorescence microscope (Leica SP8). Image analysis was performed with ImageJ software. All cells expressing a particular marker were counted on sections and normalized to the total number of cells.

### Flow cytometry analysis

First, cortical organoids were dissociated to a single cell suspension. Then, cells were fixed for 15 min in ice-cold 1% PFS in PBS, washed twice with cold PBS, and incubated for 2h at room temperature with primary antibodies for specific cell markers (NeuN, SOX2, TBR2; Abcam) at 1:500 dilutions. Following a washing step with PBS, cells were incubated with 1:500 Alexa 488-conjugated antibodies (Abcam) for 20 min at room temperature. Analysis was performed on a flow cytometer (Accuri C6, BD Biosciences). Fifty thousand events were acquired for each sample with fluorescence measured in logarithmic scale. Background fluorescence was measured using cells labeled only with secondary antibody alone and used to set the gating parameters between positive and negative cell populations. Forward and side light-scatter gates were used to exclude cell aggregates and small debris. Data were analyzed using the FlowJo software and plotted in a histogram format. All histograms were smoothed by the software. Fluorescence gates were set below 2% of blank histogram and events corresponding to a fluorescence signal exceeding this percentage were considered as positive events.

### Cell migration assay

For the *in vitro* migration assay, intact organoids were seeded in Matrigel-coated 24-well plates (3-4 organoids per well), using Media2. Organoids were allowed to attach to the bottom of the plate for 24h, then media was replaced with fresh Media2 carefully not to disrupt organoids attachment. After 72h, immunostaining was performed. Images were taken using EVOS FL Cell Imaging System. Cell counting and image analysis were performed with ImageJ software.

For live-imaging, intact organoids were seeded in Matrigel-coated p35 glass-bottom culture dishes (Greiner). After 24h, pictures were taken every 15 min using a LEICA SP8 microscope. Videos were mounted using LEICA SP8 software.

### Organoids dissociation

Cortical organoids were dissociated into single-cell suspension using Accumax (Sigma Aldrich) for 30min at 37°C with rotation (95 rpm). Then, organoids were disaggregated using a 1000µl pipette tip, incubated for another 10min at 37°C in suspension with rotation (95 rpm), and centrifuged for 3 minutes at 100x*g*. The cell pellet was resuspended in Media2 containing 5 μM of ROCK inhibitor, filtered through a 100µm mesh (Gibco) and centrifuged again for 3 minutes at 100x*g*. To further remove un-dissociated organoid tissue, the procedure was repeated but with filtering through the 40µm mesh (Gibco). Cells from suspension were counted using a Bio-Rad TC10 Cell Counter.

### Boyden chamber migration assay

Approximately 3 × 10^5^ cells from the dissociated organoids were seeded on top of a Millicell Cell Culture 8µm Insert (Millipore) in 24-well plates. The bottom of the Cell Culture Insert was filled with 500µl of Media2 supplemented with 20 ng/mL of FGF2 and 10 ng/mL of BDNF as chemo-attractants. Cells were left to freely migrate for 24h, then washed with PBS and fixed with 4% PFA for immunostaining.

After immunostaining, images were taken using EVOS FL Cell Imaging System, visualizing all cells on the Cell Culture Inserts. Then, cells on the top of the Cell Culture Insert were removed using a cell scrapper. After three washes with PBS, only cells on the bottom of the insert were visualized. Cell counting was performed with ImageJ software.

### Pharmacological treatment of cortical organoids

For phenotype rescue experiments, organoids where grown in Rhosin-treated media. Rhosin (Tocris) was added to the media during differentiation stage starting from day 6 (Rhosin was first added to second Media1 to the final concentration of 1 µM). The same amount of Rhosin was added during all subsequent media changes. The organoids were grown to 1 month, at which cell migration assays were carried out. An equivalent amount of vehicle (0.1% Dimethylsulfoxide, DMSO) was added to grow untreated CTRL, DEL and DUP organoids.

### Neuronal morphology analysis

Cortical organoids were dissociated and approximately 3 × 10^5^ cells per well were seeded on a 24-well plate coated with poly-ornithine (Sigma Aldrich) and Laminin (Invitrogen). Media2 was changed after 24h to remove ROCK inhibitor, and second media change was performed after 3 days. Cells were fixed seven days after seeding for immunostaining. Images were taken with EVOS FL Cell Imaging System and analyzed with ImageJ software. For soma area calculation, the perimeter of the MAP2-positive cell body was manually outlined and measured. For total dendrite length, each dendrite or its branch was traced separately, and the dendrite length was calculated by adding individual lengths for every neuron.

### Synaptic puncta quantification

Three channel z-stack images of organoid slices were taken using an oil-inverted 60x objective. Then, an average projection image of each stack was generated. At least six randomly selected image fields for each genotype from two different batches were used for quantification of number of synapses with Synapsin I staining. Only puncta overlapping MAP2-positive processes were scored. The number of Synapsin I puncta was quantified using a plug-in Puncta Analyzer from the Fiji analysis software platform ^80^.

### Western Blot

Cortical organoids from a quarter to a half of a well were washed once with ice cold PBS (w/o Ca^2+^/Mg^2+^). Proteins were extracted using lysis buffer (20mM Tris, pH 7.4, 140mM NaCl, 10% glycerol, 2mM EDTA, 1mM EGTA, and 1% Triton X-100) supplemented with EDTA-free Complete protease inhibitor cocktail (Roche) and Phosphatase Inhibitor cocktail (Sigma Aldrich). The suspension was centrifuged at 16,000x*g* at 4°C for 30min, and supernatants were collected. Protein concentration was quantified by a modified Lowry assay (DC protein assay; Bio-Rad). Cell lysates were resolved by SDS-PAGE and transferred onto PVDF Immobilon-P membranes (Millipore). After blocking with 1X TBS, 0.1% Tween-20 containing 5% nonfat dry milk for 1h at room temperature, membranes were first probed with primary antibodies, and then after 1h of incubation with corresponding peroxidase-conjugated secondary antibody (Abcam). Membranes were developed using the EZ-ECL chemiluminescence detection kit (Denville Scientific). The following primary antibodies were used: anti-KCTD13 (1:500; Atlas Antibodies), anti-RhoA (1:1000; Cell Signaling), and anti-β-actin (1:5000; Sigma Aldrich) as a loading control. Quantification was performed by densitometry with ImageJ software.

### RNA isolation for RNA-Seq and qPCR

Total RNA was extracted from undifferentiated iPSCs or cortical organoids at 1 month and 3 months of differentiation. Two clones from each patient were used for RNA isolation, for each time point analyzed (**fig. S2**).

Total RNA was extracted using the QIAGEN RNAeasy isolation kit (QIAGEN) following manufacturer’s instructions. RNA sequencing was performed using the same input amount of total RNA for each sample. RNA samples were ribodepleted using Ribo-Zero rRNA Removal Kit (Illumina) and library preparation was performed using the TrueSeq Stranded Total RNA kit for Illumina Sequencing according to the manufacturer’s instructions. Paired-end RNA sequencing (2×150bp) was performed on an Illumina HiSeq4000 to an average depth of 40M reads per sample.

For qPCR experiments, cDNA was synthesized, starting from 100ng of total RNA with the SuperScript III First-Strand Synthesis kit and random hexamers (Invitrogen). qPCR was performed using the CFX96 Touch™ Real-Time PCR Detection System (Bio Rad) using Power SYBR Green PCR Master Mix (Applied Biosystems). HPRT1 and β-actin were used as housekeeping genes for normalization. Fold change in expression was calculated using the ΔΔCt method.

### RNA-sequencing Data Processing Pipeline

All 108 FASTQ files (36 iPSC, 36 one month organoids and 36 three months organoids paired-end fastq) (**fig. S5**) were run through a unified RNA-Seq processing pipeline. Pipeline source code can be found on https://github.com/IakouchevaLab/16p11.2. All fastqs were trimmed for adapter sequence and low base call quality (Phred score < 30 at ends) using Cutadapt (v1.14). Trimmed reads were then aligned to the GRCH37.p13 (hg19) reference genome via STAR (2.5.3a) using comprehensive gene annotations from Gencode (v19) (**fig. S7A**). Gene-level quantifications were calculated using RSEM (v1.3). Quality control metrics were calculated using RNA-SeQC (v1.1.8), featureCounts (v1.6.), PicardTools (v2.12), and Samtools (v1.3) (**fig. S7B** and **Table S2**).

### RNA-Seq Quality Control and Normalization

RNA-Seq Quality Control and Normalization Expected counts were compiled from gene-level RSEM quantifications and imported into R for downstream analyses. Expressed genes were defined as genes with TPM > 0.5 in at least 80% of samples from each genotype (CTRL, DEL or DUP). Only expressed genes were included into the analysis. A total of 15,788; 13,348, and 13,723 expressed genes from iPSC, 1M old organoids and 3M old organoids, respectively, were used in the downstream analysis using the GENCODE V19 annotation gtf file. Outliers were defined as samples with standardized sample network connectivity Z scores <−2 ^81^, and were removed (**fig. S7C**). Highly variable genes between clones from the same individual were filtered out using the Variance Partition (v3.5) R package ^82^.

### Covariate Selection

We compiled a set of 197 RNA-Seq quality control metrics from the outputs of cutadapt, STAR, RNA-SeQC, featureCounts and PicardTools (CollectAlignmentSummaryMetrics, CollectInsertSizeMetrics, CollectRnaSeqMetrics, CollectGcBiasMetrics, MarkDuplicates) for each group of samples (iPSCs, 1M old organoids and 3M old organoids) (**Table S2**, **figs. S7A-B**). These measures were summarized by the top principal components, which explained the total variance of each group (**fig. S8A**). Batch effects and possible hidden confounding factors were detected using the Surrogate Variable Analysis (SVA) ^83^. Multivariate Adaptive Regression Splines (MARS) implemented in the earth package in R was used to determine which covariates to include in the final differential expression model (**fig. S8B**). The potential covariates included: genotype, run/batch, RIN, clone, seqPCs and SVs (**fig. S8C**). These covariates were inputted into the earth model along with gene expression data (limma voom normalized, centered, and scaled). The model was run using linear predictors and otherwise default parameters. MARS selected SV1 as a covariate for iPSC, SV1 to SV5 as covariates for 1M old organoids, and SV1 to SV6 as covariates for 3M old organoids (**fig. S8B**).

### Differential Gene Expression

Differential Gene Expression (DGE) analyses was performed using limma-voom with ‘duplicateCorrelation’ function to account for duplicate samples (clones and replicas) from the same individuals, and to avoid pseudo-replication in the analyses ^43^. Covariates were included as fixed effects in the model. The biomaRt ^84, 85^ package in R was used to extract gene names, gene biotypes and gene descriptions. Differential genes expression analyses was performed using all three datasets (CTRL, DEL and DUP) for all time points. The volcano plots for DEL *vs* CTRL and DUPs *vs* CTRL are shown in **fig. S27**, and differentially expressed genes from these datasets are listed in **Table S4**.

### Weighted gene co-expression network analysis on RNA-seq data

We used Weighted Gene Co-expression Network Analysis (WGCNA) ^44^ to define modules of co-expressed genes from RNA-seq data. All covariates except for genotype at the 16p11.2 locus were first regressed out from the expression datasets. The co-expression networks and modules were estimated using the blockwiseModules function with the following parameters: corType=bicorr; networkType=signed; pamRespectsDendro=F; mergeCutHeight=0.1. Some parameters were specific for each dataset. For iPSC data: power=14; deepSplit=0; minModuleSize=100. For 1M old organoid data: power=16; deepSplit=2; minModuleSize=50. For 3M old organoid data: power=19; deepSplit=2; minModuleSize=70. Module eigengene/genotype associations were calculated using a linear mixed-effects model, using a random effect of individual, to account for multiple clones and replicas derived from the same subject. Significance p-values were FDR-corrected to account for multiple comparisons. Genes within each module were prioritized based on their module membership (kME), defined as correlation to the module eigengene. For selected modules, the top hub genes were shown. Module preservation was tested using the modulePreservation function from the WGCNA package in R.

### Enrichment analysis of GO functions and literature curated gene sets

Enrichment for Gene Ontology (GO; Biological Process and Molecular Function) was performed using gProfileR R package ^86^. Background was restricted to the expressed set of genes by group (iPSC, 1M old organoids and 3M old organoids). An ordered query was used, ranking genes by FDR-corrected p-value for DGE analyses and by kME for WGCNA analyses.

Enrichment analyses were also performed using several established, hypothesis-driven gene sets including syndromic and highly ranked (1 and 2) genes from SFARI Gene database (https://gene.sfari.org/database/gene-scoring/); pre- and post-synaptic genes from SynaptomeDB ^87^; genes with loss-of-function intolerance (pLI) > 0.99 as reported by the Exome Aggregation Consortium ^88^; highly constrained genes ^89^; FMRP targets ^90^ and CHD8 targets ^91^. Statistical enrichment analyses were performed using permutation test. One thousand simulated lists with similar number of genes, gene length distribution and GC-content distribution as the target gene list were generated, and the overlaps between each of the simulated list and the hypothesis-driven gene sets were calculated to form the null distribution. Significance p-value was calculated by comparing the actual overlap between target list and hypothesis-driven gene sets to the null distribution. All results were FDR-corrected for multiple comparisons.

### Cell type enrichment analysis

Cell-type enrichment analysis for each co-expression module was performed using the Expression Weighted Cell Type Enrichment (EWCE) package in R ^92^. Cell type-specific gene expression data was obtained from single cell sequencing (scRNA-seq) studies of human fetal neocortex ^49^. The specificity metric of each gene for each cell type was computed as described^92^. “Neuron” cell type includes a union of ExcNeu (excitatory neurons) and IntNeu (interneurons). Enrichment was evaluated using bootstrapping. Z-score was estimated by the distance of the mean expression of the target gene set from the mean expression of bootstrapping replicates. P-values were corrected for multiple comparisons using FDR.

### CoNTExT analyses

Regional and temporal identify of organoids was assessed using CoNTExT ^36^ (https://context.semel.ucla.edu/).

### Sample preparation, protein identification and quantification by TMT-Mass Spectrometry

TMT mass-spectrometry experiments were performed on the organoid samples from the same well as those used for RNA-seq, by splitting the content of each well into two approximately equal amounts (**fig. S13**). Organoids were lysed in 100 mM TEAB with 1% SDS, protease inhibitor cocktails (Sigma) and PhosSTOP (Sigma) by 2-3 times of brief probe sonication and then centrifuged at 18,000x*g* for 15 min at 4°C. Supernatants were reduced (10 mM TCEP at 55°C for 20 min) and alkylated (50 mM chloroacetamide at room temperature in the dark for 20 min), and then MeOH/CHCl_3_ precipitation was performed. Pellets were dissolved by adding 6M urea in 50 mM TEAB, and then LysC/Tryp (Promega) was added by 1:25 (w/w) ratio to the peptides. After 3-4 h incubation at 37°C, reaction mixture was diluted with 50 mM TEAB for urea to be less than 1 M. After the o/n digestion, peptide concentration was estimated by colorimetric peptide BCA assay (Thermo), and the peptides were labelled with TMT 10-plex reagents (Thermo) for one hour, followed by 15 min quenching with hydroxylamine according to the manufacturer’s protocol. Equal amount of reaction mixtures for each channel were pooled together and dried using SpeedVac.

Since the total number of samples exceeded the maximum number of TMT channels, samples were divided into multiple sets (one replicate per each set). To compare and normalize different sets of TMT-labeled samples, pooled peptides were labeled with 131N and 131C as duplicates, and these samples were commonly included in all sets within each age (1M and 3M old organoids) set. A total of 100μg of peptides were fractionated using Pierce™ High pH reversed-phase peptide fractionation kit (Thermo) and then dried in SpeedVac. Dried peptides were dissolved with buffer A (5% acetonitrile, 0.1% formic acid), and half of each fraction was injected directly onto a 25cm, 100μm-ID column packed with BEH 1.7μm C18 resin (Waters). Samples were separated at a flow rate of 300 nL/min on nLC 1000 (Thermo). A gradient of 1– 25% B (80% acetonitrile, 0.1% formic acid) over 200min, an increase to 50% B over 120 min, an increase to 90% B over another 30min and held at 90% B for a final 10min of washing was used for 360min total run time. Column was re-equilibrated with 20μL of buffer A prior to the injection of sample. Peptides were eluted directly from the tip of the column and nanosprayed directly into the mass spectrometer Orbitrap Fusion by application of 2.8kV voltage at the back of the column.

Fusion was operated in a data-dependent mode. Full MS1 scans were collected in the Orbitrap at 120k resolution. The cycle time was set to 3s, and within this 3s the most abundant ions per scan were selected for CID MS/MS in the ion trap. MS3 analysis with multi-notch isolation (SPS3) ^93^ was utilized for detection of TMT reporter ions at 60k resolution. Monoisotopic precursor selection was enabled, and dynamic exclusion was used with exclusion duration of 10s. Tandem mass spectra were extracted from the raw files using RawConverter ^94^ with monoisotopic peak selection. The spectral files from all fractions were uploaded into one experiment on Integrated Proteomics Applications (IP2, Ver.6.0.5) pipeline. Proteins and peptides were searched using ProLuCID ^95^ and DTASelect 2.0 ^96^ on IP2 against the UniProt *H. sapiens* protein database with reversed decoy sequences (UniProt_Human_reviewed_05-05-2016_reversed.fasta). Precursor mass tolerance was set to 50.0ppm, and the search space allowed all fully-tryptic and half-tryptic peptide candidates without limit to internal missed cleavage and with a fixed modification of 57.02146 on cysteine and 229.1629 on N-terminus and lysine. Peptide candidates were filtered using DTASelect parameters of -p 2 (proteins with at least one peptide are identified) -y 1 (partial tryptic end is allowed) --pfp 0.01 (protein FDR < 1%) -DM 5 (highest mass error 5 ppm) -U (unique peptide only). Quantification was performed by Census ^97^ on IP2.

### Differential protein expression

Proteomics data was first summarized to peptide level by adding up the intensities of constituting spectra. Quantitation results from different TMT runs were merged and normalized using the pooled samples channel which was present in all runs. For each peptide, multiple measurements from the same subject were collapsed to one measurement by taking the median of all measurements. The data was then log2 transformed. Differential protein expression (DPE) was calculated by fitting a linear mixed-effects model for each protein, using the lme4 package in R ^98^. Genotype was included as fixed effect in the model. We included a random effect term for each peptide to account for the fact that different peptides from the same protein are not entirely independent. Significance p values were calculated using lmerTest package in R ^99^. Resulting P-values were FDR-corrected using the Benjamini-Hochberg method to control for multiple comparisons. The volcano plots for DEL *vs* CTRL and DUPs *vs* CTRL are shown in **fig. S28**, and differentially expressed proteins from these datasets are listed in **Table S8**.

### Weighted protein co-expression network analysis

Proteomics data was first summarized to protein level by adding up the intensities of constituting peptides. Quantitation results from different TMT runs were merged and normalized using the pooled samples channel which was present in all runs. The merged data was then log2 transformed. Outlier samples detection, highly variable proteins removal, surrogate variables calculation and covariates selection were subsequently performed using the same methods as described for RNA-seq data processing. All covariates except for genotype at the 16p11.2 locus were first regressed out from the expression datasets. Protein co-expression networks and modules were estimated using the blockwiseModules function with the following parameters: corType=bicorr; networkType=signed; pamRespectsDendro=F; mergeCutHeight=0.1. Some parameters were specific for each dataset. For 1M old organoid data: power=13; deepSplit=3; minModuleSize=40; and for 3M old organoid data: power=17; deepSplit=2; minModuleSize=10. Module eigengene/genotype associations were calculated as described for the RNA-seq WGCNA. Module preservation was tested using the modulePreservation function from the WGCNA package in R.

### Quantification and statistical analyses

The statistical analyses for above experiments were performed using Prism software (GraphPad). Statistical tests used and exact values of n are described in Figure legends. Significance was defined as *p* < 0.05(*), *p* < 0.01(**), or *p* < 0.001(***). Blinded measurements were performed for any comparison between control and 16p11.2 genotypes. The samples used for each type of experiments are shown in **Table S12**.

For WB quantification analysis, the confidence intervals (CIs) at 95%, 99% and 99.99% for the standard error of the ratios of mean was calculated from the WB intensities of DEL-CTRL pairs (n =6), DUP-CTRL pairs (n = 6) and DUP-DEL pairs (n=6) (**fig. S25**) using confidence interval calculator for a ratio method ^100^. The maximum possible P-values were inferred from the above CIs.

### Materials availability

New induced pluripotent stem cells (iPSCs) lines generated in this study will be deposited after publication to Rutgers repository by the Simons Foundation for Autism Research that funded this study.

## Acknowledgments

This work was supported by a grant to LMI and ARM from the Simons Foundation for Autism Research (#345469), and by grants from the National Institute of Mental Health to LMI and ARM (MH109885, MH108528), to LMI (MH105524, MH104766), to JRY and ARM (MH100175), and to JS (MH119746). We thank Gabriel Hoffman, Karen Messer, Minya Pu, and Ruifeng Chen for suggestions regarding data analyses. We thank Lucas Bazier, Nicholas Chew and Alexander Sun for help with image analysis; and Jiaye Chen for help with uploading transcriptomic data to GEO database. RNA-seq data was generated at the UC San Diego IGM Genomics Center, University of California San Diego (grant P30CA023100). The images were acquired at the UCSD School of Medicine Microscopy Shared Facility (grant NS047101).

## Author Contributions

L.M.I. and A.R.M. conceived the study. J.U., P.Z., P.M-L., N-K.Y., P.D.N., C.A.T., D.A., J.S, J.R.Y. III, A.R.M. and L.M.I. designed the experiments and analyses. J.U., N-K.Y., P.D.N., C.T., M.A., J.D., L.T., and S.R. performed experiments and analyses. P.Z., P.M-L., D.A., K.C., and A.B.P. performed computational data processing and analyses. J.U. and L.M.I. wrote the manuscript, with input from all co-authors. Supervision was performed by J.S., J.R.Y. III, A.R.M, and L.M.I.

## Ethics declarations

Dr. Muotri is a co-founder and has equity interest in TISMOO, a company dedicated to genetic analysis and human brain organogenesis, focusing on therapeutic applications customized for autism spectrum disorders and other neurological disorders origin genetics. The terms of this arrangement have been reviewed and approved by the University of California, San Diego in accordance with its conflict of interest policies.

## Supplementary information

### Supplementary Figures Titles and legends

**Figure S1. Immunohistochemical validation of iPSC’s pluripotency.** Representative image of patient-derived iPSCs immunostained with DAPI, NANOG and Lin28 (left), or DAPI, Tra-1-60 and Oct4 (right).

**Figure S2. Transcriptional validation of iPSCs pluripotency**. Six pluripotency markers were quantified in patient-derived fibroblasts and corresponding iPSCs by qRT-PCR. Graph shows the average of three biological replicates for each cell line. 16pA, 16pB and 16pC are lines derived from the 16p11.2 DEL patients; and 16pX, 16Y and 16pZ are lines derived from 16p11.2 DUP patients.

**Figure S3. CNV burden analysis of patient-derived iPSC clones**. Microarray genotyping of fibroblasts and iPSCs and CNV burden analyses using PennCNV is shown. The presence of 16p11.2 CNV in DELs and DUPs was confirmed in all fibroblast and iPSC clones, and 16p11.2 CNV was removed in subsequent burden analyses. Patients’ mean CNV burden is defined as the CNV burden in fibroblasts, and standard deviation as the burden in all iPSC clones from the same patient. CNV burden between fibroblasts and iPSC clones for each patient was compared. Only iPSC clones with a CNV burden score <1 SD *vs* patients fibroblasts are shown. Fibroblasts CNV burden score is shown for reference. Arrows point to the iPSC clones selected for cortical organoids production.

**Figure S4. Time course of the cell type markers.** Transcriptional analysis of three cell type markers (NANOG, PAX6 and MAP2) in 16p11.2 patient-derived iPSCs, 1M and 3M cortical organoids by qRT-PCR. Graph shows the average of all patient cell lines (*n*=6), with three biological replicates per patient.

**Figure S5. RNA-seq experimental design and data analysis workflow.** A total of 108 transcriptomes have been sequenced in this study. A rigorous quality control including principal component analyses, sample connectivity analyses, surrogate variable analyses and multivariate adaptive regression spline (MARS) for covariate selection has been performed. The Limma-voom model with duplicate correlation function has been applied to account for clone replicates derived from the same patient in order to avoid pseudo-replication in the differential gene expression analyses ^43^.

**Figure S6. Predicted laminar transitions for 1M and 3M organoids compared to fetal neocortex**. The Figure was produced using TMAP ^36^ and the transcriptome of germinal zones of six 13–16 PCW human fetal neocortices ^38^. Rank-rank hypergeometric overlap (RRHO) maps for CTRL organoids (n=12 datasets) from 3 patients, 2 clones per patient, 2 replicates per clone are shown, with CTRL iPSCs (n=12 datasets) used as a second time point. Each pixel represents the overlap between fetal brain and organoids transcriptome, color-coded according to the −log_10_ p-value of a hypergeometric test. On each map, the extent of shared upregulated genes is displayed in the bottom left corner, whereas shared downregulated genes are displayed in the top right corners. Ventricular zone (VZ), Inner Subventricular zone (ISVZ), Outer Subventricular zone (OSVZ) and Cortical Plate (CP) are shown.

**Figure S7. Initial quality control metrics for RNA-seq data. (A)** Sequencing metrics from STAR (2.5.3a) for each group of samples (iPSCs, 1M and 3M old organoids). **(B)** Sequencing metrics from PicardTools (v2.12) for each group of samples (iPSC, 1M and 3M organoids). **(C)** Sample outlier removal performed with WGCNA package in R for each group of samples (iPSCs, 1M and 3M old organoids) based on Z-scores of standardized network connectivity. Outliers were defined as samples with Z scores of <(−2).

**Figure S8. Quality control metrics for RNA-seq data. (A)** Correlation plots among the top seqPCs (PCs that summarize the RNA-Seq QC metrics), surrogate variables selected by MARS, and other potential covariates (Run, RIN, Individual, Lab, Clone, Replica, Genotype, Z score) for each group of samples (iPSCs, 1M and 3M old organoids) generated by corrplot package in R. The spearman correlation coefficients values correspond to the areas of the circles. The legend shows the spearman correlation coefficient rho values. **(B)** Covariates for each group of samples (iPSCs, 1M and 3M old organoids) selected by MARS (implemented in earth package in R). **(C)** First two principal components (PCs) of gene expression values, calculated using “prcomp” function in R, are shown before (left panel) and after (right panel) covariate correction for the iPSC, 1M and 3M organoids. Colors represent saples (iPSCs, 1M and 3M organoids), and shapes represent genotypes: CTRL (circles), DEL (triangles) and DUP (squares).

**Figure S9. Gene co-expression modules for iPSCs.** Upper panel: WGCNA cluster dendrogram for iPSCs. Bottom panel: Module eigengene-genotype association. Comparisons are made between each genotype (DEL or DUP) and control (CTRL) for each module. Rows are genotypes (relative to CTRL) and columns are modules. Number in each tile is a beta value from linear mixed effect model, and color of each tile indicates statistical significance (see **STAR Methods**). A total of 7 modules were significantly associated with DEL or DUP genotypes in iPSCs.

**Figure S10. Gene co-expression modules for 1M old organoids.** Upper panel: WGCNA cluster dendrogram for 1M old organoids. Bottom panel: Module eigengene-genotype association. Comparisons are made between each genotype (DEL or DUP) and control (CTRL) for each module. Rows are genotypes (relative to CTRL) and columns are modules. Number in each tile is a beta value from linear mixed effect model, and color of each tile indicates statistical significance (see **STAR Methods**). A total of 7 modules were significantly associated with DEL or DUP genotypes in 1M old organoids.

**Figure S11. Gene co-expression modules for 3M old organoids.** Upper panel: WGCNA cluster dendrogram for 3M old organoids. Bottom panel: Module eigengene-genotype association. Comparisons are made between each genotype (DEL or DUP) and control (CTRL) for each module. Rows are genotypes (relative to CTRL) and columns are modules. Number in each tile is a beta value from linear mixed effect model, and color of each tile indicates statistical significance (see **STAR Methods**). A total of 21 modules were significantly associated with DEL or DUP genotypes in 3M old organoids.

**Figure S12. 16p11.2 gene modules in iPSCs, 1M and 3M old organoids.** The significantly associated with genotype 16p11.2 module was detected in all datasets (*10purple* in iPSCs, *11greenyellow* in 1M organoids, and *16lightcyan* in 3M old organoids). Left column of each panel: module-trait association. Middle column of each panel: module eigengene expression for DEL, DUP and CTRL datasets. Dots correspond to each replicate derived from 3 patients (CTRL, DEL or DUP), 2 clones per patient and 2 replicates per clone. Some replicates were removed before the analyses during outlier detection procedure (see **STAR Methods**). Right column of each panel: top 20 hub genes (based on kME) from each module are shown.

**Figure S13. Proteomics experimental design and data analysis workflow.** A total of 72 proteomes have been processed in this study by LC-MS/MS with TMT 11-plex labeling. Protein Quantification was carried out by Census. A rigorous quality control including principal component analyses, sample connectivity analyses, surrogate variable analyses and multivariate adaptive regression spline (MARS) for covariate selection has been performed. Linear mixed effect model (LME) was implemented for differential protein expression analyses.

**Figure S14. Protein co-expression modules for 1M old organoids.** Upper panel: WPCNA cluster dendrogram for 1M old organoids. Bottom panel: Module eigengene-genotype association. Comparisons are made between each genotype (DEL or DUP) and control (CTRL) for each module. Rows are genotypes (relative to CTRL) and columns are modules. Number in each tile is a beta value from linear mixed effect model, and color of each tile indicates statistical significance (see **STAR Methods**). A total of 5 modules were significantly associated with DEL or DUP genotypes in 1M old organoids.

**Figure S15. Protein co-expression modules for 3M old organoids.** Upper panel: WPCNA cluster dendrogram for 3M old organoids. Bottom panel: Module eigengene-genotype association. Comparisons are made between each genotype (DEL or DUP) and control (CTRL) for each module. Rows are genotypes (relative to CTRL) and columns are modules. Number in each tile is a beta value from linear mixed effect model, and color of each tile indicates statistical significance (see **STAR Methods**). A total of 7 modules were significantly associated with DEL or DUP genotypes in 3M old organoids.

**Figure S16. Protein co-expression modules enrichment analyses.** Hierarchical clustering of protein co-expression modules by module eigengene is shown. Module-genotype associations (* FDR<0.1) are shown below each module. Module enrichment analyses against literature-curated gene lists with previous evidence for involvement in autism are shown at the bottom (* FDR<0.05). The lists include syndromic and highly ranked (1 and 2) genes from SFARI Gene database (https://gene.sfari.org/database/gene-scoring/); pre- and post-synaptic genes from SynaptomeDB ^87^; genes with probability of loss-of-function intolerance (pLI)>0.99 as reported by the Exome Aggregation Consortium ^88^; constrained genes ^89^; FMRP target genes ^90^, and CHD8 target genes ^91^.

**Figure S17.** Correlation of differentially expressed genes and proteins in 3M organoids. Genes/proteins within 16p11.2 CNV locus are colored in pink.

**Figure S18. Module preservation analyses between gene co-expression and protein co-expression modules.** Module preservation scores of gene co-expression modules (vs. corresponding protein co-expression modules) for 1M old organoids (upper panel) and 3M old organoids (bottom panel) are shown. Gene modules significantly associated with genotypes (DEL and DUP) are marked. Z-scores above 2 are considered to be conserved, and above 10 are highly conserved.

**Figure S19. Module preservation analyses for migration, neuronal/synaptic and 16p11.2 modules from 3M old organoids**. Module preservation scores of 16p11.2 (*16lightcyan*), neuronal/synaptic (*25orange*) and migration (*32violet*) gene co-expression modules (vs. iPSC and 1M transcriptomic, and 1M and 3M proteomic modules) for 3M old organoids are shown. Z-scores above 2 are considered to be conserved.

**Figure S20. Flow cytometry of cerebral organoids.** Representative images of histograms used for the flow cytometry analysis of dissociated cerebral organoids. Events with higher fluorescence than background histograms (within the area delimited by bars) were considered positive and quantified.

**Figure S21. Proliferation rate in 1M old organoids.** Upper panel: **r**epresentative images of 1M organoid slices immunostained with DAPI, Ki67 and Edu. Bottom panel: quantification of the percentage of positive cells for each marker in each genotype and cell cycle exit ratio. Symbols represent organoids from the same differentiation batch, where batch is defined as CTRL, DEL, DUP from one patient, one clone and one replica. Data is presented as mean ± SEM (*n=*2 patients per genotype, at least 4 organoids per patient). Significance was calculated using one-way ANOVA; \*\*\**p*<0.001, \*\**p*<0.01, \**p*<0.05. Significance above bars represents comparison against CTRL.

**Figure S22.** Wnt signaling genes downregulated in 16p11.2 DEL organoids. Heat plot represents TPM values of gene expression for Wnt signaling genes from *22darkgreen* module.

**Figure S23. Impaired migration in DEL and DUP organoids.** Upper panel: **r**epresentative images of neurons migrating out of Matrigel-attached organoids at the indicated time points after the start of time-lapse. Arrows mark individual neurons. Bottom panel: quantification of total distance traveled by individual neurons (mean±SEM; one-way ANOVA, ***p<0.001, *p<0.05; n=4 neurons per organoid, 3-4 organoids per genotype). Tracing of cell movement of individual representative neurons for each genotype is shown on the right. Each dot represents location of the neuron after 1h time period.

**Figure S24. Immunostaining of 1M old organoids with neuronal, developmental and intermediate filament markers.** Representative images of 1M old organoid attached in Matrigel, stained with DAPI, Sox2 and NeuN (left), DAPI and Nestin (right).

**Figure S25. Western Blots images of total and active RhoA.** Western Blot images of 1M organoids for KCTD13, actin as loading control, total RhoA, and active RhoA (RhoA-GTP). Organoids were grown in the batches of CTRL, DEL and DUP for each experiment. Six batches were grown for each, total RhoA and active RhoA experiments.

**Figure S26. Rhosin does not rescue neurite length phenotype.** Representative images of neurons from dissociated organoids 8 days after dissociation, immunostained with DAPI (blue) and MAP2 (green). Quantification of total neurite length for each genotype with or without Rhosin treatment is shown. Symbols represent organoids from the same differentiation batch, where batch is defined as CTRL, DEL, DUP from one patient, one clone and one replica. Data is presented as mean ± SD (n=2 patients per genotype, at least 15 neurons per patient). Significance is calculated using one-way ANOVA; ***p<0.001, **p<0.01, *p<0.05. Significance above bars represents comparison against CTRL.

**Figure S27. Differential gene expression (DEL *vs* CTRL and DUP *vs* CTRL) for iPSCs, 1M and 3M cortical organoids.** Volcano plots of differentially expressed genes in DEL *vs* CTRL (left column) and DUP *vs* CTRL (right column) for iPSCs, 1M and 3M old organoids. Genes within 16p11.2 CNV are colored in pink. Genes colored in orange are upregulated in DEL or DUP compared to CTRL; genes colored in blue are downregulated in DEL or DUP compared to CTRL.

**Figure S28. Differential protein expression (DEL *vs* CTRL and DUP *vs* CTRL) for 1M and 3M cortical organoids.** Volcano plots of differentially expressed proteins in DEL *vs* CTRL and DUP *vs* CTRL for 1M and 3M old organoids. Proteins within 16p11.2 CNV are colored in pink. Genes colored in orange are upregulated in DEL or DUP compared to CTRL; genes colored in blue are downregulated in DEL or DUP compared to CTRL. Gene Ontology enrichment analyses are shown as bar plots on the right. The contribution of up- or down-regulated proteins to specific GO terms are shown in blue and red, respectively.

### Supplementary Table titles and legends

**Supplementary Table 1.** 16p11.2 patient-derived fibroblast selection and patients’ clinical information.

**Supplementary Table 2.** RNA-seq parameters and quality control metrics from Cutadapt, STAR, Picard, and RNA-SeQ for iPSCs, 1M and 3M old organoids representing 108 transcriptomes.

**Supplementary Table 3.** Cortical organoids size analyses.

**Supplementary Table 4.** Differentially Expressed Genes (DEGs) in iPSCs, 1M and 3M cortical organoids.

**Supplementary Table 5.** Gene Ontology enrichment analysis of differentially expressed genes.

**Supplementary Table 6.** Module membership from gene co-expression (WGCNA) analysis in iPSCs, 1M and 3M cortical organoids.

**Supplementary Table 7.** Gene Ontology enrichment analyses of significantly genotype-associated gene co-expression modules from WGCNA.

**Supplementary Table 8.** Differentially Expressed Proteins (DEPs) in 1M and 3M cortical organoids detected by LC-MS/MS with TMT labeling.

**Supplementary Table 9.** Gene Ontology enrichment analysis of differentially expressed proteins.

**Supplementary Table 10.** Module membership from protein co-expression (WPCNA) analysis in 1M and 3M cortical organoids.

**Supplementary Table 11.** Gene Ontology enrichment analyses of significantly genotype-associated protein co-expression modules from WPCNA.

**Supplementary Table 12.** Summary of experiments by patients, clones and replicates.

**Supplementary Table 13.** Raw data for all experimental Figures.

