## Supplementary Figures for "Cortical Organoids Model Early Brain Development Disrupted by 16p11.2 Copy Number Variants in Autism"

### Supplementary Fig. 1

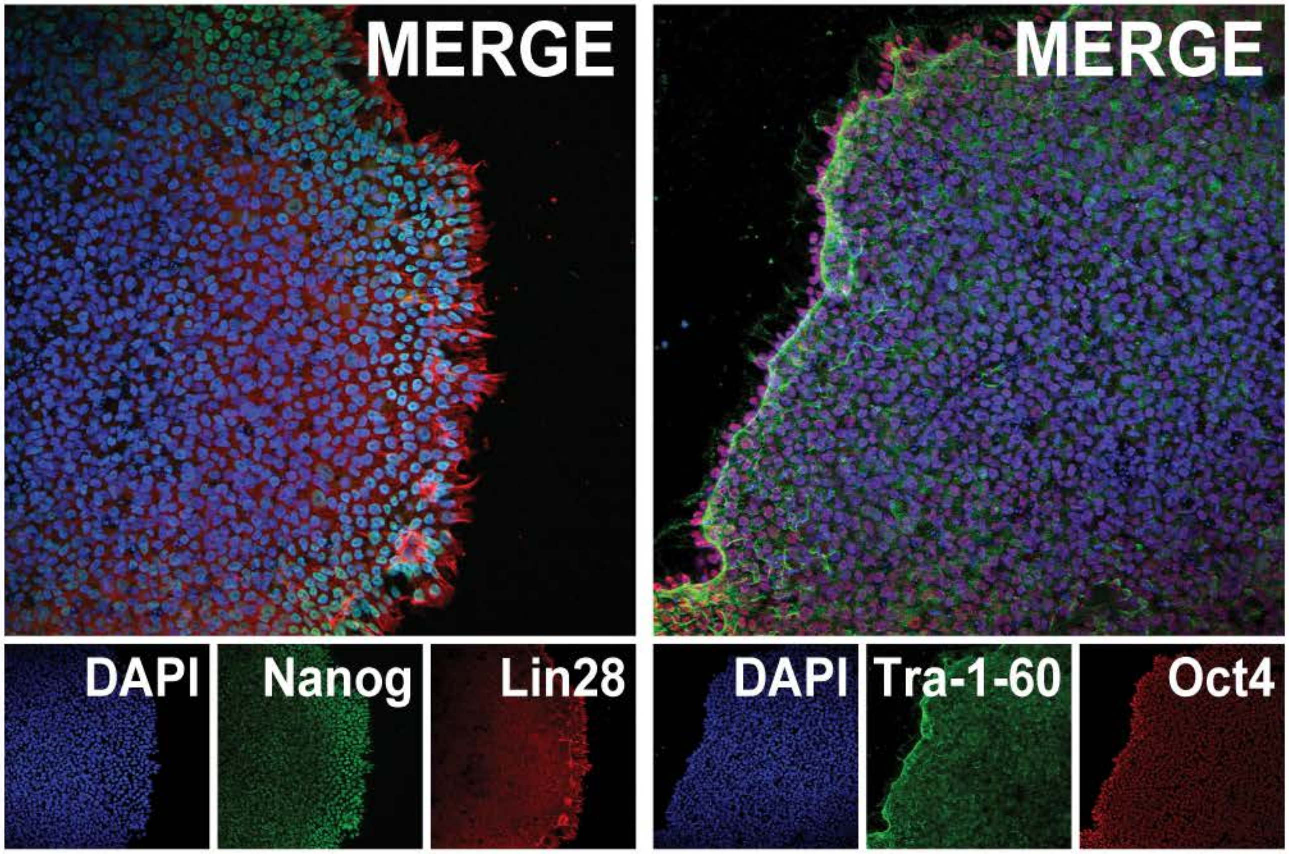


### Supplementary Fig. 2

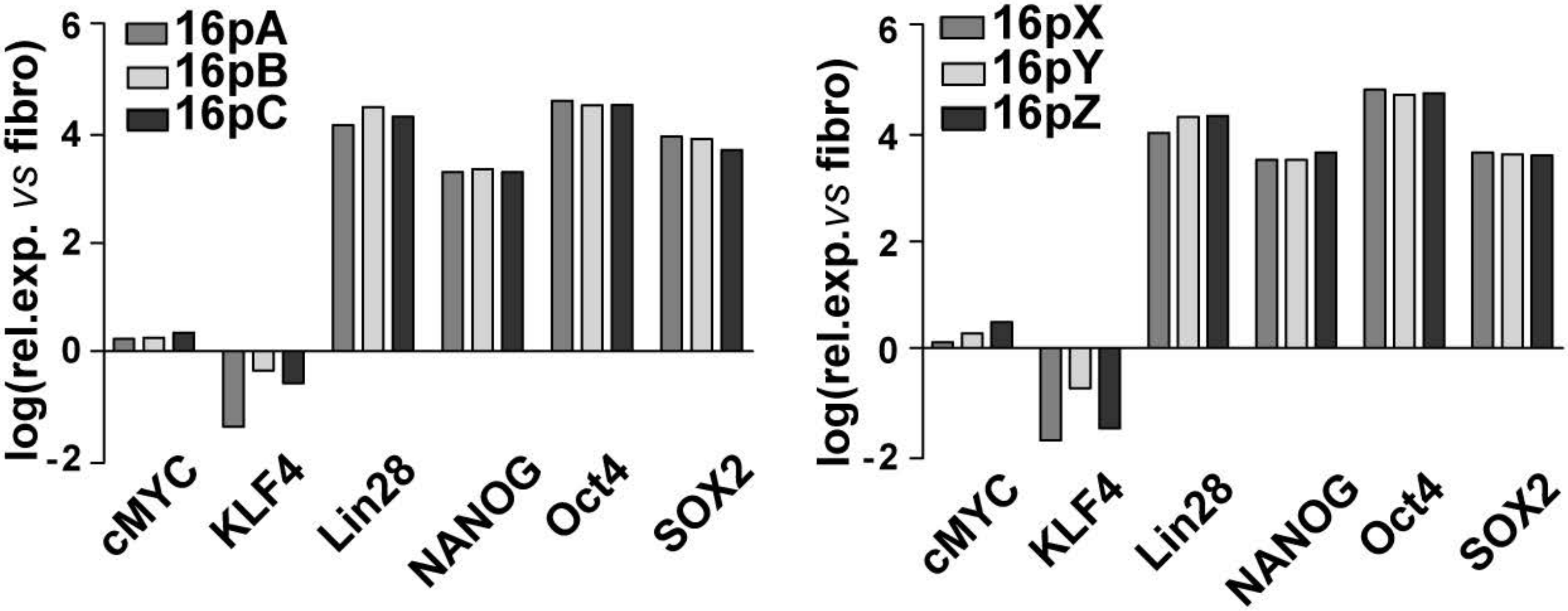

### Supplementary Fig. 3

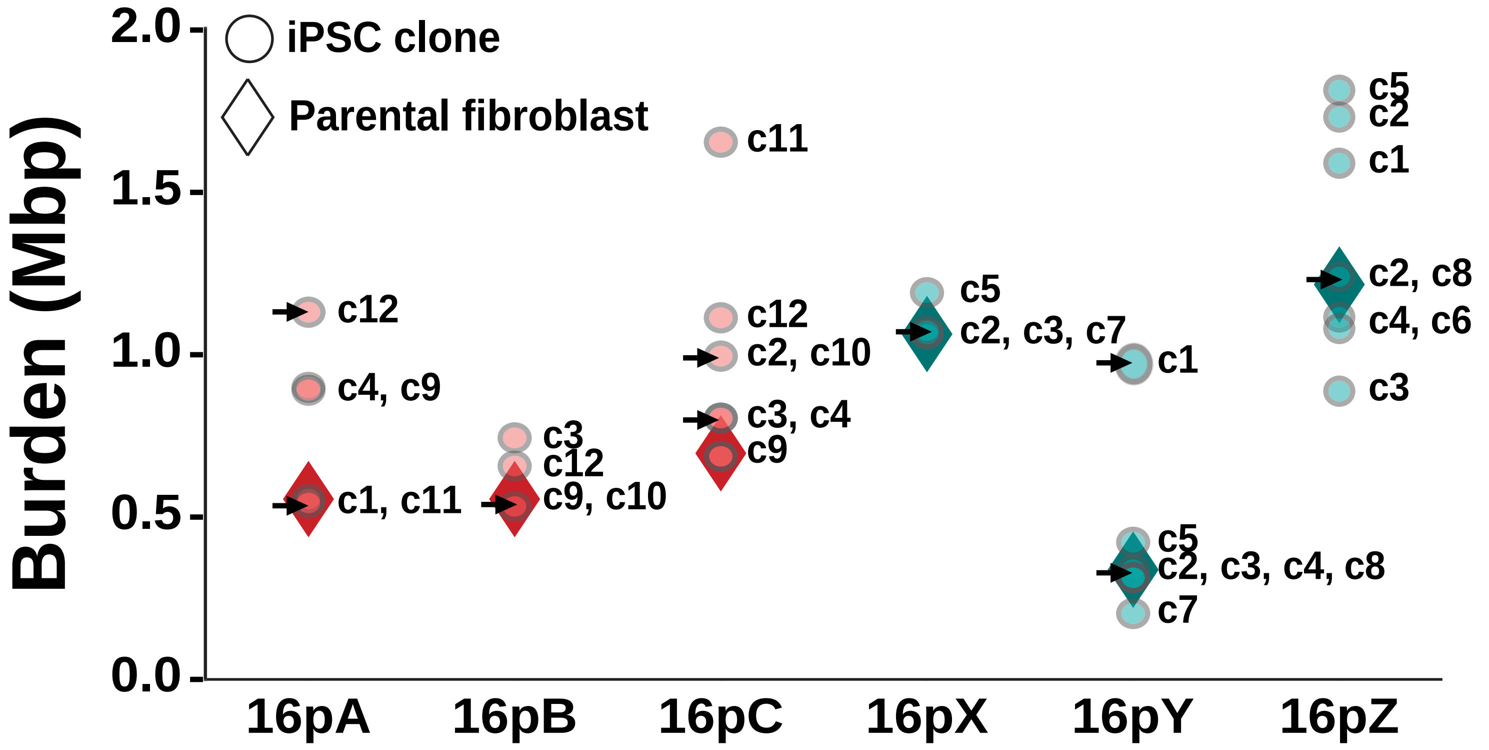


### Supplementary Fig. 4

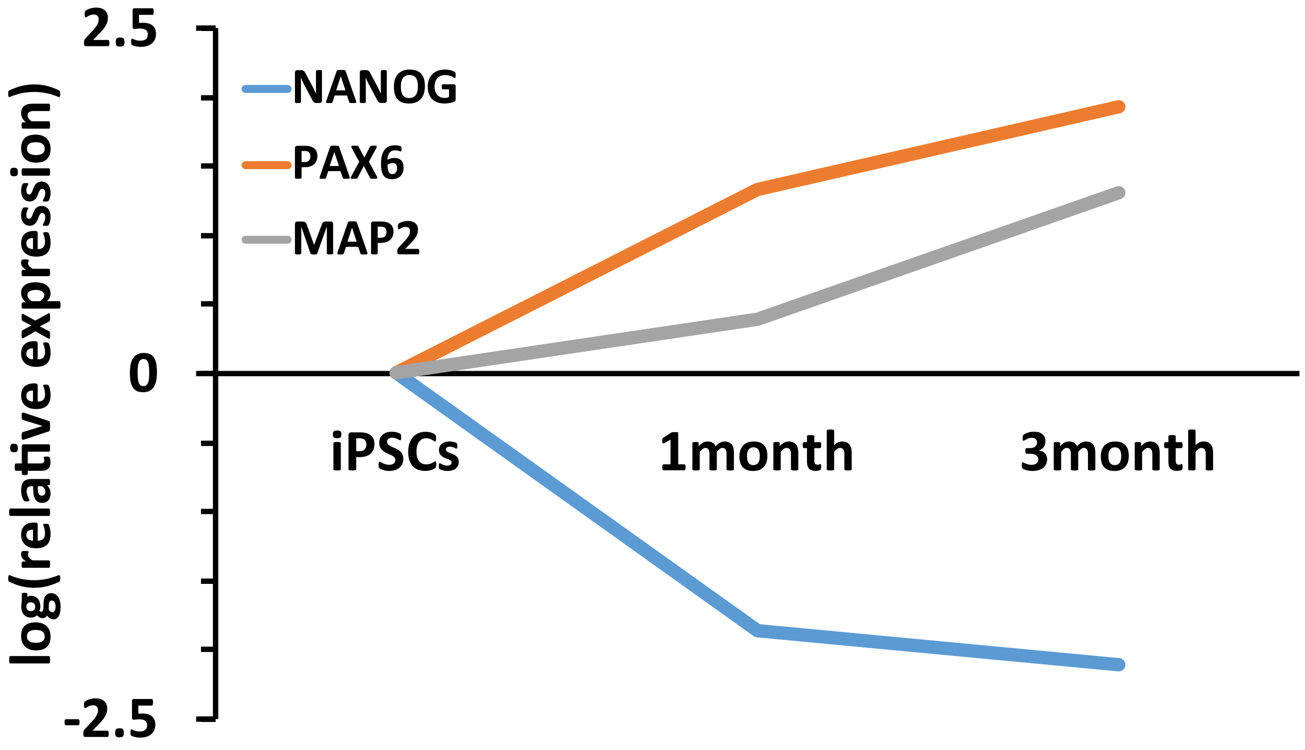

#### Supplementary Fig. 5

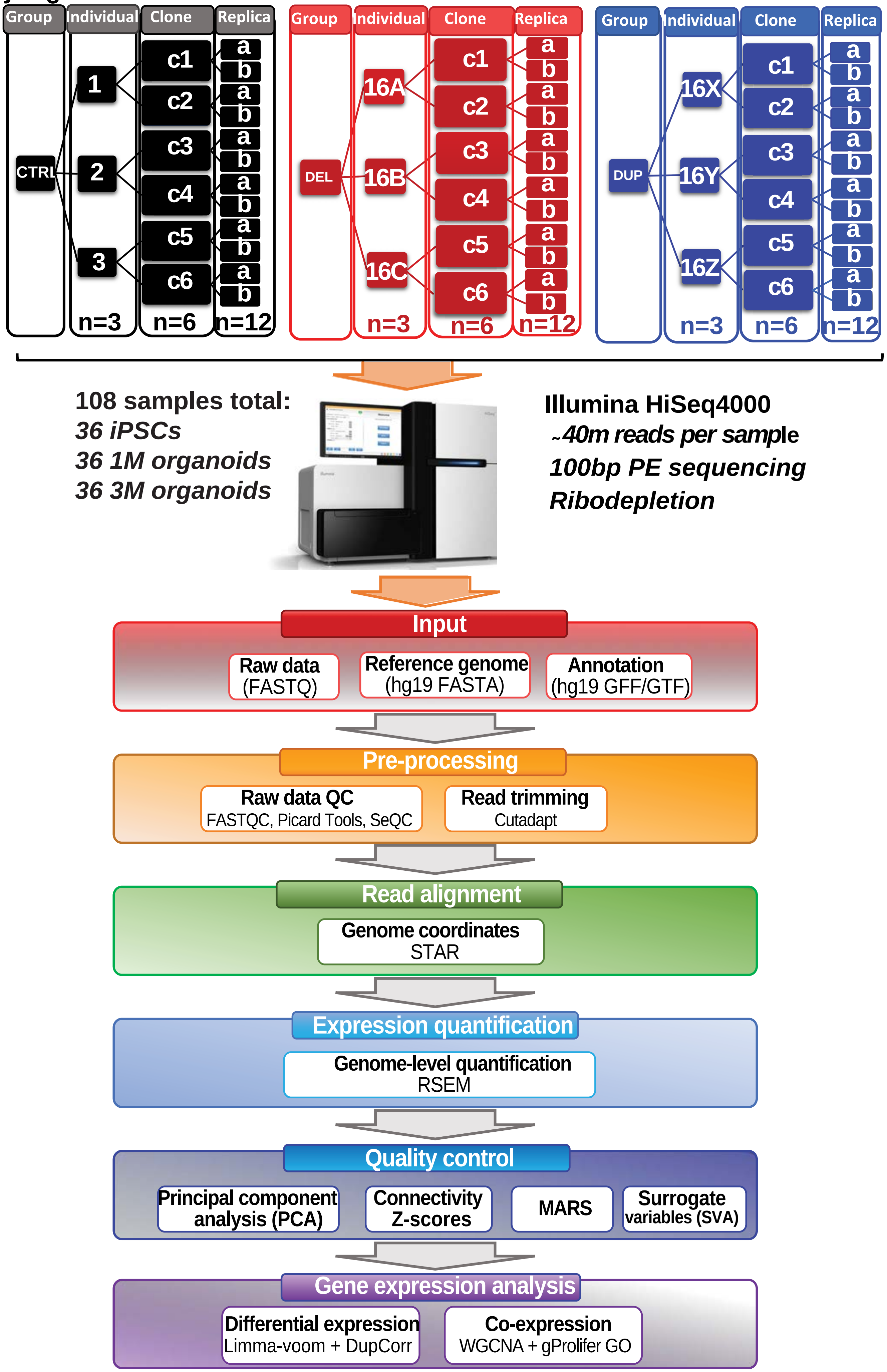

### Supplementary Fig. 6

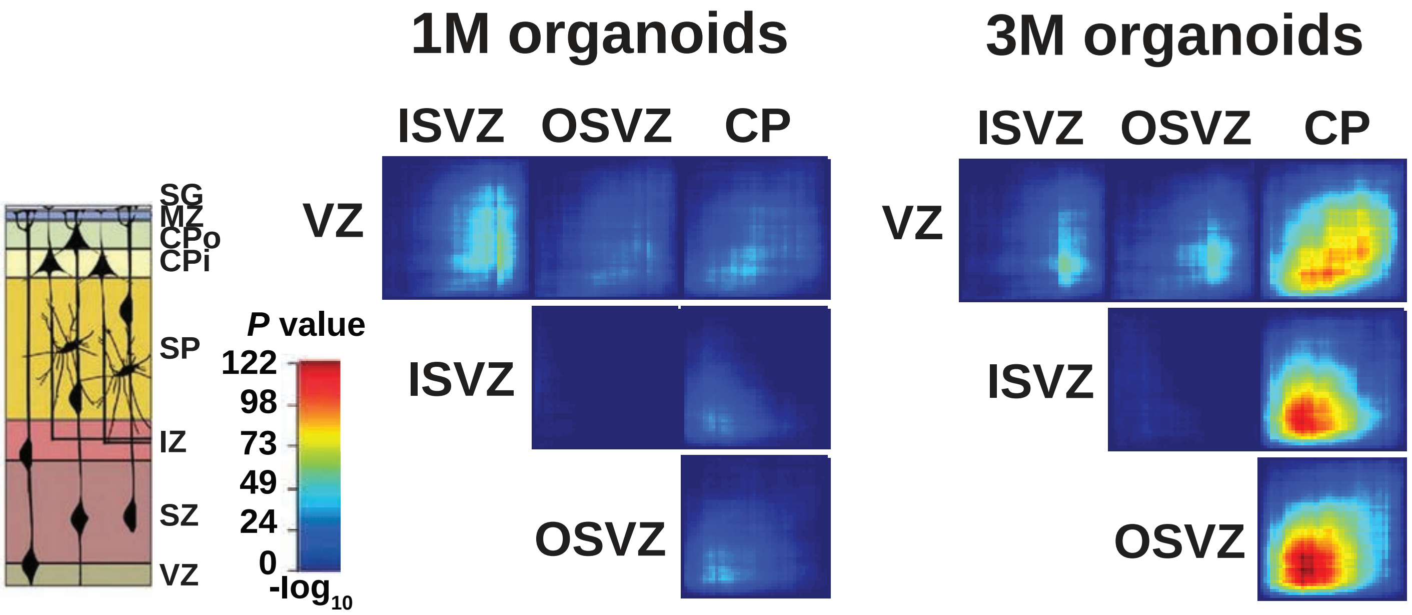

### Supplementary Fig. 7

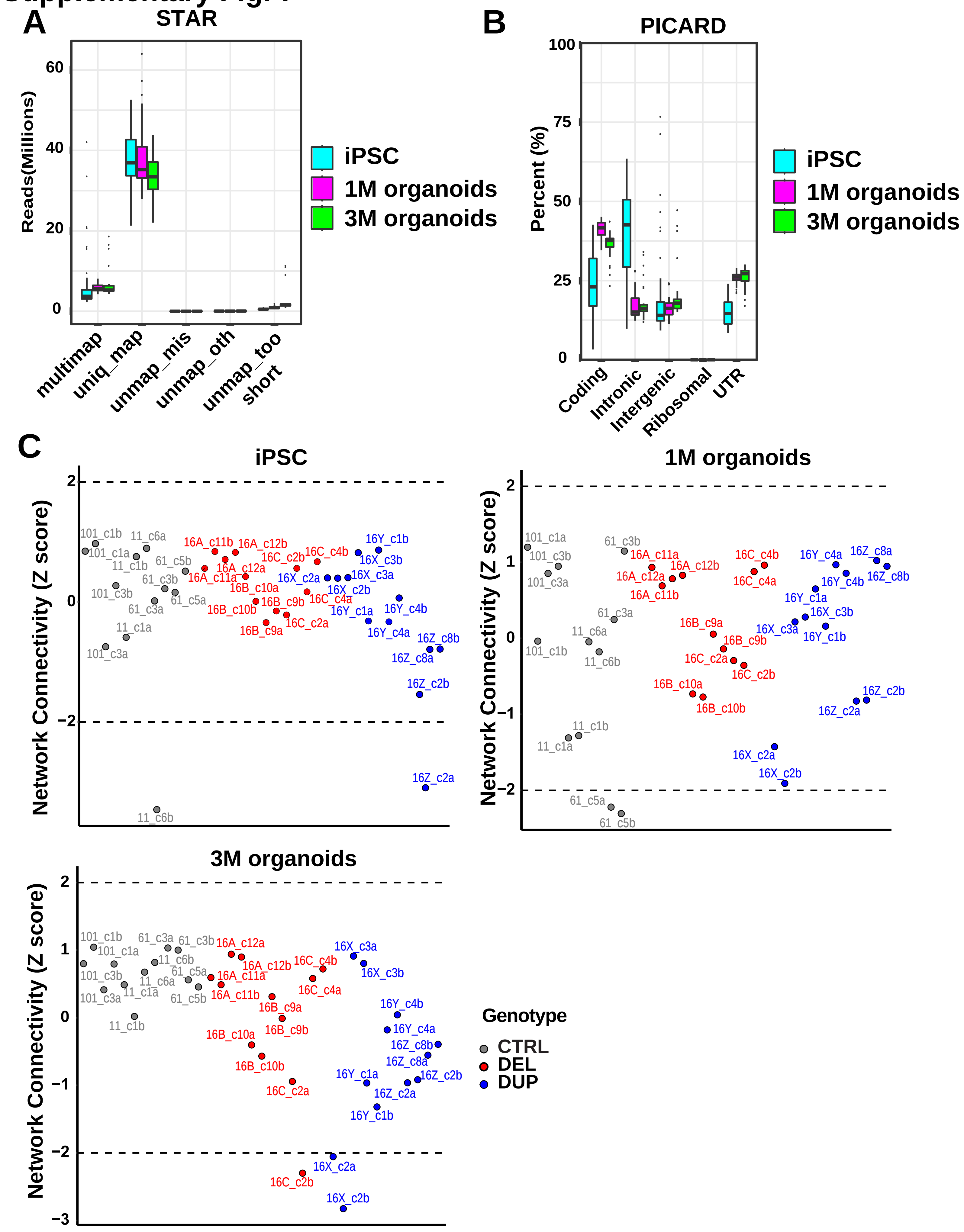

### Supplementary Fig. 8

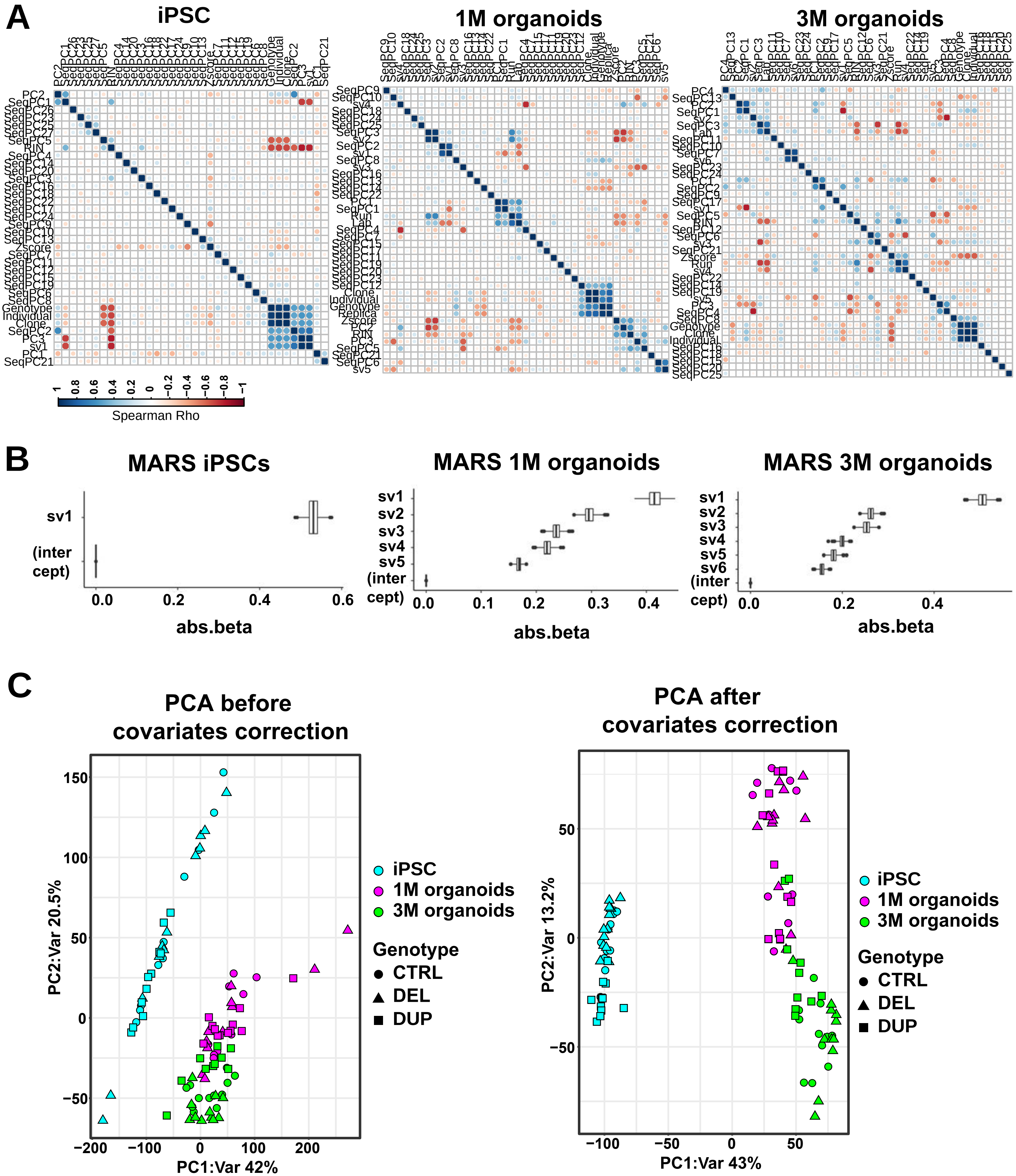

### Supplementary Fig. 9

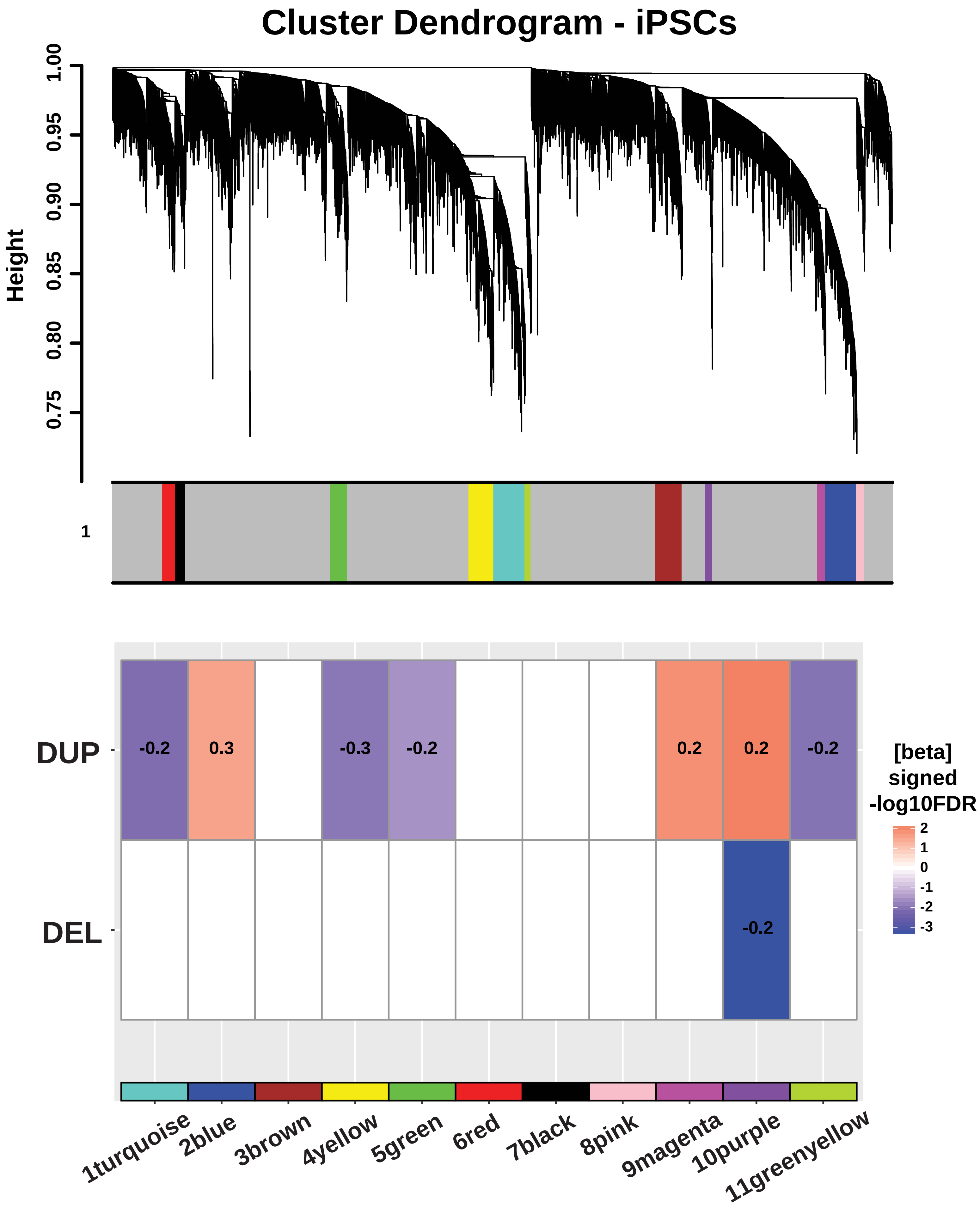

### Supplementary Fig. 10

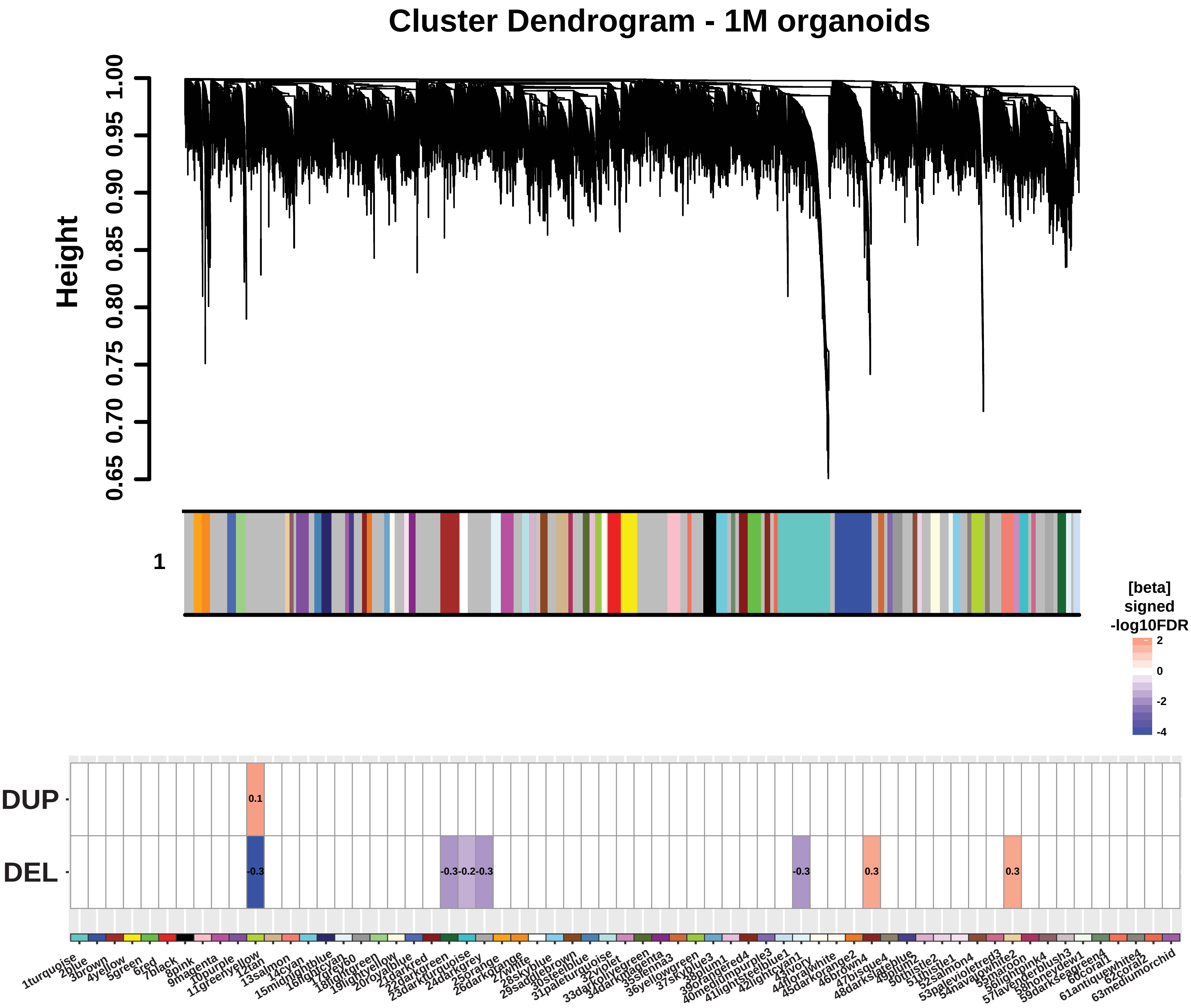

### Supplementary Fig. 11

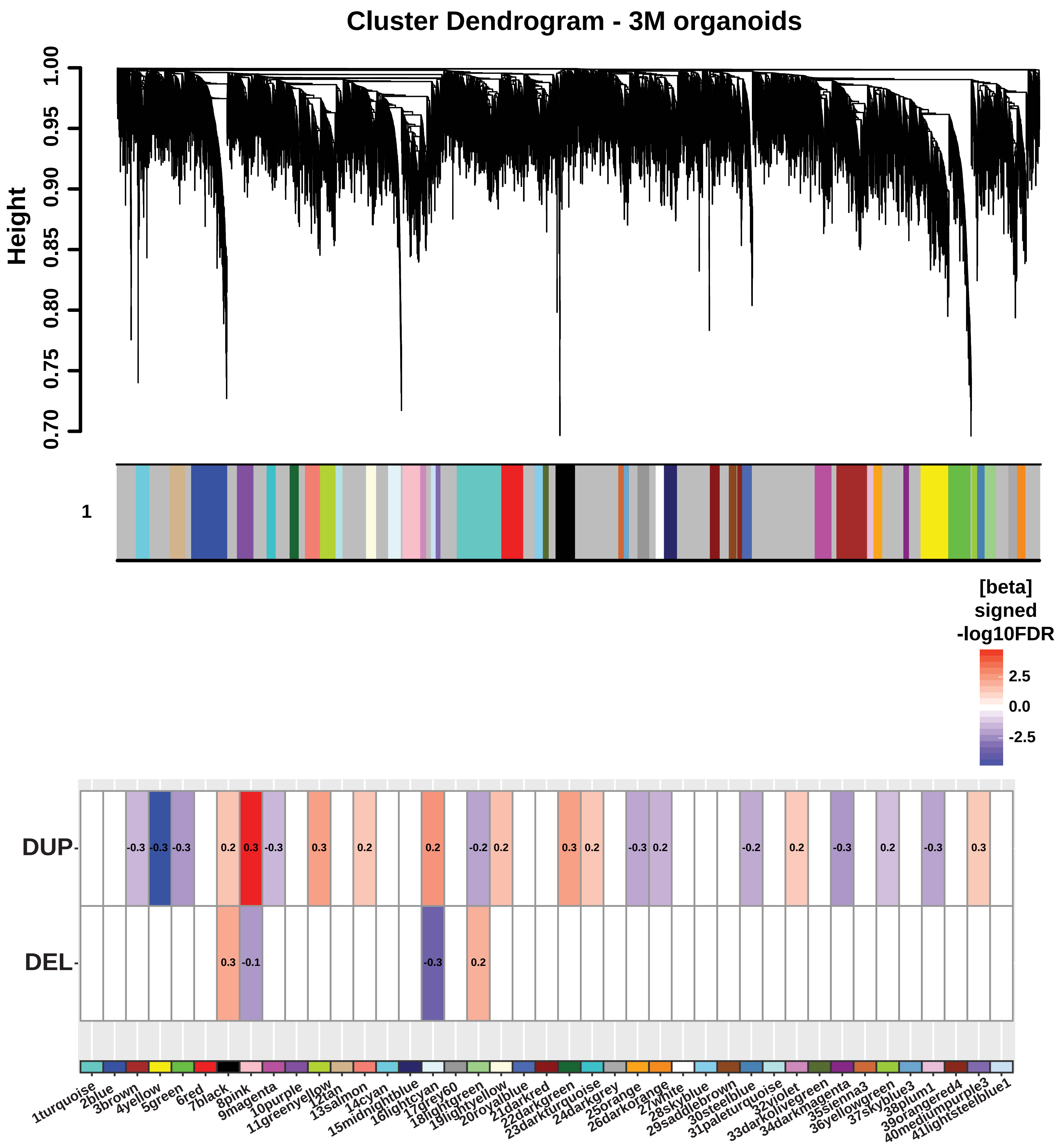

### Supplementary Fig. 12

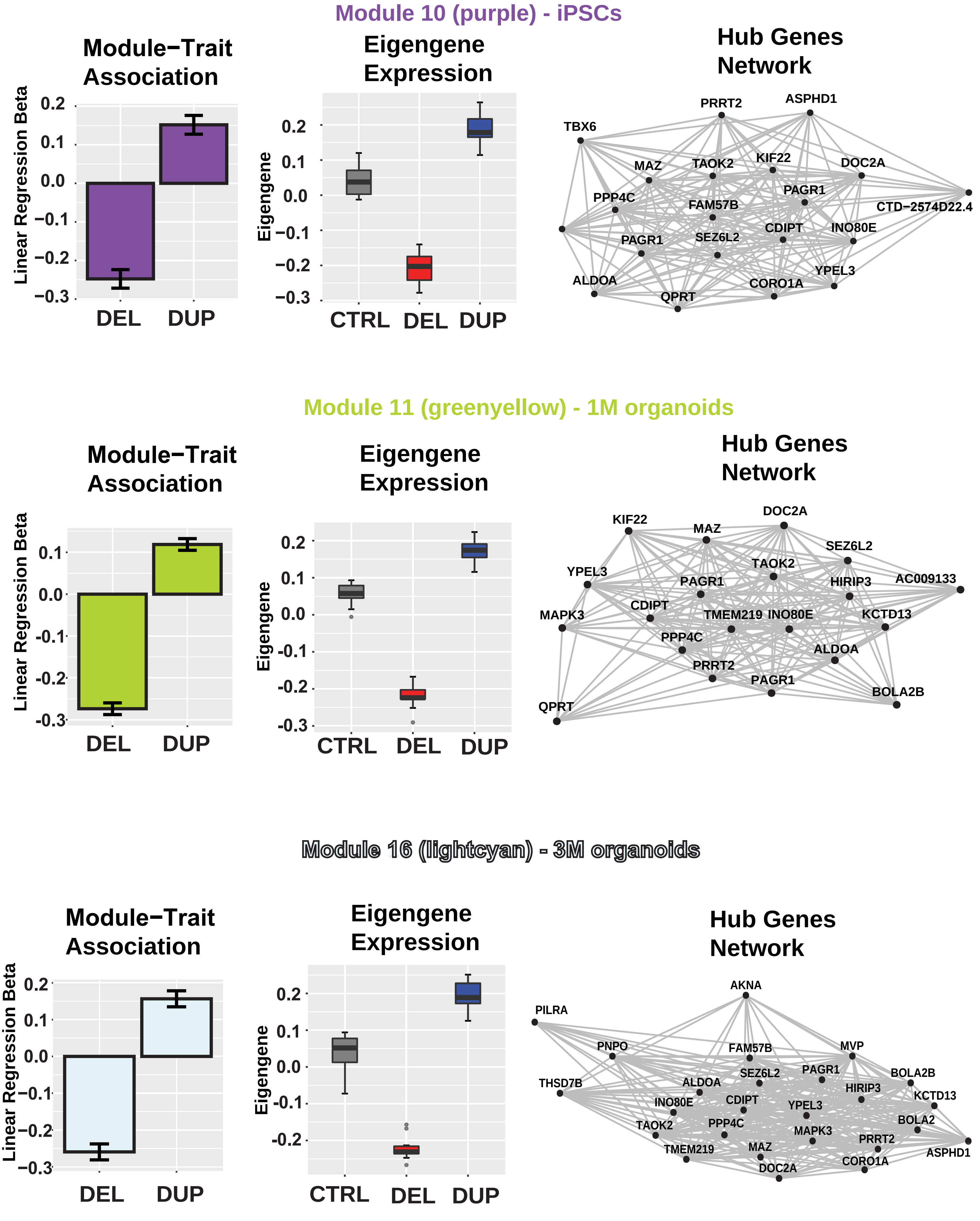

### Supplementary Fig. 13

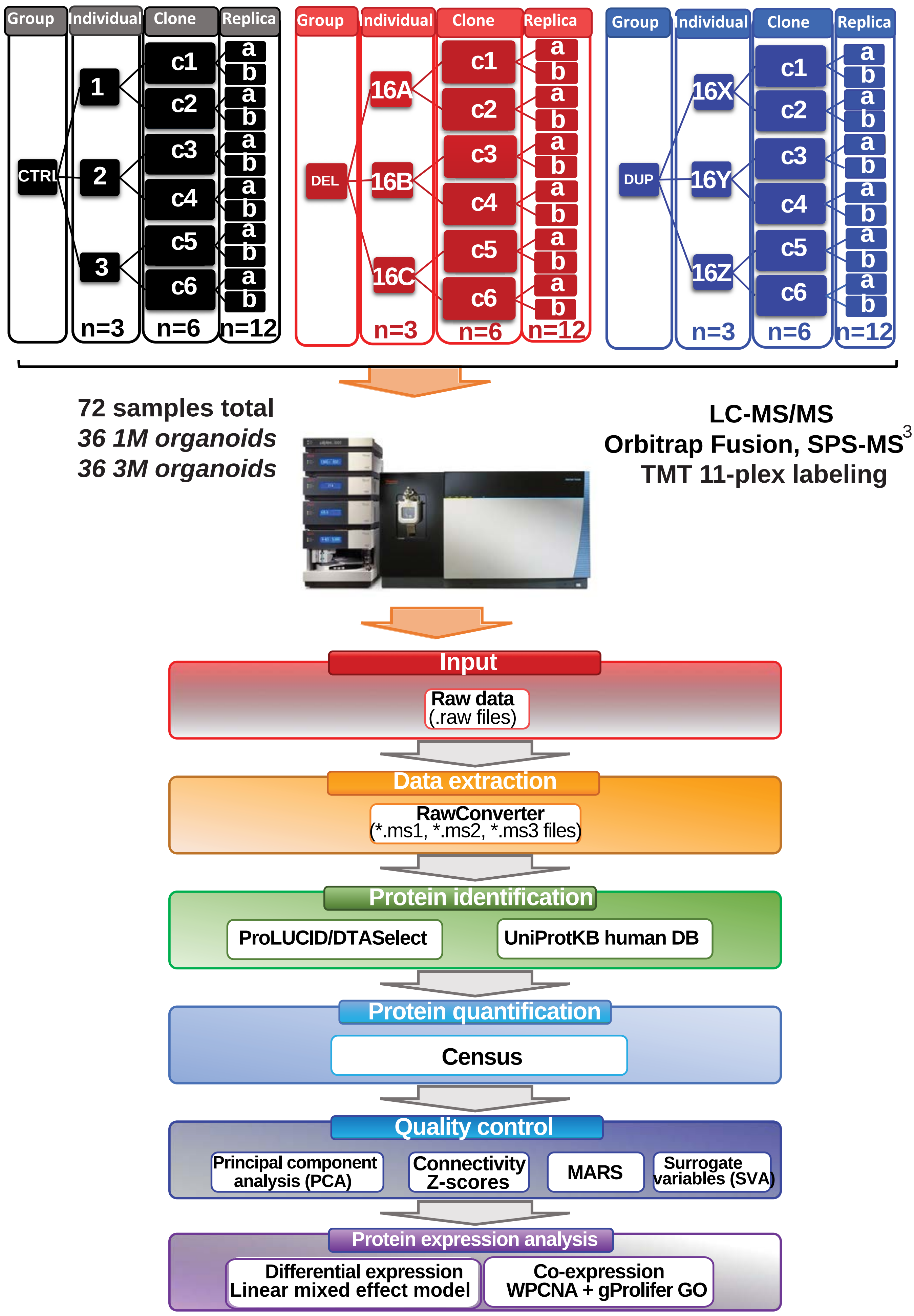

### Supplementary Fig. 14

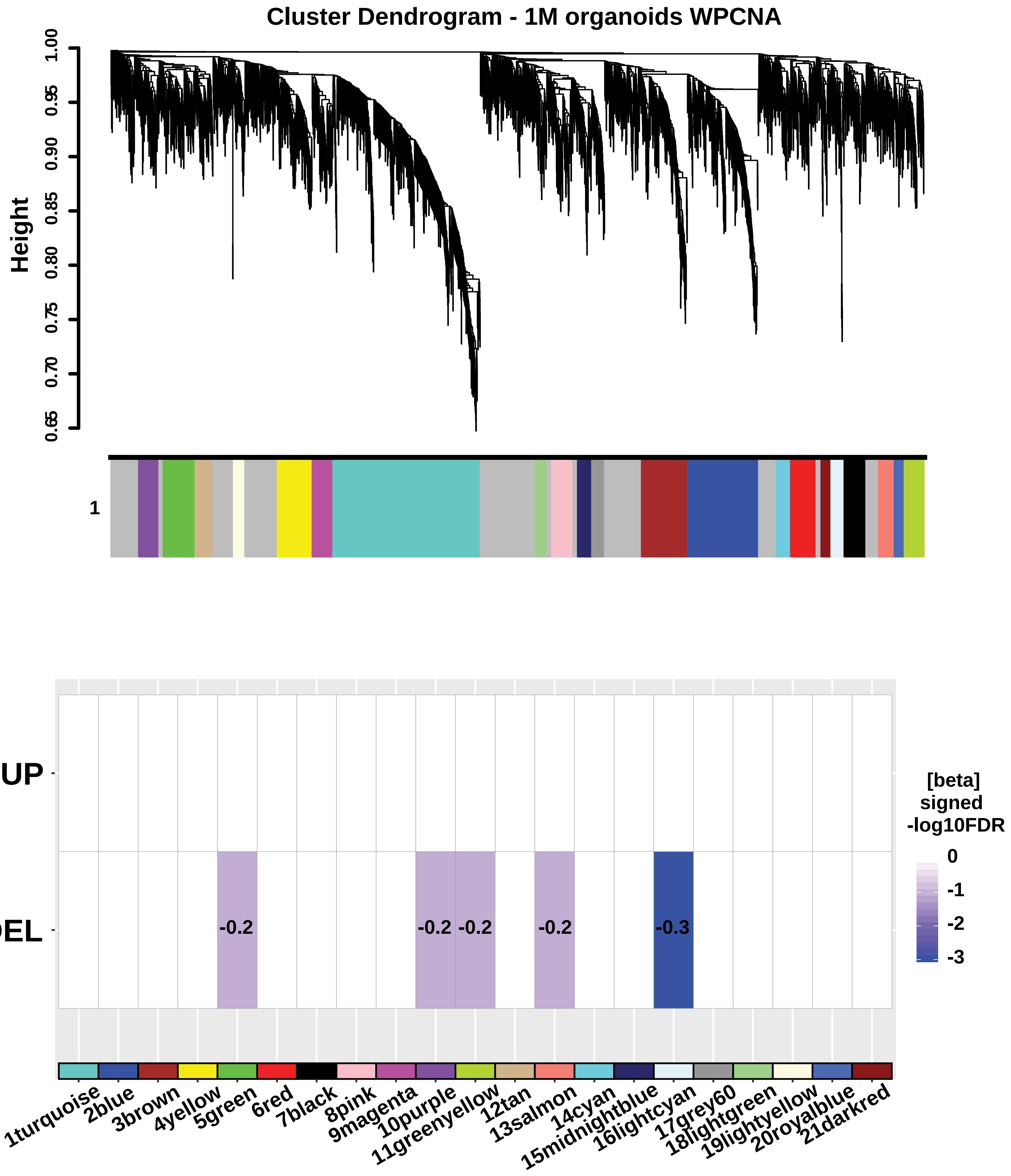

### Supplementary Fig. 15

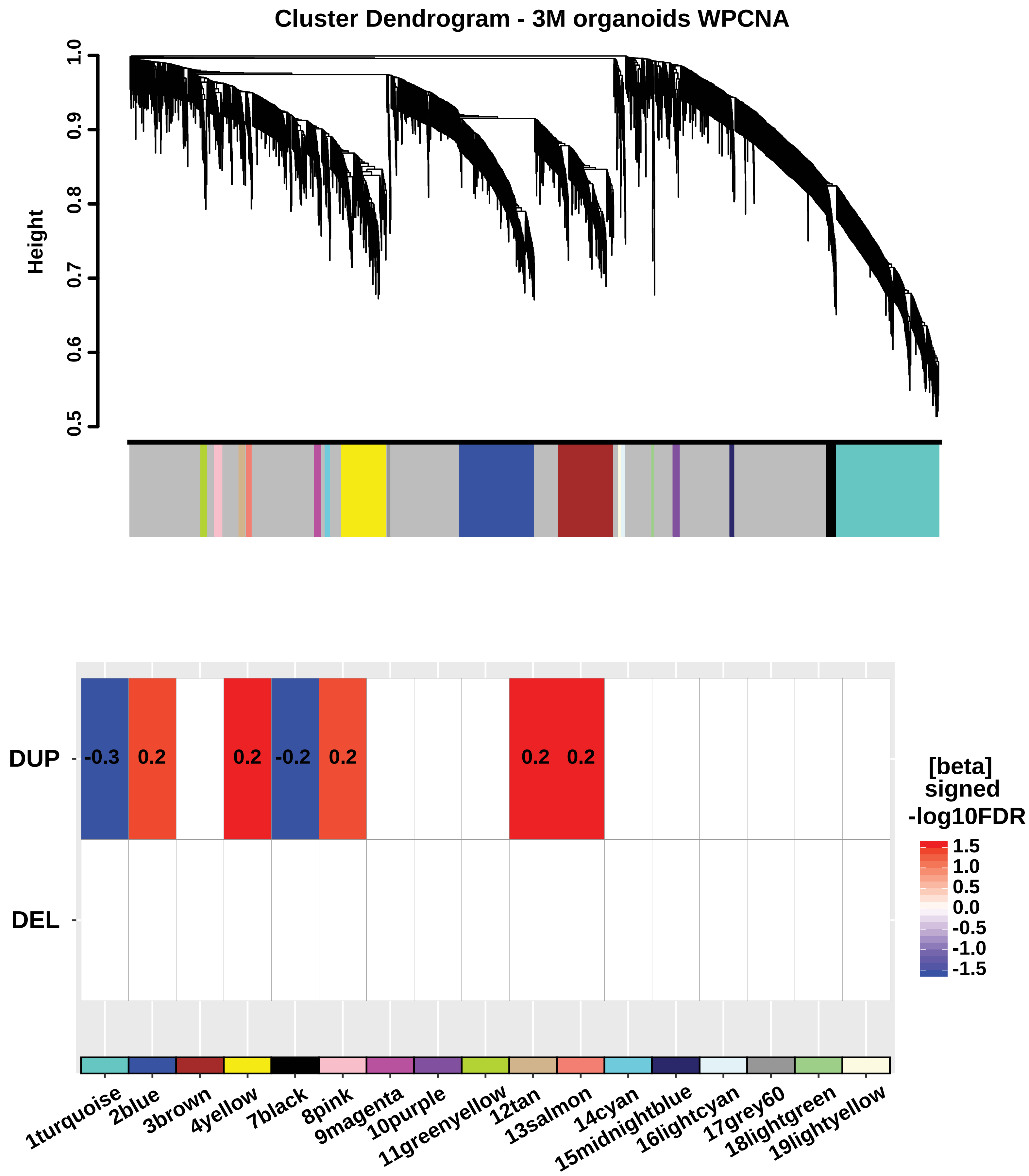

### Supplementary Fig. 16

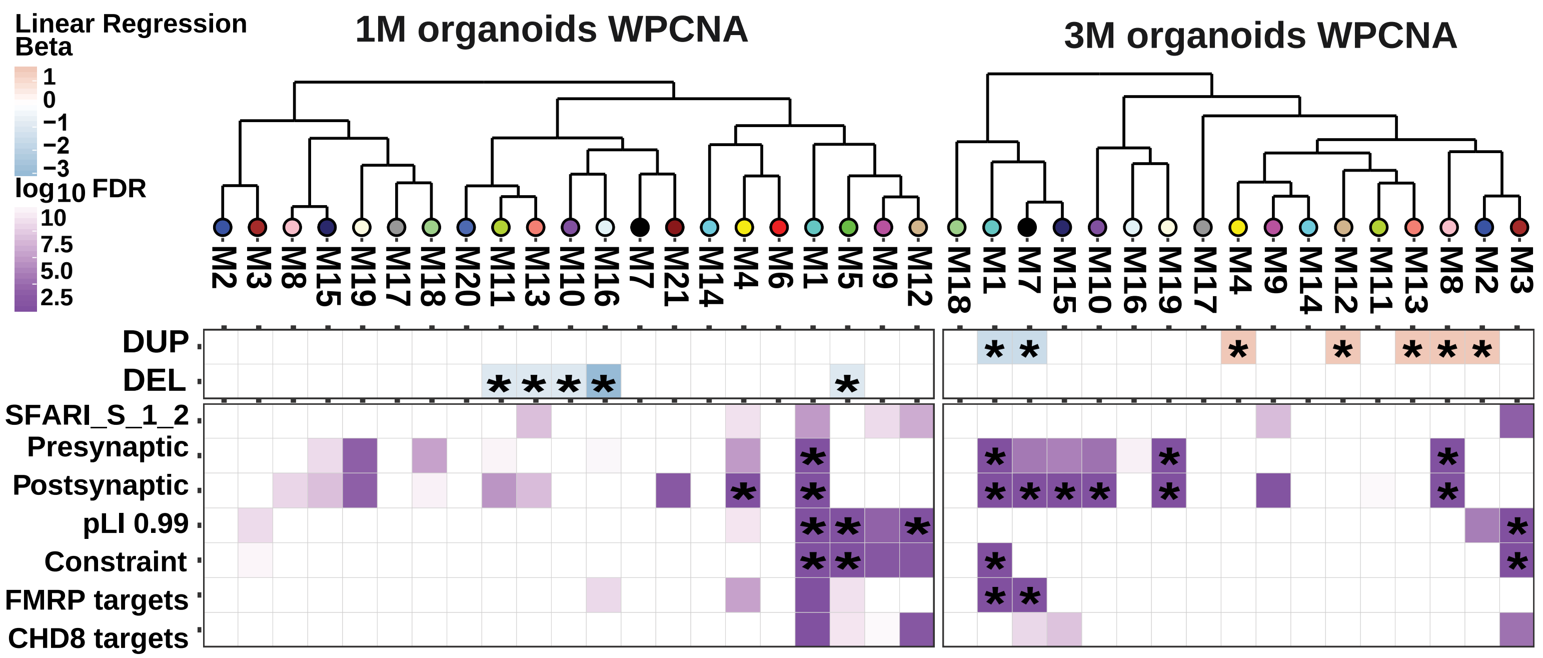

### Supplementary Fig. 17

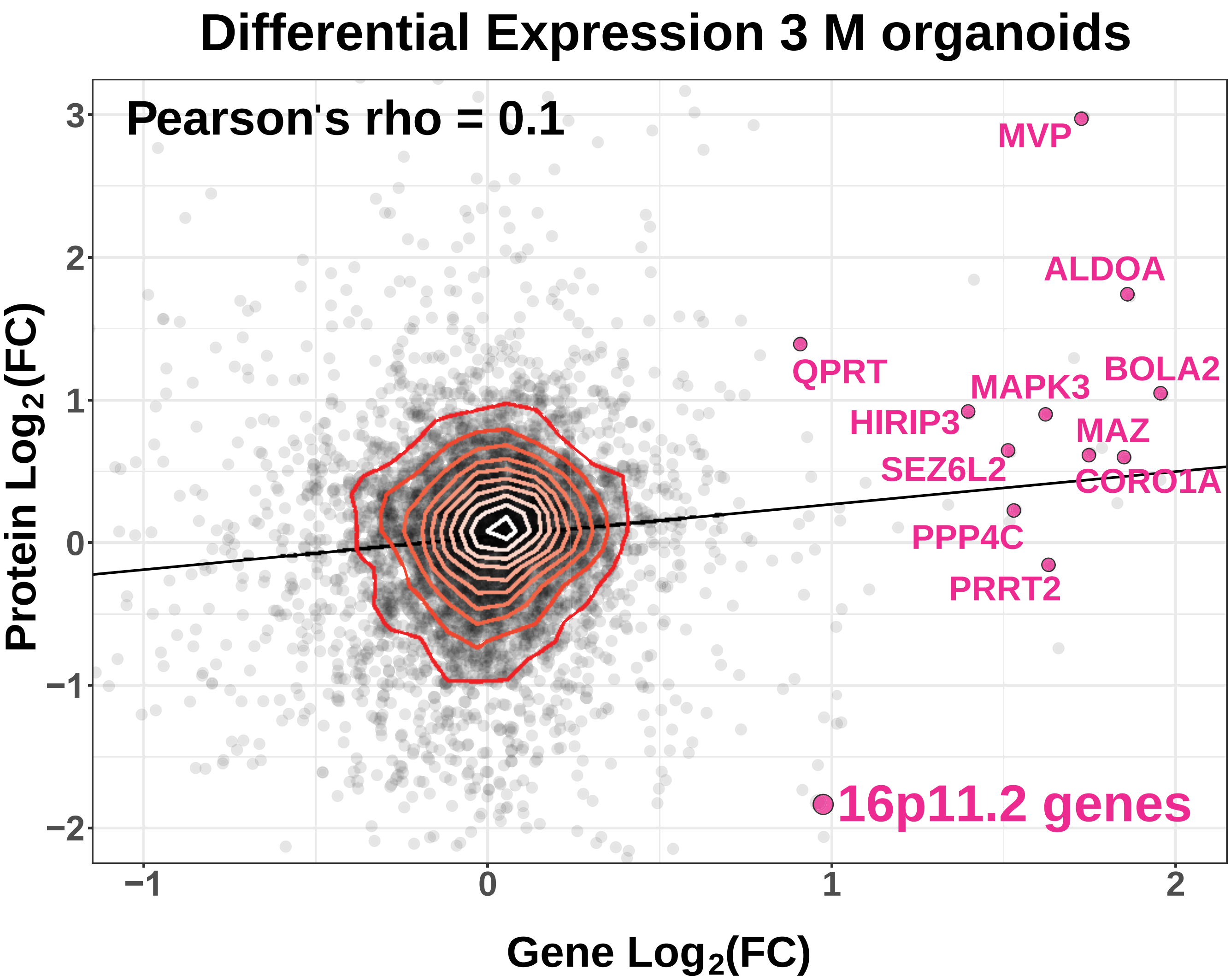

### Supplementary Fig. 18

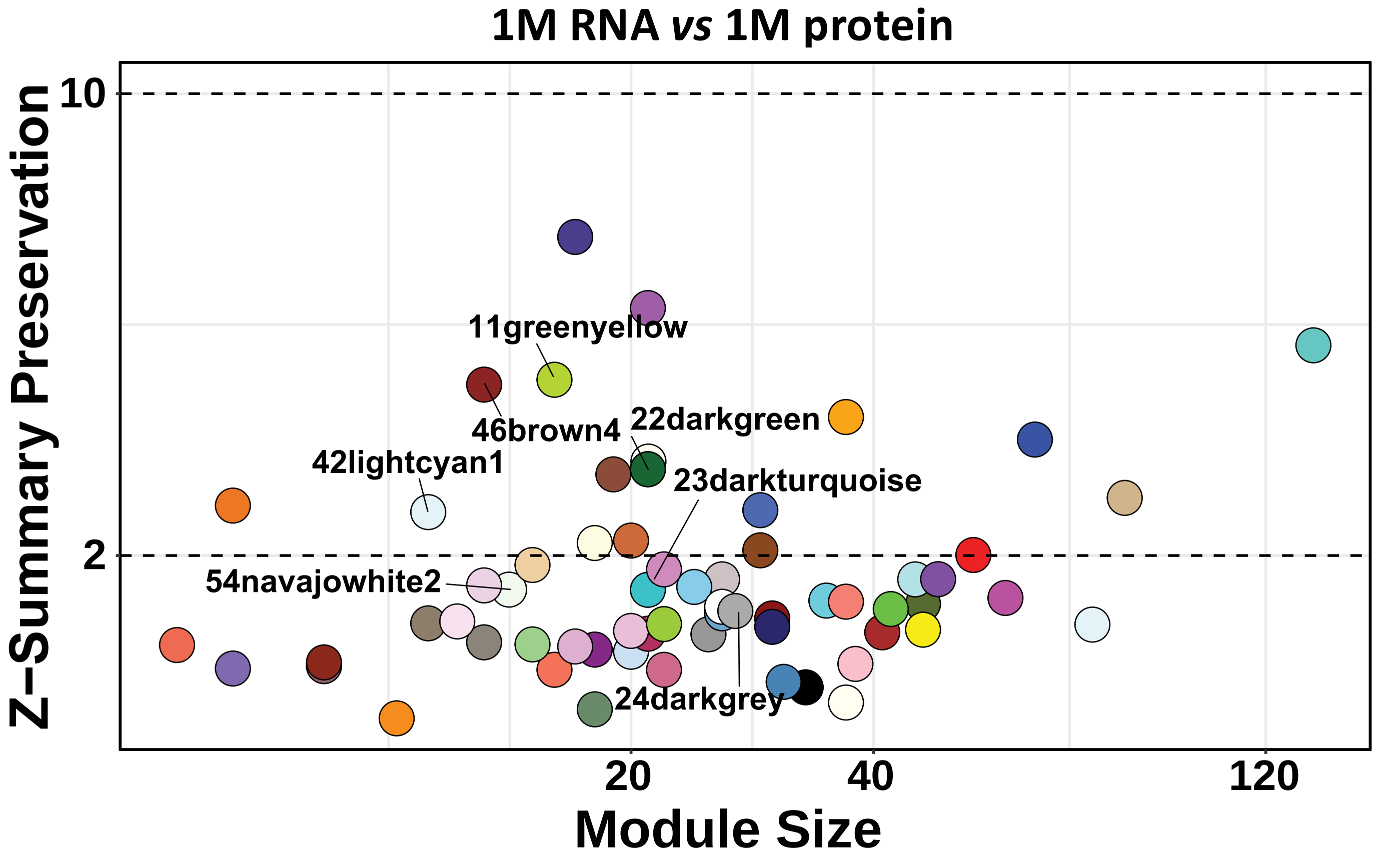


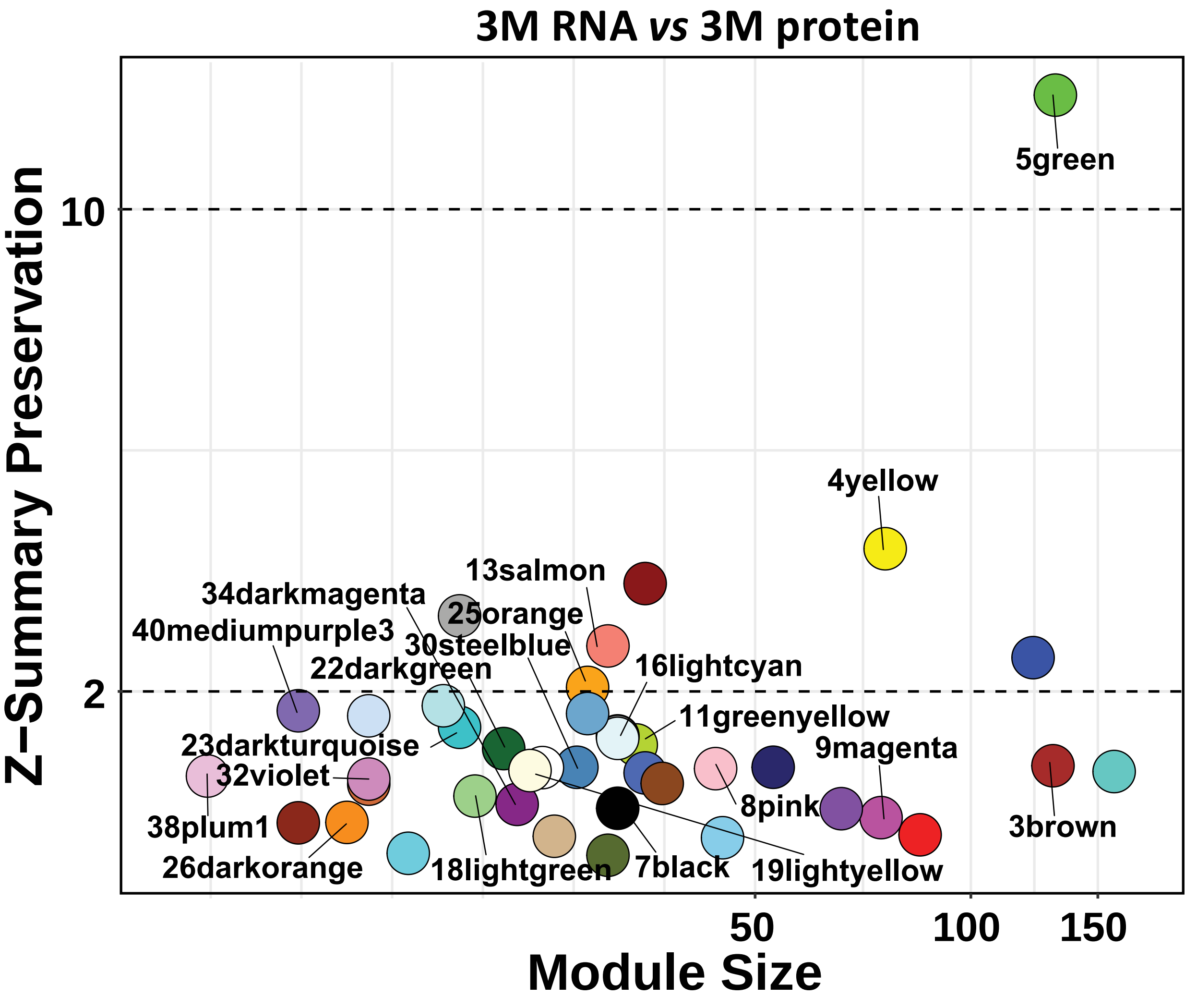

### Supplementary Fig. 19

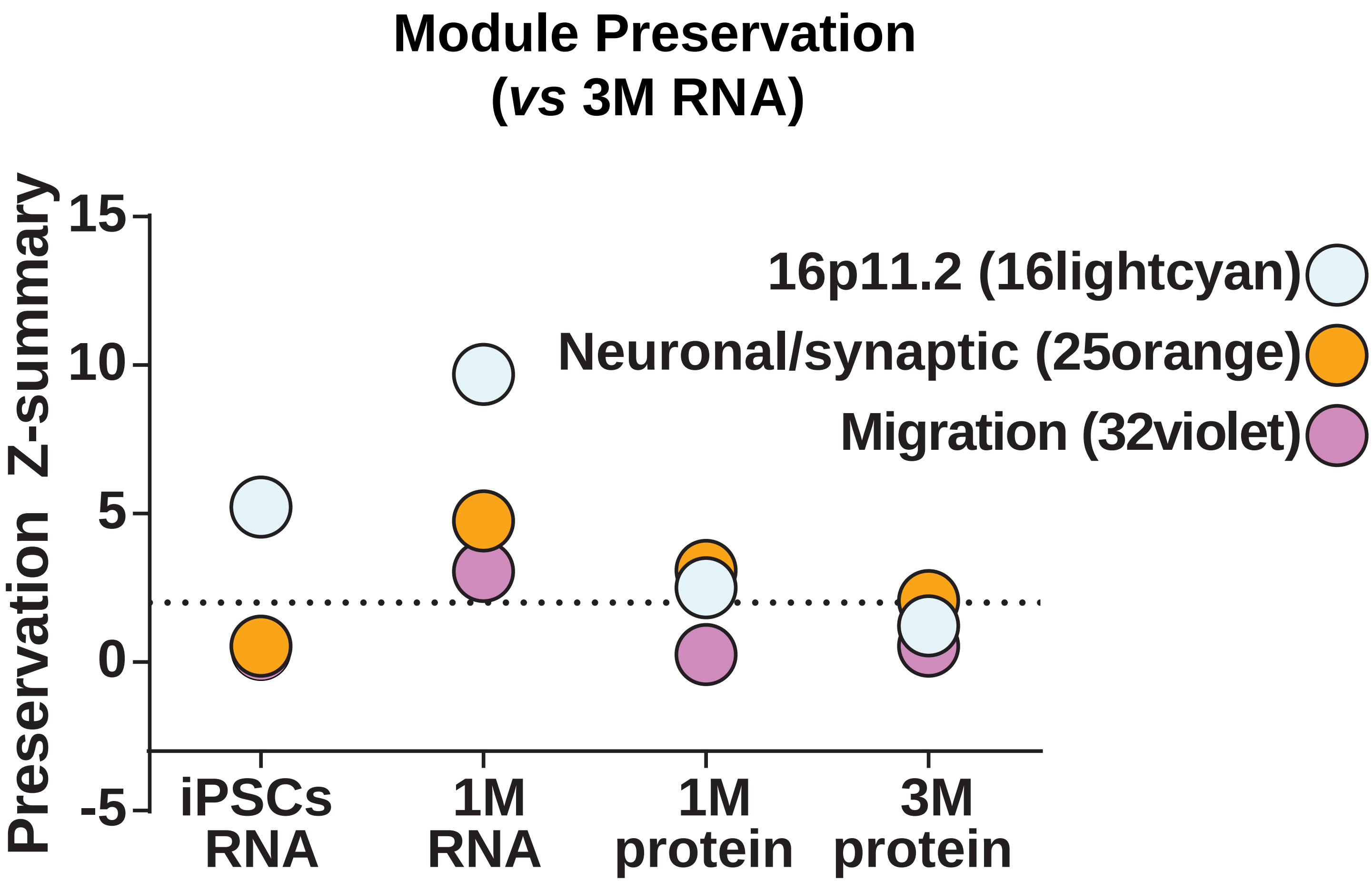

### Supplementary Fig. 20

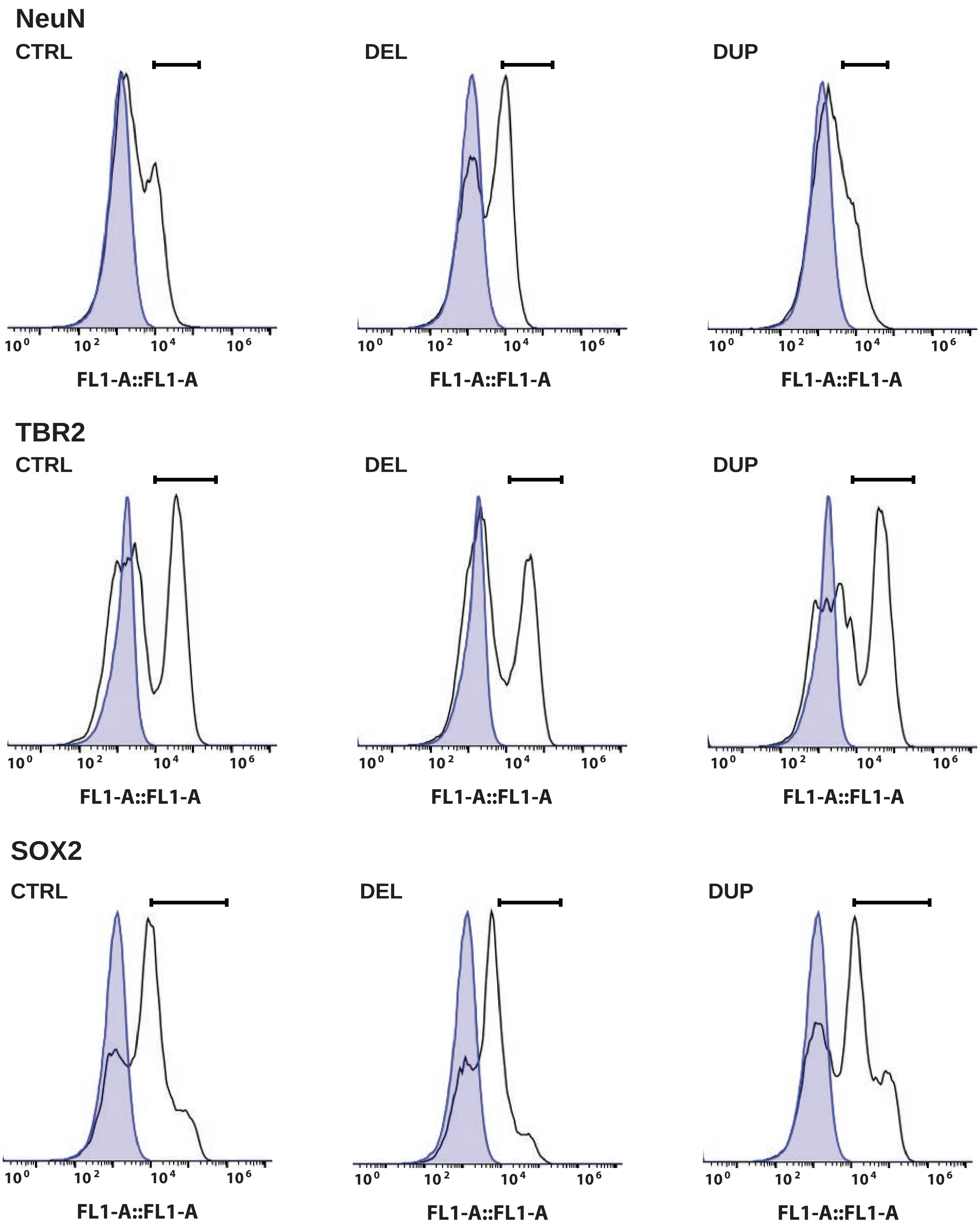

### Supplementary Fig. 21

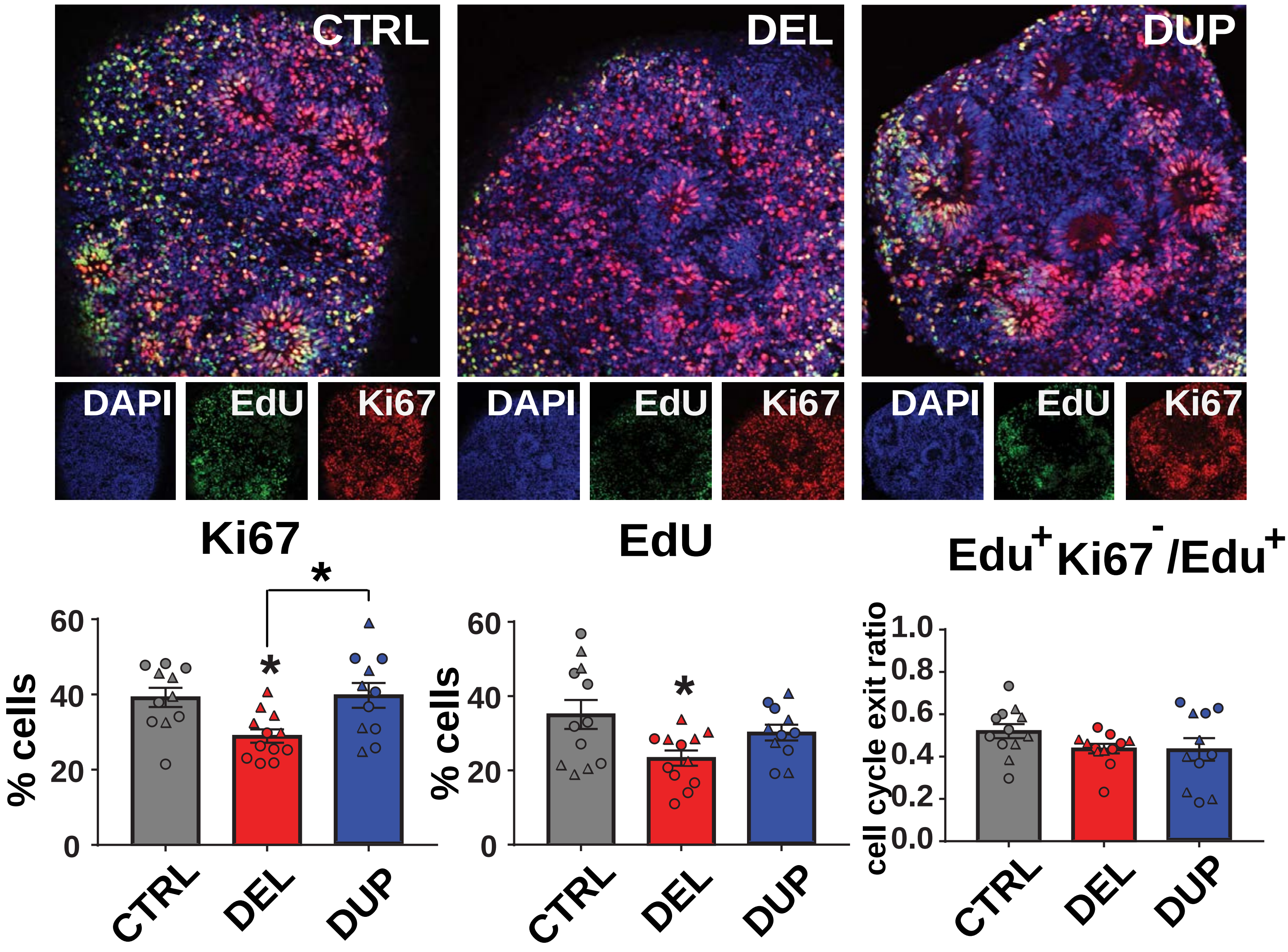

### Supplementary Fig. 22

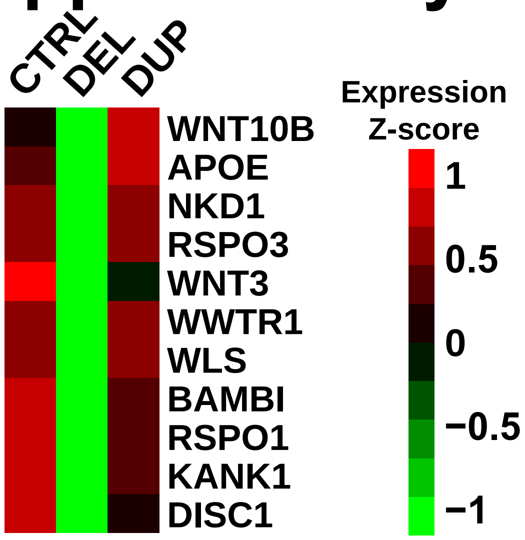

### Supplementary Fig. 23

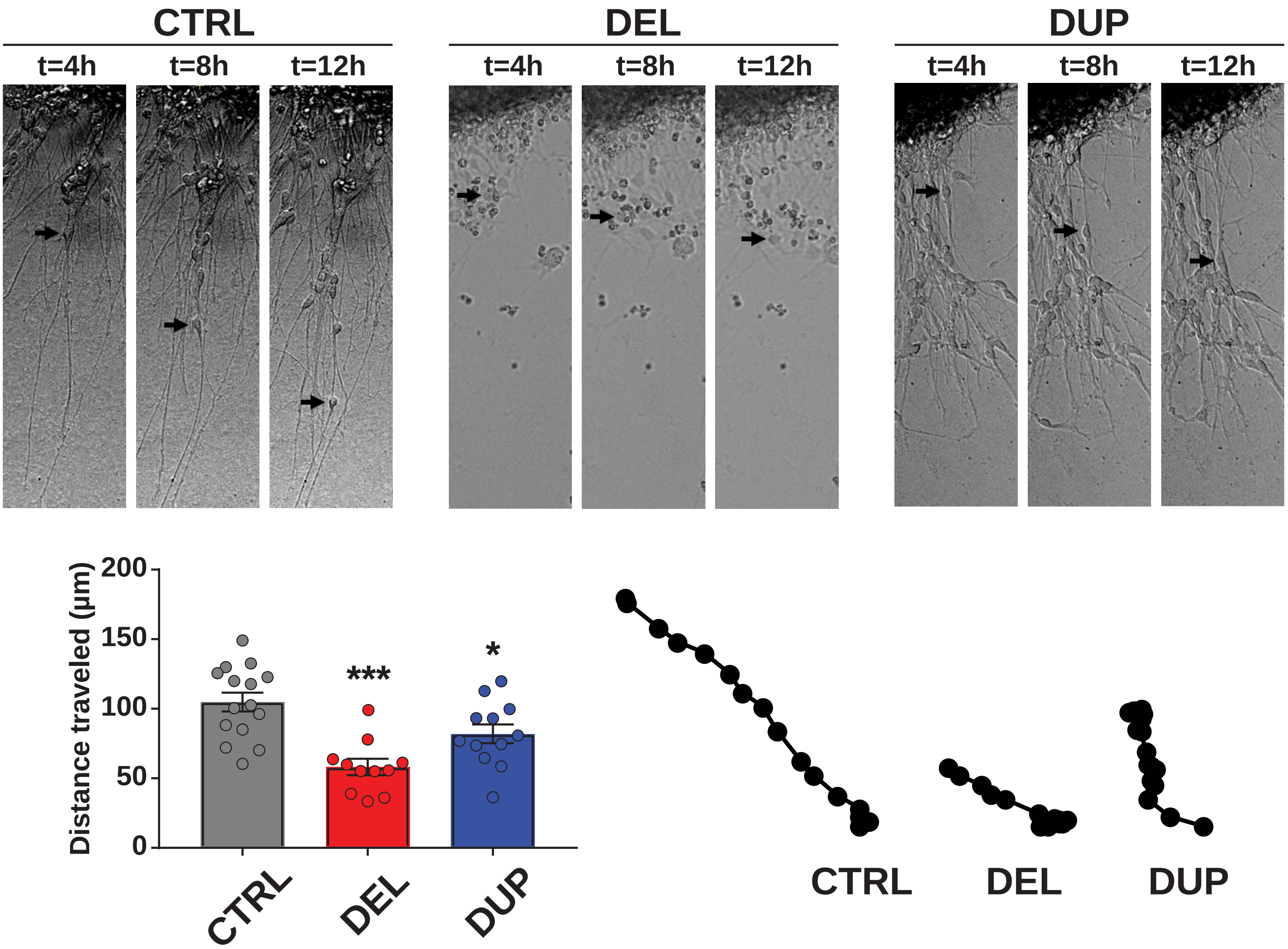

### Supplementary Fig. 24

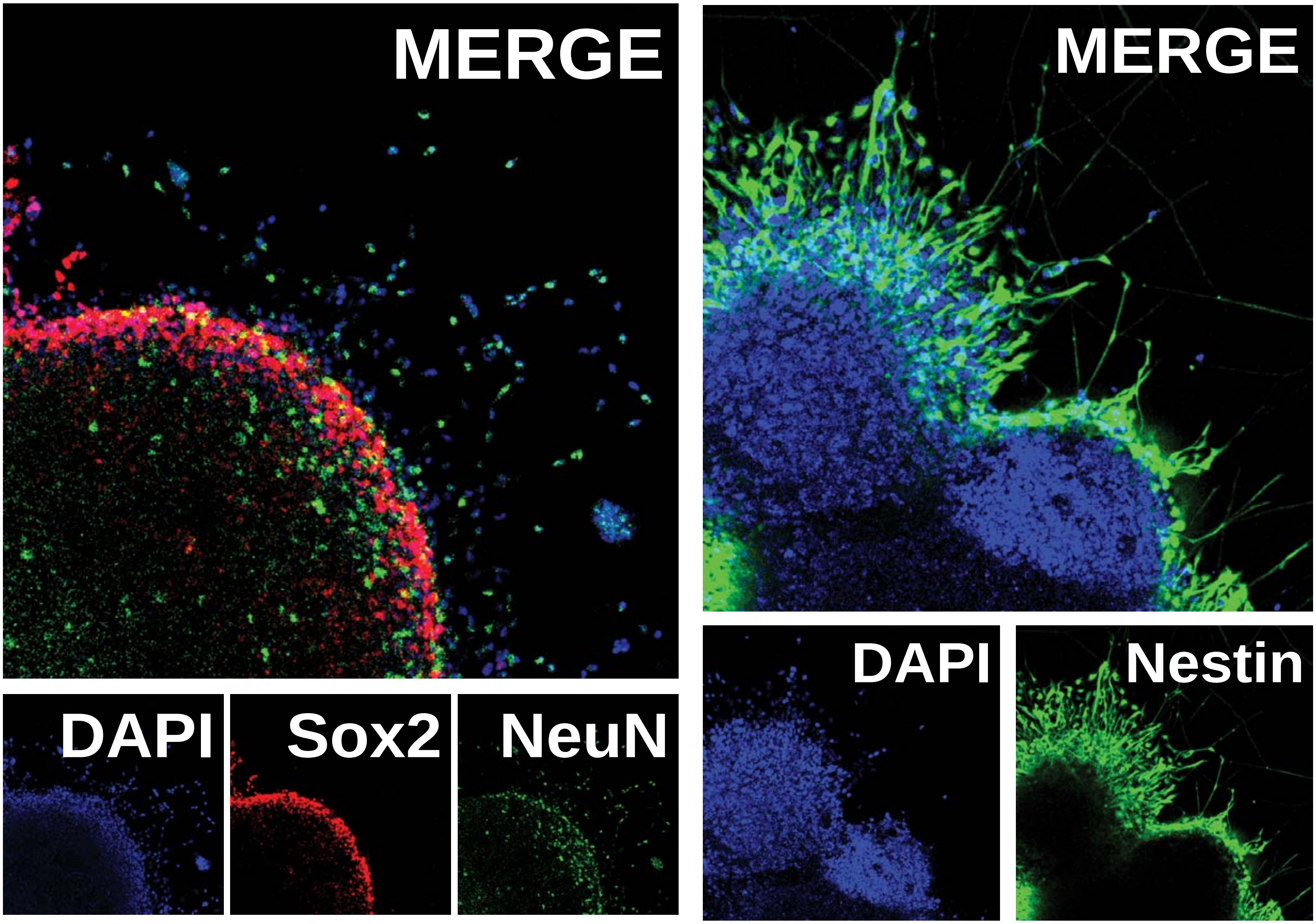

Extended Data Fig. 25

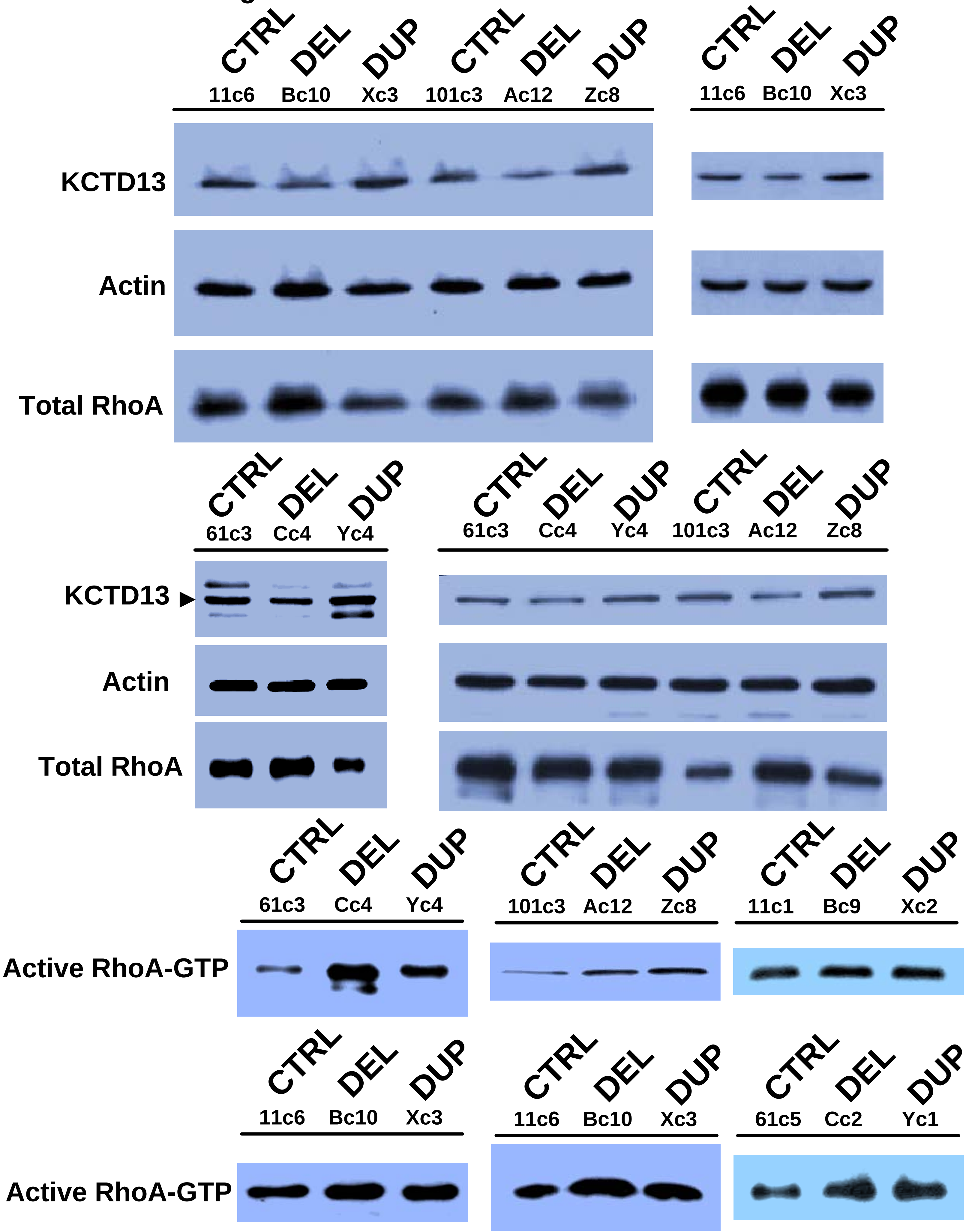

### Supplementary Fig. 26

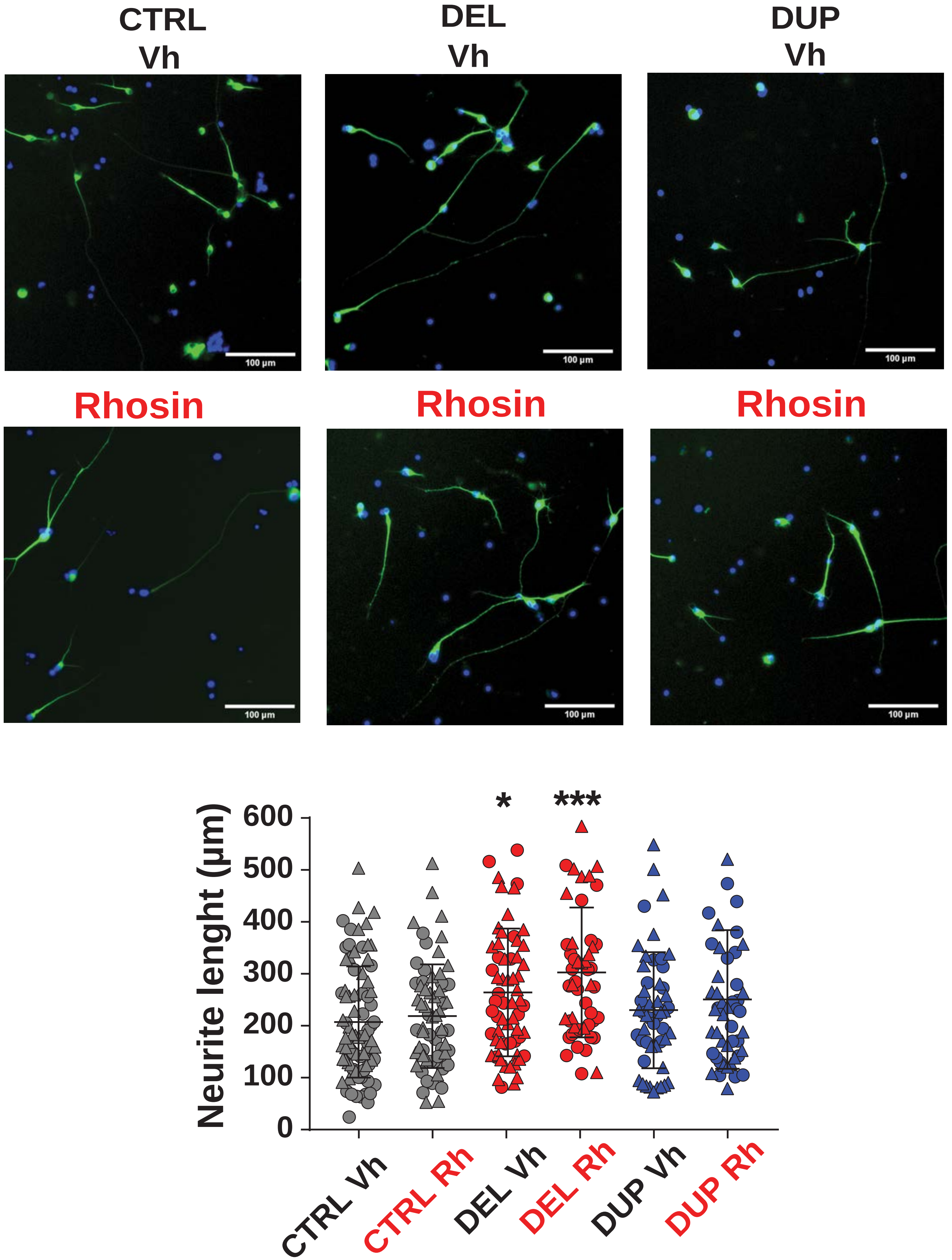

### Supplementary Fig. 27

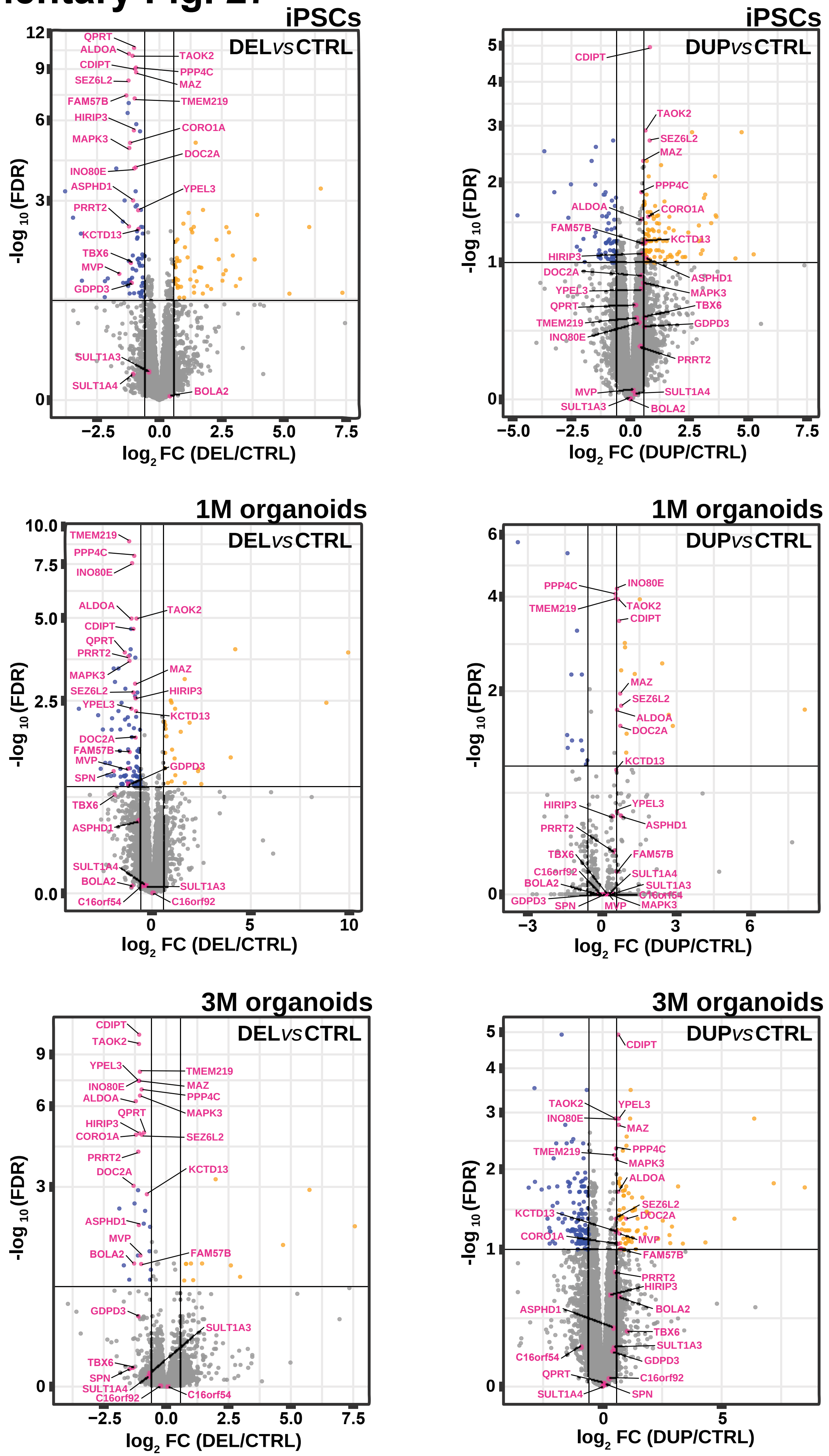

Supplementary Fig. 28

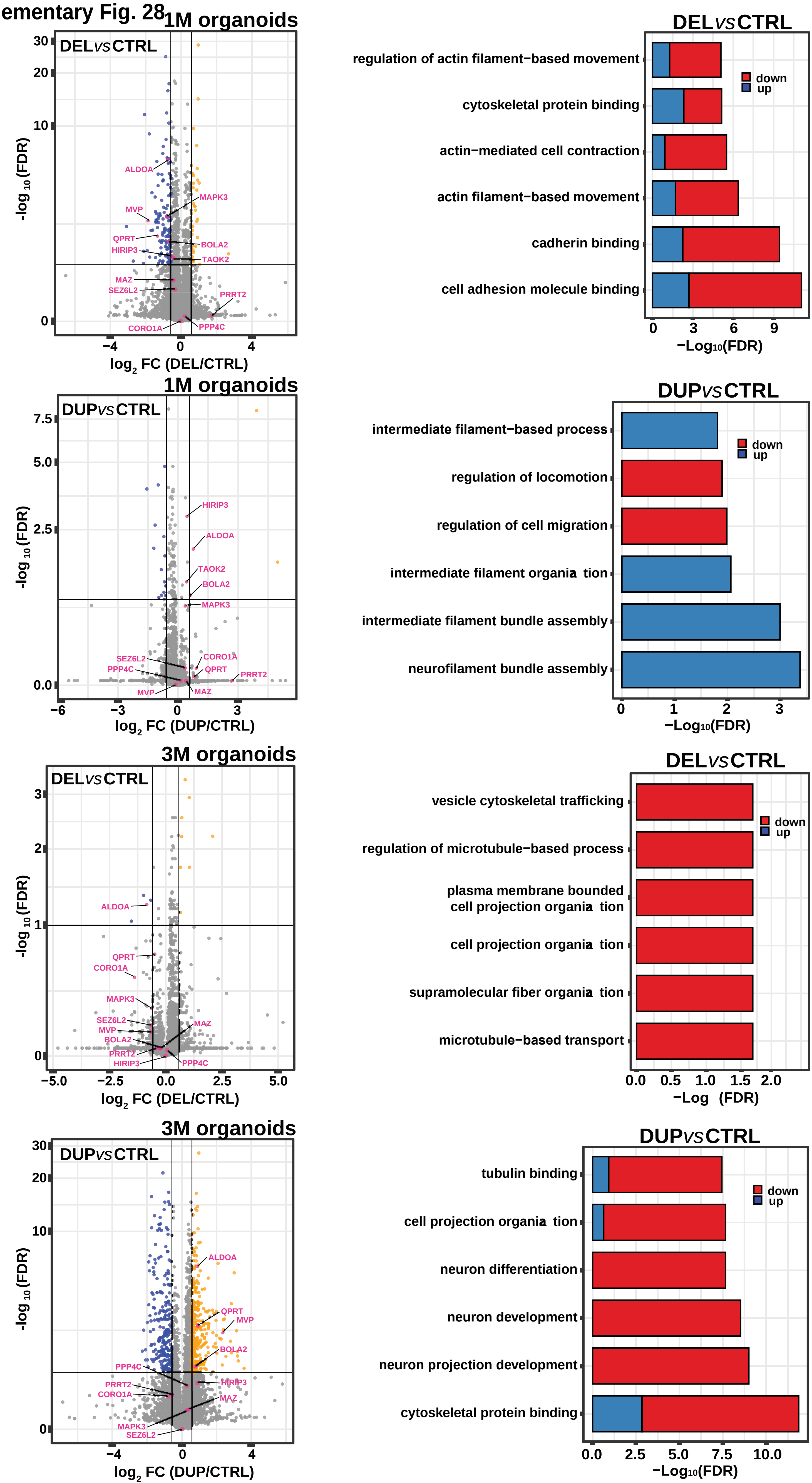
